# Curing “GFP-itis” in Bacteria with Base Editors: Development of a Genome Editing Science Program Implemented with High School Biology Students

**DOI:** 10.1101/2023.02.06.527367

**Authors:** Carlos A. Vasquez, Mallory Evanoff, Brodie L. Ranzau, Sifeng Gu, Emma Deters, Alexis C. Komor

## Abstract

The flexibility and precision of CRISPR-Cas9 and related technologies have made these genome editing tools increasingly popular in agriculture, medicine, and basic science research over the past decade. Genome editing will continue to be relevant and utilized across diverse scientific fields in the future. Given this, students should be introduced to genome editing technologies and encouraged to consider their ethical implications early on in pre-college biology curricula. Furthermore, instruction on this topic presents an opportunity to create partnerships between researchers and educators at the K-12 levels that can strengthen student engagement in science, technology, engineering, and mathematics (STEM). To this end, we present a three-day student-centered learning program to introduce high school students to genome editing technologies through a hands-on base editing experiment in *E. coli*, accompanied by a relevant background lecture and facilitated ethics discussion. This unique partnership aims to educate students and provides a framework for research institutions to implement genome editing outreach programs at local high schools.

## Introduction

Genome engineering (or genome editing) is the manipulation of the genomic sequence of a living organism through the addition, deletion, correction, or replacement of DNA in a precise, efficient, and controllable manner. Due to the prevalent and ever-evolving nature of genome editing technologies and their impact on society, introducing these tools to students in high school curricula is becoming critical (1,2). Further, given the application of genome editing technologies in controversial areas such as germline editing and gene drives, educating today’s students about these subjects prepares them to advocate for the appropriate use of these tools in the future (3–9).

In the United States, institutions of higher education have a duty to engage with, inspire, and train the next generation of STEM students at the K-12 public school level (10,11). Several scholars have urged higher education institutions to reach out to these classrooms to remove the wall between academia and their community schools (12). Data have shown that limited interactions between university professors and public-school educators can negatively affect students’ ability to transition to universities successfully (10). Research groups can address this issue by initiating research-practice partnerships with their local communities, in which they partner with local educators and design or implement student-centered learning programs that emphasize connections between innovative research and real-world applications (13). By focusing on hands-on STEM activities and exposing students to STEM careers, universities can contribute to developing the next generation of STEM professionals while simultaneously reducing educational disparities by broadening access to higher education (14–17).

To address this, we designed the Genome Editing Technologies Program and partnered with a local public high school to implement the program and evaluate its effectiveness. This program centers around a time- and resource-effective hands-on base editing laboratory experiment, with an intended audience of junior and senior high school students who have completed introductory biology coursework. While educating students on base editing technologies was the primary goal, we also aimed to leverage students’ exposure to scientists from diverse backgrounds and invited questions about personal experiences in school, professional development, and the day-to-day life of a graduate student or faculty member within academia (18,19).

The wild-type CRISPR-Cas9 system is commonly featured in science news outlet stories and introduced to pre-college students (1). In fact, several high school-accessible experiments that use CRISPR-Cas9 to “cut” DNA have been developed (such as the Out of the Blue CRISPR Kit; Bio-Rad 17006081EDU) (20). However, pre-college students are less aware of newer genome engineering tools like base editors (BEs). BEs enable scientists to introduce point mutations at targeted sites in the genome of living cells with high efficiency and precision (21,22) and thus have the therapeutic potential to treat thousands of human genetic disorders (**Figure 1A-B**) (23–25). A hands-on laboratory-based experiment that uses base editing would introduce students to cutting-edge genome editing technologies while building upon previous knowledge of CRISPR technologies (as BEs are modified CRISPR systems). In this manner, even students with prior knowledge of genome editing are engaged, and all students can connect what they learn to examples of CRISPR in the media.

**Figure 1.**
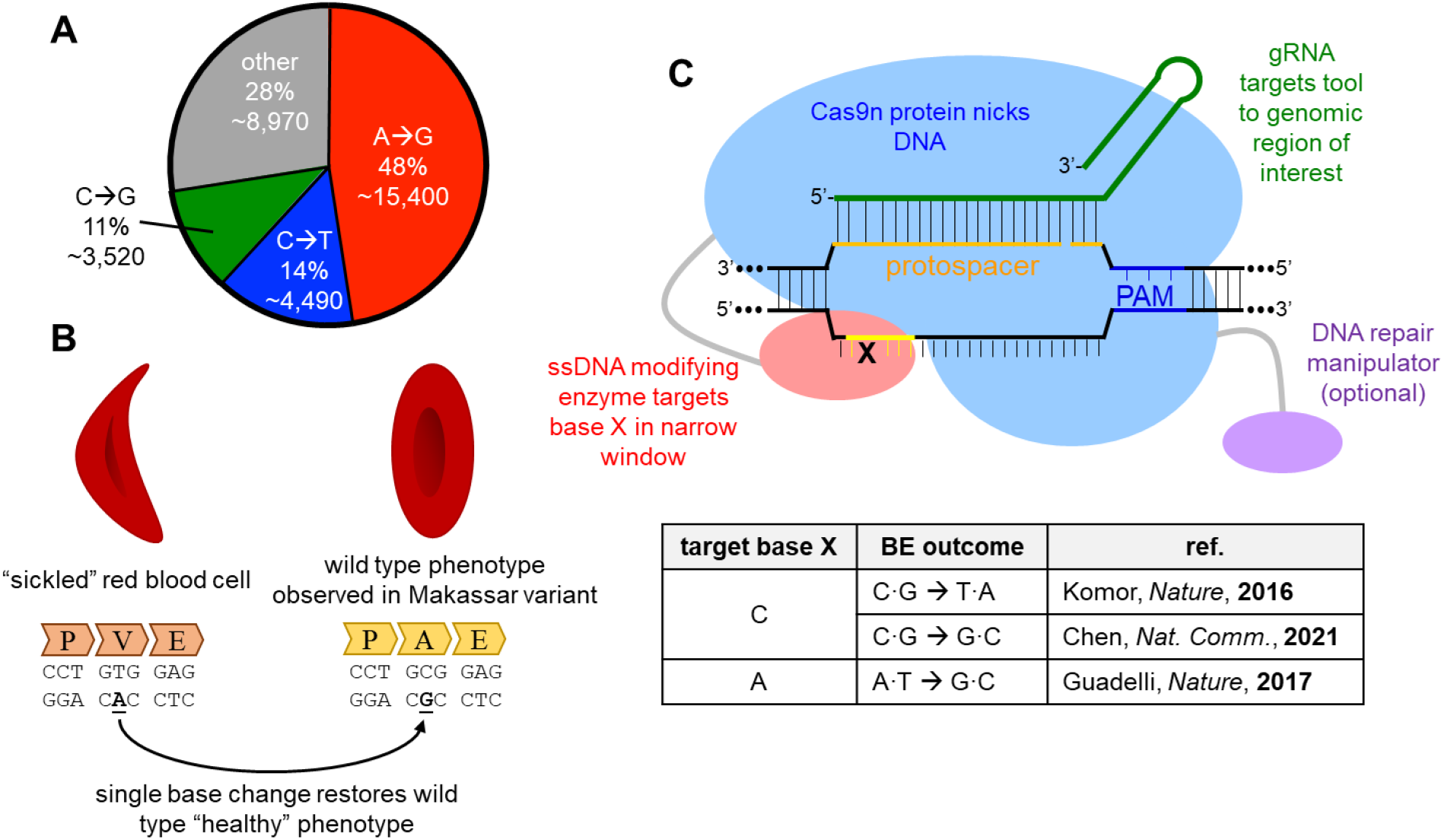
**A.** Current base editor technologies have the capacity to correct a large fraction of human pathogenic single nucleotide variants back to wild-type. **B.** BEs can also be used therapeutically to treat genetic disorders beyond simply correcting mutations back to wild-type. This includes the therapeutic example highlighted in this activity in which an ABE is used to modify the sickle cell anemia-causing mutation in the *HBB* gene. While the resulting A•T to G•C point mutation does not correct the gene back to wild-type, it does result in a phenotypically healthy variant. **C.** Current base editors use a common architecture to achieve single base conversions. Cas9n (blue) in combination with a targeting gRNA (green) directs the editor to the genomic site of interest by base-pairing with the protospacer (orange) sequence. A protospacer adjacent motif (PAM, blue) is also required for Cas9:DNA binding. This then allows the ssDNA modifying enzyme (red) to chemically modify a target base of interest within a small window of exposed ssDNA (yellow). Overall base pair conversions (listed in the table at the bottom) are determined by the nature of the ssDNA enzyme, and the inclusion of DNA repair manipulation components (purple).

The CRISPR-based laboratory experiments that have been developed for high school students use Cas9 to install double-stranded DNA breaks (DSBs) in genes of interest in *Escherichia coli* (*E. coli*). However, DSB-reliant genome editing methods are typically limited in their therapeutic potential, as precision editing outcomes are generally inefficient and come with the concurrent introduction of undesirable byproducts when using DSBs to perform genome editing in mammalian cells. BEs avoid this problem by linking a DNA-modifying enzyme to Cas9n, a partially inactive version of Cas9 that retains the ability to bind to DNA in a programmable manner but nicks rather than breaks the DNA backbone (**Figure 1C**) (26). BEs unwind and bind to a target DNA sequence in the genome (programmed by the sequence of their guide RNA, gRNA), and chemically modify DNA nucleobases within a small “editing window”. Cytosine base editors (CBEs) make targeted C•G to T•A point mutations, while adenine base editors (ABE) make A•T to G•C point mutations within living cells (**Figure 1C**).

Since their creation in 2016, BEs have been optimized and widely applied (27). Proof-of-concept studies have already demonstrated their potential in cell therapies and for treating progeria, sickle cell disease, and liver diseases (28–34). Given this and the ever-expanding use of these technologies, we believe that as researchers who develop these tools, we have a responsibility to educate the next generation of scientists about their function, use, and applications.

## Developing a Base Editing Activity Utilizing Fluorescence

While there are Cas9-based activities for high schoolers (20), to our knowledge, no one has introduced base editing to this demographic through a hands-on experiment. We therefore sought to develop a base editing experiment that high school students could accomplish in a single class period and with a minimal set of resources that might be accessible to a public high school laboratory classroom. With these restrictions in mind, we generated a base editing reporter system for use in *E. coli.* In this system, base editing activity results in expression of the green fluorescent protein (GFP). GFP-expressing colonies fluoresce green under blue light and can easily be seen by eye using a commercially available blue flashlight (**Figure 2**) (35). To enable a direct comparison with sickle cell anemia (36), a genetic disorder caused by a single point mutation that inactivates a crucial protein (**Figure 1B**), we installed a G•C to A•T mutation (Ala111Val) in the *GFP* gene to abolish its fluorescence (termed “dead” GFP or dGFP; **Figure 2**) (37). Targeting this mutation with an ABE and a properly designed gRNA corrects the mutation back to wild-type with an A•T to G•C edit, resulting in bacterial cells that fluoresce green. To emphasize the connection to genetic diseases, we call this phenotype “GFP-itis”, and students are therefore tasked with “curing” bacteria containing the *dGFP* reporter plasmid (“GFP-itis pSel”, addgene: 195344) by treating them with an ABE.

**Figure 2.**
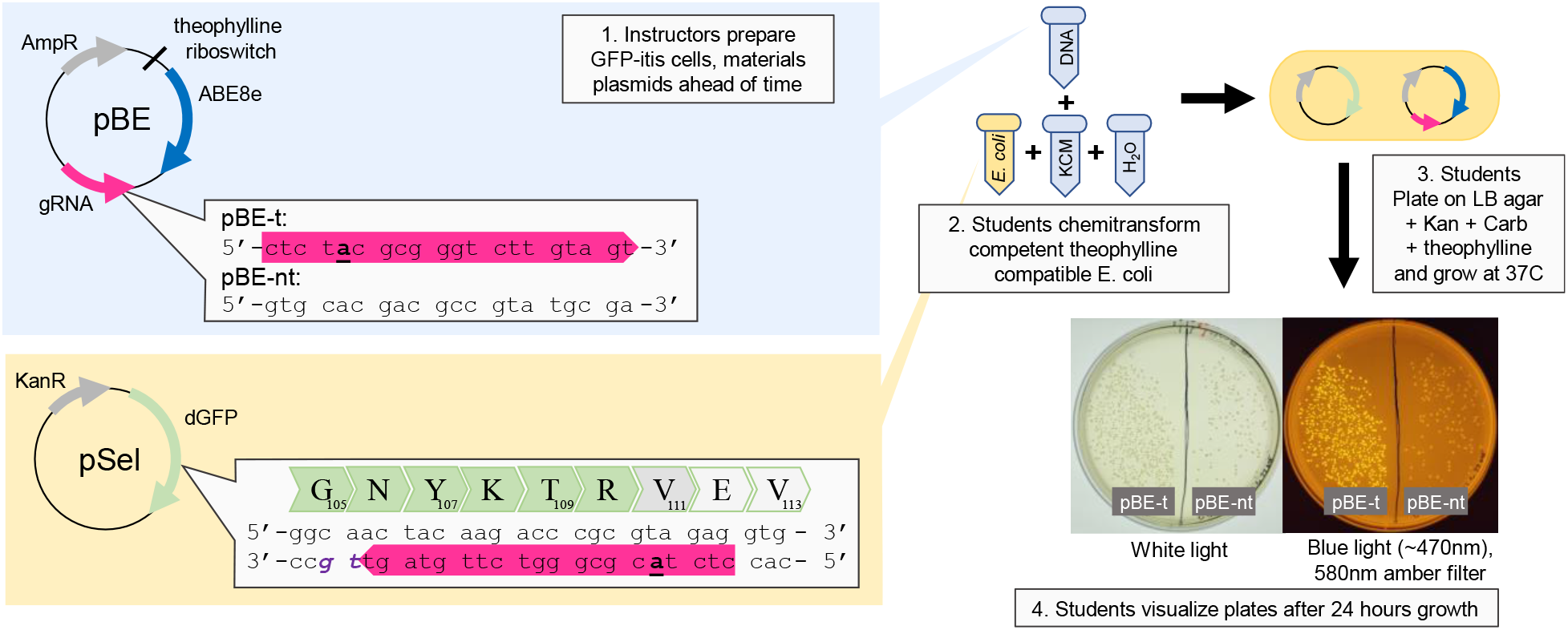
Illustration of constructs used in the “GFP-itis”activity. The inactivated *GFP* (*dGFP*) gene is in the pSel construct (bottom left, yellow background). Shown is the sequence of the *dGFP* gene, zoomed in on the inactivating A111V mutation that is corrected by editing of the target **<u>a</u>** to g. The pBE plasmids (top left, blue background) contain the ABE8e editor under control of a theophylline-responsive riboswitch, and one of two gRNA sequences. In pBE-t, the gRNA matches the pSel dGFP mutation site (pink arrow) and will lead to correction of the *GFP* gene and green fluorescence. In pBE-nt, the gRNA has a non-targeting sequence and acts as a negative control. Instructors prepare all plasmids and, prior to student transformation, incorporate pSel into *E. coli* to create “GFP-itis cells”. Base editing activity (GFP fluorescence) can be visualized 24 hours post-transformation (shown on the right).

Students deliver the ABE- and gRNA-encoding plasmid (pBE, **Figure 2**) into bacteria via chemical transformation (38). This process requires minimal equipment, is unaffected by imprecise volume measurements and timings that may result from differences in classroom materials, and is suitable for the skill level of high school biology students. The ABE used is the ABE8e variant, an editor evolved for fast kinetics and high activity levels (39). Because constitutively expressed BEs have shown higher toxicity levels in *E. coli* (40), our system uses a theophylline-responsive riboswitch to limit editing activity to the period in which the bacteria are plated on theophylline-containing agar plates (41). The ABE-containing construct also includes a gRNA cassette. Two variations of the pBE plasmid are available: “GFP-itis pBE-t” (addgene: 195342), which contains a targeting gRNA that targets the ABE to the *dGFP* point mutation, and “GFP-itis pBE-nt” (addgene: 195343), which includes a nonsense gRNA that acts as a negative control (**Figure 2**). The two plasmids (pBE and pSel)were designed with unique maintenance antibiotic resistance genes to ensure the retention of both plasmids and control for environmental bacterial contamination when plated on corresponding antibiotic-containing agar plates.

The full protocol is listed in the Supplemental Information, and is split into two parts. The first is preparatory work to be done by the research practitioners with access to standard microbiological laboratory equipment. The second details the practical activity to be undertaken by students and can be done in a classroom with access to a water bath, an incubator, and a set of pipettes. The protocol also contains a detailed list of materials, chemicals, and equipment (required and recommended). We list recommended portable equipment in the SI if the classroom does not have access to certain equipment.

## Implementing the Genome Editing Technologies Program

Our goal was not only to make base editing accessible to high school students but also to have students think critically and reflect on base editing in a social and cultural context. We developed a three-day program that centered around the following activities:

- Day 1: An interactive lecture on genome editing technologies (Time: 50-90 minutes)
- Day 2: A hands-on base editing experiment and discussion of ethics (Time: 50-90 minutes)
- Day 3: Reviewing experimental results and an open forum panel discussion (Time: 50-90 minutes)
  - Activity lengths can be adjusted according to the high school’s classroom schedules.

To implement the Genome Editing Technologies Program, we reached out to a local public high school and worked with three intermediate-level biology classes consisting of 20-25 students each. In total, 61 high school seniors enrolled in the Project Lead the Way biomedical sciences curriculum (42,43) participated in the Genome Editing Technologies Program.

## Day 1: Genome Editing Interactive Lecture

The first day introduces the students to genome editing technologies through an interactive lecture. The lecture focuses on teaching students about genome editing applications, the mechanics of CRISPR-Cas9 and base editing technologies, and the premise of Day 2’s experiment and its relationship to current therapeutic strategies. The learning outcomes prioritized in this activity are:

1. Describe how genome editing technologies are applied in medicine, agriculture, and basic science research.
2. Identify the components of base editors and explain how they work as a genome-editing tool.
3. Explain how a base editor can make a DNA mutation (within the context of Day 2’s activity).

During the lecture session, we emphasized active learning exercises to engage students in learning (44). Specifically, we used Mentimeter (45) to allow students to anonymously give their responses to questions such as “what are words that come to mind when you hear the term ‘Genome Editing’?” (Supplemental Figure 1). The lecture also includes several real-world applications of genome editing, such as CRISPR-modified anti-browning mushrooms (46) and the current state of a therapeutic application of genome editing for a patient with Hunter’s syndrome (47). We then facilitated an informal discussion with the students after presenting our examples by asking open-ended questions such as, “can you think of additional applications of genome editing?”. Through these discussions, students seemed most interested in medical applications of genome editing, further demonstrating the importance of using base editing as a hands-on example rather than DSB-reliant tools.

The lecture material also explains the mechanics of BEs, emphasizing their potential to cure “GFP-itis”. Specifically, this includes defining and illustrating the gRNA, Protospacer Adjacent Motif (PAM) sequence, Cas9n, and the deaminase enzyme components of base editing systems. Kahoot, a game-based digital platform, can administer review questions about BEs to help facilitate student learning (48,49). Students then form groups and complete a worksheet together to facilitate further distillation of the material. In the worksheet, students are asked to label the different components of the BE system, including writing in the spacer sequence of the gRNA to correct “GFP-itis” (based on the dGFP DNA sequence that is presented to them in the lecture slides). Students then identify the appropriate pBE plasmid (t or nt) to cure the “GFP-itis”. The worksheet also includes questions relevant to the investigation (such as

“which base is being targeted”, and “why are antibiotics needed on the plate”) to prompt discussions about the base editing experiment that the students will do the following day. The worksheet can also serve as comprehension feedback check for the instructors and guide any relevant remedial lessons. The lecture slides and the worksheet are included as Supplemental Material.

## Day 2: Base Editing Experiment & Discussion on Ethics

The second day consists of the hands-on laboratory experiment described previously. Students have the opportunity to perform base editing in *E. coli* using simple bacterial microbiology techniques and equipment present at the local high school. We had students form groups of 3-4, work together to complete the transformation, and plate the bacteria to visualize the next day. Students should be encouraged to record experimental observations.

The transformation recovery step requires a 10 to 60-minute incubation period, which we used to engage students in a conversation about the ethics of genome editing, emphasizing therapeutic examples to use our time efficiently. We presented different perspectives on defining a “genetic disease” (i.e., how do we differentiate between a “trait” and a “disease”) and encouraged them to think about how their genetics affect their personal identities. If time permits, we recommend screening portions of the documentary “Human Nature”, produced by The Wonder Collaborative (50). We also broached the subject of germline genome editing, in which edits are inherited by all future descendants of the edited individual, regardless of whether these future descendants consent to the procedure (3–9,51). We asked students to consider the risks involved in germline editing and the issues surrounding medical consent in hypothetical cases of germline editing. These examples introduced students to alternative perspectives about genome editing therapeutics and demonstrated that these ethical dilemmas are not one-size-fits-all.

## Day 3: Experimental Results and an Open Forum Panel Discussion

The students begin the third day by evaluating the success of their attempts to cure “GFP-itis” with base editing by visualizing their plates under blue light. We invited students to think critically about their results and identify shortcomings of the “GFP-itis” treatment by referring to their experimental observations from the day before. Some groups observed fundamental issues, such as very few (or sometimes no) colonies, and could usually connect this back to a technical issue during their transformation. Other groups observed very few colonies with GFP fluorescence, presenting an excellent opportunity to discuss the limitations of these tools in terms of scale and efficiency. Students completed written feedback surveys on the program as we worked through the groups to visualize the experiment. We briefly discuss these results below, and used them to improve upon our worksheet and lecture slides.

After visualizing their results, we had an open forum panel where we encouraged the students to ask us questions about current genome editing research, ethical issues, and professional development. This element is a crucial component of the program, as the students built upon the connections fostered with us through the previous days to ask questions and seek advice about college and graduate school during the panel. Feedback from the high school instructor (included as Supplemental Material) was overwhelmingly positive and strongly advocated for the program in the future. Our experiences implementing the Genome Editing Technologies Program strongly support the conclusions of previous studies that research-practice partnership such as this strengthen ties between academia and their communities and provide opportunities for students to nourish their interests in STEM (10,13,16).

## Discussion on Student Experiences and Evaluation of the Program

The results from our feedback survey are shown in **Figure 3**. We first asked the students to evaluate the accessibility of the various components of the program. The students indicated that most of the components were accessible (for example, 87% of the students [n=60] indicated that the lecture was accessible, and 85% of the students [n=60] indicated that the ethics discussion was accessible **Figure 3A**). However, student-evaluated accessibility of the worksheet was only 43%. We have since improved the worksheet using the students’ feedback by further clarifying some questions and updating instructions for labelling diagrams. The updated version is included as Supplemental Material.

**Figure 3.**
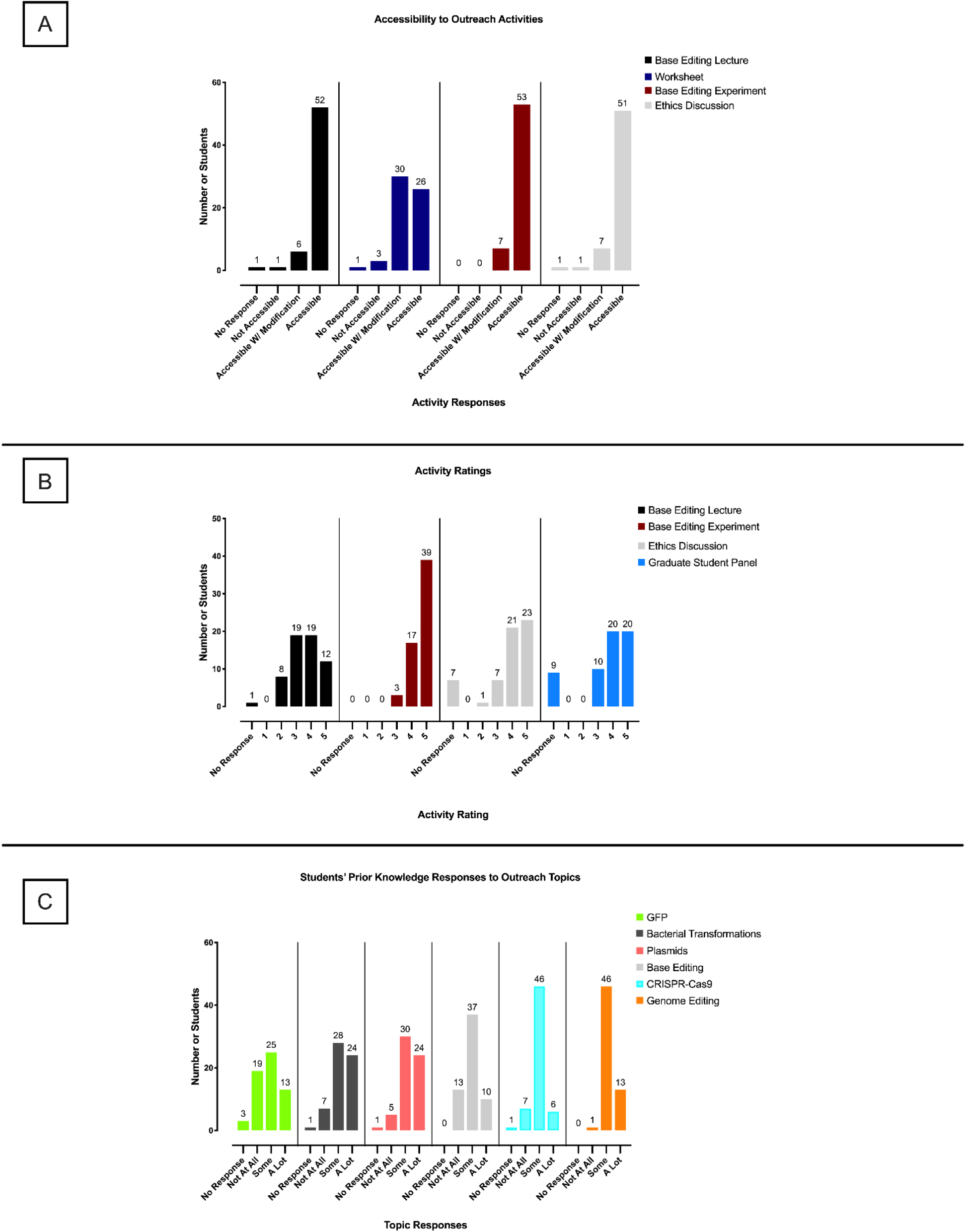
Student Survey Responses. (**A**) Accessibility rating for each of the four outreach activities: (from left to right) the base editing lecture (87% accessibility rating), the base editing worksheet (47% accessibility rating), the base editing experiment (88% accessibility rating), and the ethics discussion (85% accessibility rating). In total, there were 60 student responses. (**B**) Activity rating of each of the outreach activities. Answer options ranged from 1 being “not so much” to 5 being “it was great!”. Topics measured were (from left to right): the base editing lecture (3.60 activity rating average), the base editing experiment (4.61 activity rating average), the ethics activity (4.20 activity rating average), and the open forum discussion (4.27 activity rating average). In total, there were 59 student responses. (**C**) Measurements of the students’ prior knowledge to various topics covered in the program. Answer options include “no response,” “not at all,” “some,” and “a lot.” The topics queried were (from left to right): GFP, bacterial transformations, plasmids, base editing, CRISPR-Cas9, and genome editing. In total, there were 60 student responses.

We also evaluated the engagement of the program by asking the students to quantify how much they liked each component (ranging from a 1 being “not so much” to a 5 being “it was great!”, **Figure 3B**). We also included an “open comments” section for anonymous feedback. The lowest rating was observed for the lecture, which overall scored a 3.6 out of 5 (n=59), with 50 students indicating a 3 or higher. In the open comments section, several students commented favorably on the active learning elements of the lecture, prompting us to consider including more of these strategies in future designs. We have since incorporated more open discussion segments within the slides that allow students to reflect on the material, including a 7-question Kahoot quiz that is included as Supplemental Material.

Although the hands-on laboratory experiment was the most challenging element to prepare, it was the highest-rated activity in our program, with 88% of students (n=60) saying the activity was accessible (**Figure 3A**), and an average rating of 4.61 out of 5 (n=59; **Figure 3B**). The feedback results highlight the need for programs like ours; not only were these students introduced to tools and techniques that are used in many areas of biology via the lecture, but also researchers provided students the opportunity to use them in a hands-on laboratory experiment, which has been shown to increase student learning and retention (44,52,53).

## Conclusion and Future Outlook

We developed the Genome Editing Technologies Program to facilitate other academic researchers in the field of genome editing to implement research-practice partnerships with local public schools. Our program lasts three days, during which students are introduced to innovative genome editing techniques, partake in a hands-on base editing experiment, participate in ethics conversations, and are provided with opportunities for professional development. Notably, we include here requisite materials for others to reproduce our program with their local communities. To encourage the expansion of this program, we highlight the flexible nature of the three-day activity, which allows for modification depending on a research group’s time, resources, and expertise/interests. For example, a group might conduct the same activities but focus on some practical issues relevant to their specific research interests (delivery methods, off-target editing, or efficiency).

We additionally gauged prior knowledge of different topics that were discussed during the program by the students (**Figure 3C**). We were particularly excited to see that self-perceived prior knowledge of GFP and base editing were quite low (19 out of 60 students reported no prior knowledge of GFP, and 13 out of 60 students reported no prior knowledge of base editing), demonstrating the potential of our program to increase student knowledge of these topics. However, we did not assess this knowledge in a pre-post format. A more educational research-intensive group might choose to examine student learning more formally.

Furthermore, we provide a protocol that includes preparatory work to be carried out by educators without student participation. However, students with additional resources and time might find value in preparing their materials. To adapt this program for a more advanced class, such as an undergraduate laboratory course unit, instructors could emphasize the role of designing genome editing tools and ask students to generate their own gRNAs to correct the dGFP sequence. We invite researchers to use our program as a platform to build future outreach activities and incorporate innovative technology into early public education curricula.

## Supporting information

Supplemental Material - Instructor Feedback

Supplemental Material - Lecture Slides

Supplemental Material - Worksheet

## Acknowledgements

We gratefully acknowledge the mentorship, guidance, and support from Dr. Thomas Bussey, Dr. Stacey Brydges (from the Department of Chemistry and Biochemistry at UC San Diego), and graduate student Monica Molgaard (in the Education Studies Program at UC San Diego). We also acknowledge support from lab members of the Komor Group, Zulfiqar Mohamedshah, Rachel Anderson, Sam Mawson, Michael Hollander, and Annika So. Thank you to Valerie Park and her students at Sage Creek High School.

## Author contributions

C.A.V. contributed to conceptualization of the research project, experimental design, data curation, data analysis, writing of the manuscript, and acquisition of the funding. M.E. contributed to conceptualization of the research project, experimental design, data curation, data analysis, and writing of the manuscript. B.L.R. contributed to experimental design, data curation, data analysis, and writing of the manuscript. S.G. contributed to experimental design. E.D. contributed to data curation and analysis. A.C.K. contributed to conceptualization of the research project, experimental design, data analysis, supervision of the work, writing of the manuscript, and acquisition of the funding. All authors contributed to editing of the manuscript.

## Data and material availability

Plasmids from this study are available at Addgene:

- Targeting base editor plasmid (pBE-t), Addgene: 195342
- Non-targeting control base editor plasmid (pBE-nt), Addgene: 195343
- Selection plasmid (pSel), Addgene: 195344

