## Supplemental Information for "Curing “GFP-itis” in Bacteria with Base Editors: Development of a Genome Editing Science Program Implemented with High School Biology Students"

**Protocol**

The described protocol has been split into two parts. The first is preparatory work to be completed by the instructors and requires access to standard microbiological laboratory equipment. The second half details the practical activity to be undertaken by students and can be done in a classroom with access to a water bath, an incubator, and a set of pipettes. If a high school classroom does not have all the requisite equipment, we offer suggestions of portable models that could be provided by the research practitioners in the "Additional Resources" column of the following table.

**Materials:**

| Material | Preparatory Need | Classroom Need | Additional Resources |
|---|---|---|---|
| | *Plasmids* | | |
| Targeting base editor plasmid (GFP-itis pBE-t) | X | X | Addgene #195342 |
| Non-targeting control base editor plasmid (GFP-itis pBE-nt), | X | X | Addgene #195343 |
| Selection plasmid (GFP-itis pSel) | X | | Addgene #195344 |
| | *Standard Lab Supplies* | | |
| 1.7mL centrifuge tubes | X | X | |
| Micropipette Tips: 10uL, 200uL, 1000uL | X | X | |
| Bacterial Spreader Loops | X | X | |
| Petri Dishes | X | | |
| Culture flasks: 5mL, 125mL | X | | |
| 50mL centrifuge tubes | X | | |
| | *Chemicals* | | |
| Carbenicillin or Ampicillin | X | | GoldBio C-103 |
| Kanamycin | X | | GoldBio - K-120-SL |
| Theophylline (CAS 58-55-9) | X | | ThermoFisher - 250310050 |
| Agar | X | | Sigma Aldrich - A5306 |
| TSS (10% w/v PEG 3350, 5% w/v DMSO, 20mM $MgCl_2$ in bacterial media) | X | | |

| | | | |
|---|---|---|---|
| Sterile Bacterial Media (standard LB or 2xYT recommended) | X | X | |
| Sterile DI water | X | X | |
| 5x KCM Buffer (0.5M KCl, 150mM $CaCl_2$, 0.25M $MgCl_2$) | X | X | |
| *Equipment* | | | |
| 37°C shaker | X | | |
| OD measuring spectrophotometer | X | | |
| Centrifuge | X | | |
| Micropipettes (p10, p200, p1000) | X | X | |
| 42°C water bath | X | X | |
| 37°C incubator | X | X | VEVOR 25 L capacity, 5 °C – 60 °C temperature range, 20.95-pound portable incubator |
| Blue light source or UVA light source (488 nm works best)* | | X | Invitrogen Dual LED Blue/White Light Transilluminator (LB0100) |
| Amber blue light filter glasses** | | X | |
| *Miscellaneous* | | | |
| 100uL aliquot of S1030 chemically competent bacterial cells*** | X | | Addgene #105063 |
| ice | X | X | |

*While a blue light source (488 nm) works best, UV can be used in its place. Note that the GFP in pSel is less visible with UV excitation. A UV flashlight with excitation 395nm can be substituted

** blue light filter glasses (ThermoFisher S37103) should be used if a handheld flashlight is chosen rather than an illuminator with built in screen.

*** Alternative strains of chemically competent bacterial cells must be compatible with the theophylline riboswitch

### Protocol:

*<u>Preparatory work (to be done by instructor in advance)</u>*:

1. Prepare 100mM theophylline stock solution in sterile DI water, for instructor use.

2. Prepare TSS solution (1) (10% w/v PEG 3350, 5% w/v DMSO, 20mM $MgCl_2$ in bacterial media) in sterile bacterial media, for instructor use.
3. Prepare 5x KCM buffer (0.5M KCl, 150mM $CaCl_2$, 0.25M $MgCl_2$) in sterile water, for student and instructor use (http://cshprotocols.cshlp.org/content/2008/9/pdb.rec11454.full).
4. Prepare agar plate containing 50ng/uL Kan for instructor use.
5. Prepare agar plates containing 50ng/uL Carb, 50ng/uL Kan and 1mM theophylline for student use. Note, each group of students needs 1 plate. Agar plates may be stored at 4°C until use.
6. Prepare chemically competent theophylline riboswitch compatible cells with “GFP-itis pSel” incorporated (“GFP-itis" cells), for student use:
    a. To 70uL sterile DI water, add 20uL 5x KCM solution and 1uL “GFP-itis pSel” (10-100ng plasmid). Rest on ice 5min.
    b. Add 100uL aliquot chemically competent bacterial cells that are theophylline riboswitch-compatible (S1030 Addgene #105063 recommended) to the plasmid solution. Return to ice 10min.
    c. Heat shock in 42°C water bath for 75 seconds. Return to ice for 2 min.
    d. Add 800uL sterile bacterial media for recovery.
    e. Shake to aerate and recover for 1hr at 37°C. Plate 20-50uL on agar containing 50ng/uL kanamycin maintenance antibiotic. Grow overnight in a 37°C incubator.
    f. The following day, pick a single colony and inoculate 1mL growth bacterial media containing 50ng/uL kanamycin in a 5mL culture tube. Grow to saturation in 37°C shaker (Note: this step can be done overnight in 5mL media).
    g. Dilute saturated solution 100-fold into 50mL pre-warmed growth media containing 50ng/uL kanamycin in 125mL culture flask and grow to $OD_{600}$ 0.4-0.5 (approximately 2-3hrs).
    h. Transfer 50mL culture to centrifuge tube on ice and chill 15min.
    i. Centrifuge cells at 4000rcf for 10min at 4°C.
    j. Decant supernatant and resuspend pellet in 5mL TSS solution.
    k. Pipet chemically competent “GFP-itis" cells in 100uL aliquots in 1.7mL centrifuge tubes (makes ~50 aliquots, each student group will use two).
7. Prepare solutions of “GFP-itis pBE-t” and “GFP-itis pBE-nt” at ~10ng/uL in sterile water

*Student base editing activity:*

1. Prepare two aliquots of 70uL sterile DI water + 20uL 5x KCM in a 1.7mL snap top tube on ice. To one, add 1uL “GFP-itis pBE-t” (10-100ng DNA) and to the other 1uL “GFP-itis” pBE-nt (10-100ng DNA). Rest on ice 5 minutes. Note: to conserve time, this step can be done by the researchers prior to beginning the activity.
2. Treat the “GFP-itis" cells by adding a 100uL aliquot of “GFP-itis” bacterial cells to each pBE solution. Incubate on ice additional 5 minutes.
3. Heat shock for 75 seconds in 42°C water bath. Return to ice immediately, and rest for 2 minutes.

4. Add 800uL of sterile bacterial media for recovery. Recover for 10min-1hr (Note longer recovery will result in more colonies), shaking at 37°C. Note: shaking is optional and the recovery can be done in an incubator if there is not access to an incubator/shaker in the school, though transformation efficiency and number of resultant colonies may be impacted.
5. After recovery, spread 30-50uL of the transformed culture on agar plates containing 50ng/uL kanamycin, 50ng/uL carbenicillin (or ampicillin) and 1mM theophylline.
6. Allow to grow at 37°C for 24hrs. Image plates with transilluminator or handheld blue light using blue light filter glasses and compare GFP correction in targeting (pBE-t) and non-targeting samples (pBE-nt).

Supplemental Figure 1

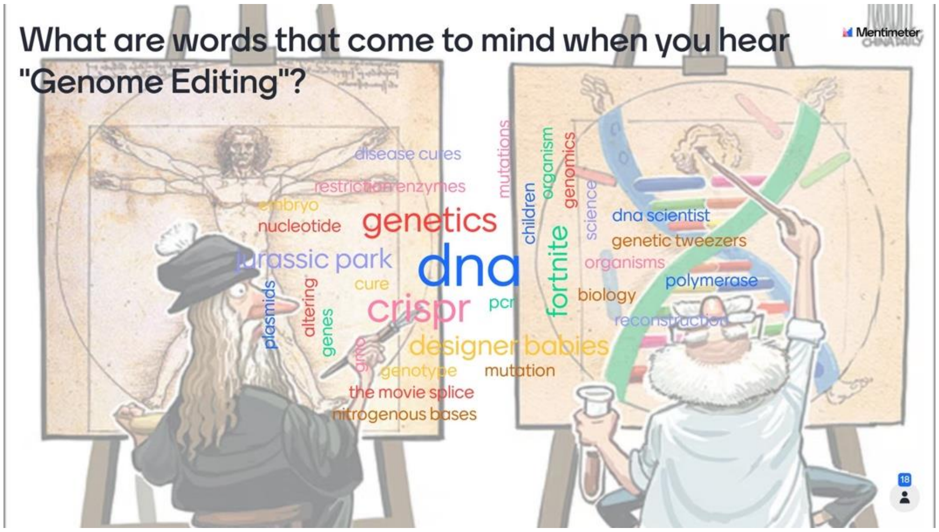


Supplemental Figure 1. Mentimeter results for class session 1. Students were asked to insert three words that come to mind when hearing the word “Genome Editing.” Most used terms are larger in size. In total, 19 students participated in this activity.
