## Supplemental Material - Instructor Feedback for "Curing “GFP-itis” in Bacteria with Base Editors: Development of a Genome Editing Science Program Implemented with High School Biology Students"

8. Is there any part of the activity you would drop? Anything you would add? Leave any additional comments or feedback here!

**Teacher Program Evaluation Survey**

1. What are your perspectives on the current approach used in this program and its ability to foster excitement in further STEM education?

The UCSD team was very enthusiastic from day 1. They made students feel comfortable to ask questions and gave multiple examples to make learning relatable and applicable.

2. What are your perspectives about the potential of implementing this project in other schools?

This program that was developed was well executed + ran seemlessly. It worked extremely well with my senior biomed course and would be amazing to implement to AP BIO courses as well. I think students who have an understanding of biology truly benefit from this program

3. Are there any activities within the 3-day that you feel can be improved? What aspects of the program do you think worked well?

Honestly - this program was done so well that I do not have any recommendations for improvement. Each day brought something new + kept students engaged.

4. Please provide any suggestions and feedback on how to better improve the program.

My students found the grad students very relatable + responded well to them. Some comments that I heard as I circled the room were - "That was so fun!" and "Wow - look at all this cool lab equipment". and "Wow - did Dr. Komor really develop this technology!?" She is so humble.

* They felt very lucky that Dr. Komor was present for the experiment as well.
