## Supplemental Material - Lecture Slides for "Curing “GFP-itis” in Bacteria with Base Editors: Development of a Genome Editing Science Program Implemented with High School Biology Students"

### Slide 1
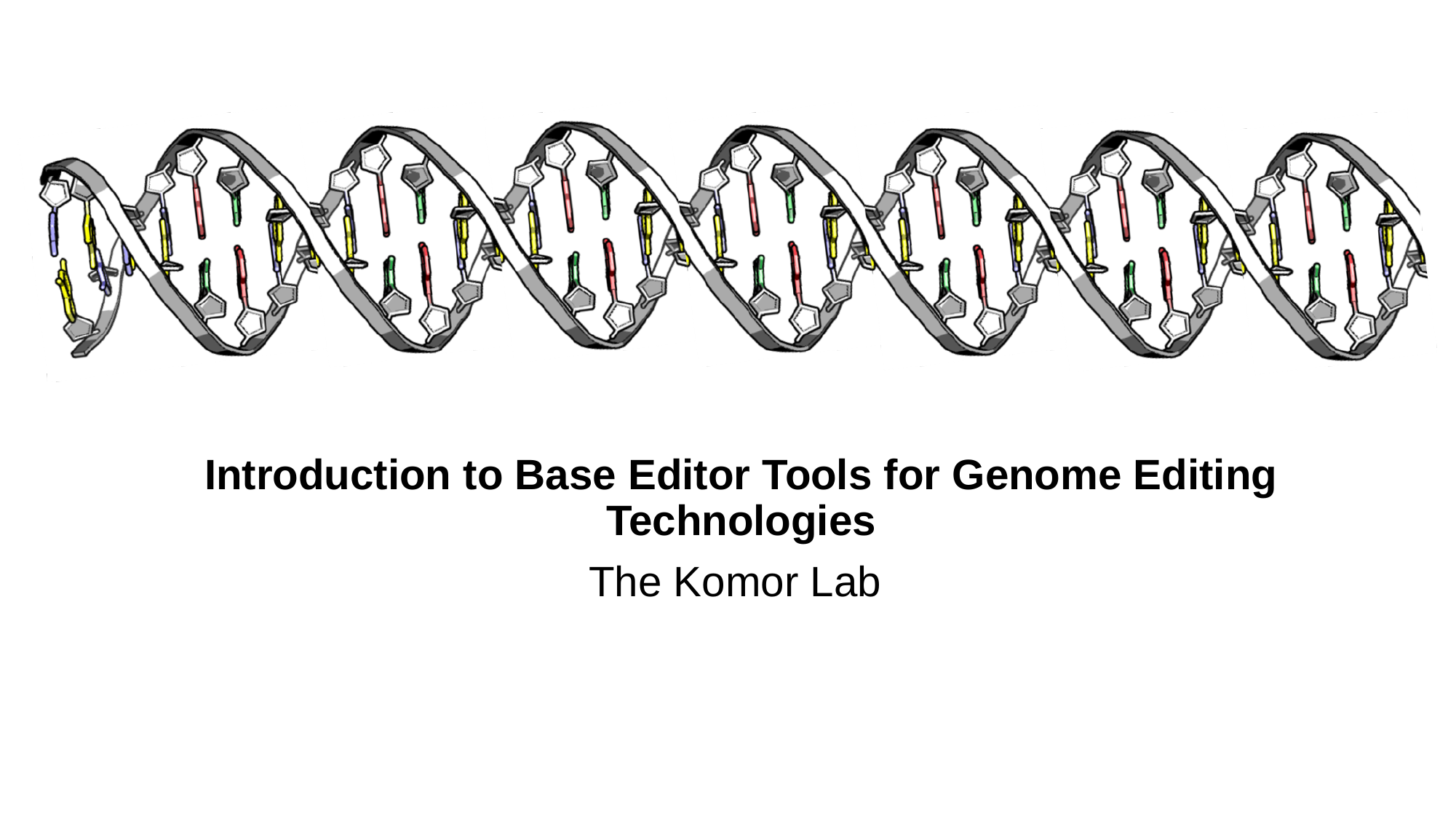

Introduction to Base Editor Tools for Genome Editing Technologies
The Komor Lab

### Slide 2
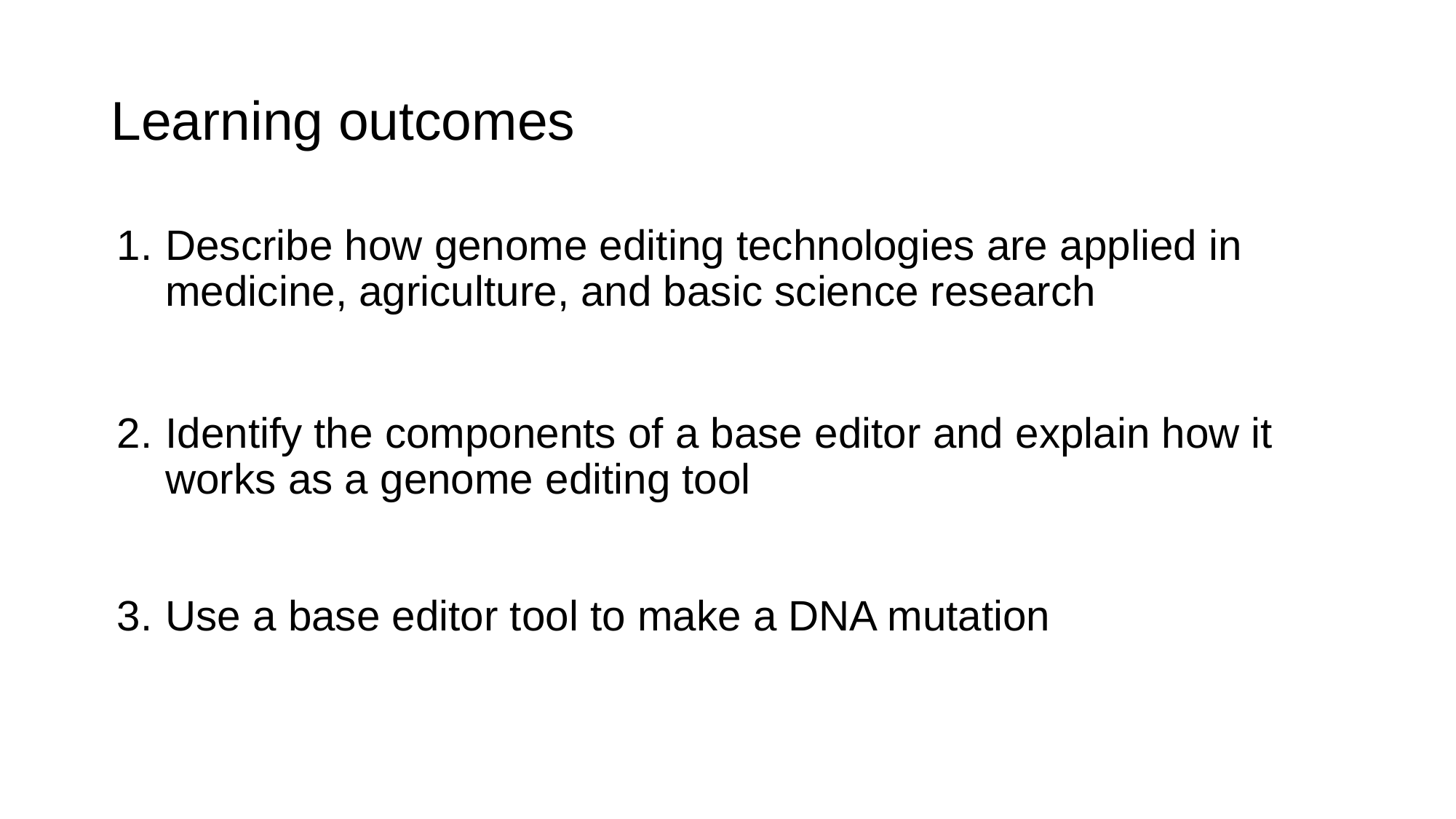

#### Learning outcomes
Describe how genome editing technologies are applied in medicine, agriculture, and basic science research
Identify the components of a base editor and explain how it works as a genome editing tool
Use a base editor tool to make a DNA mutation

### Slide 3
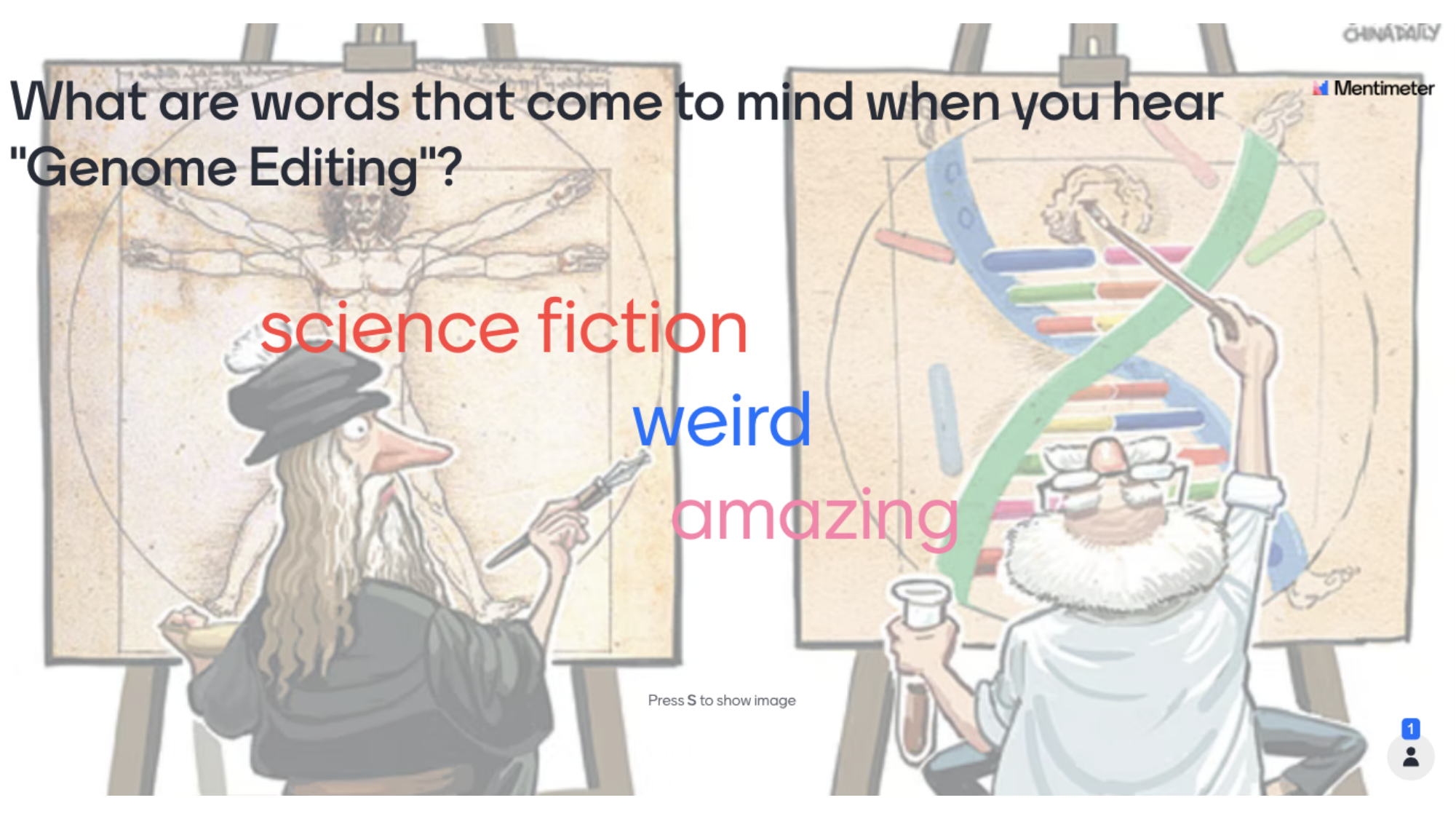

#

### Slide 4
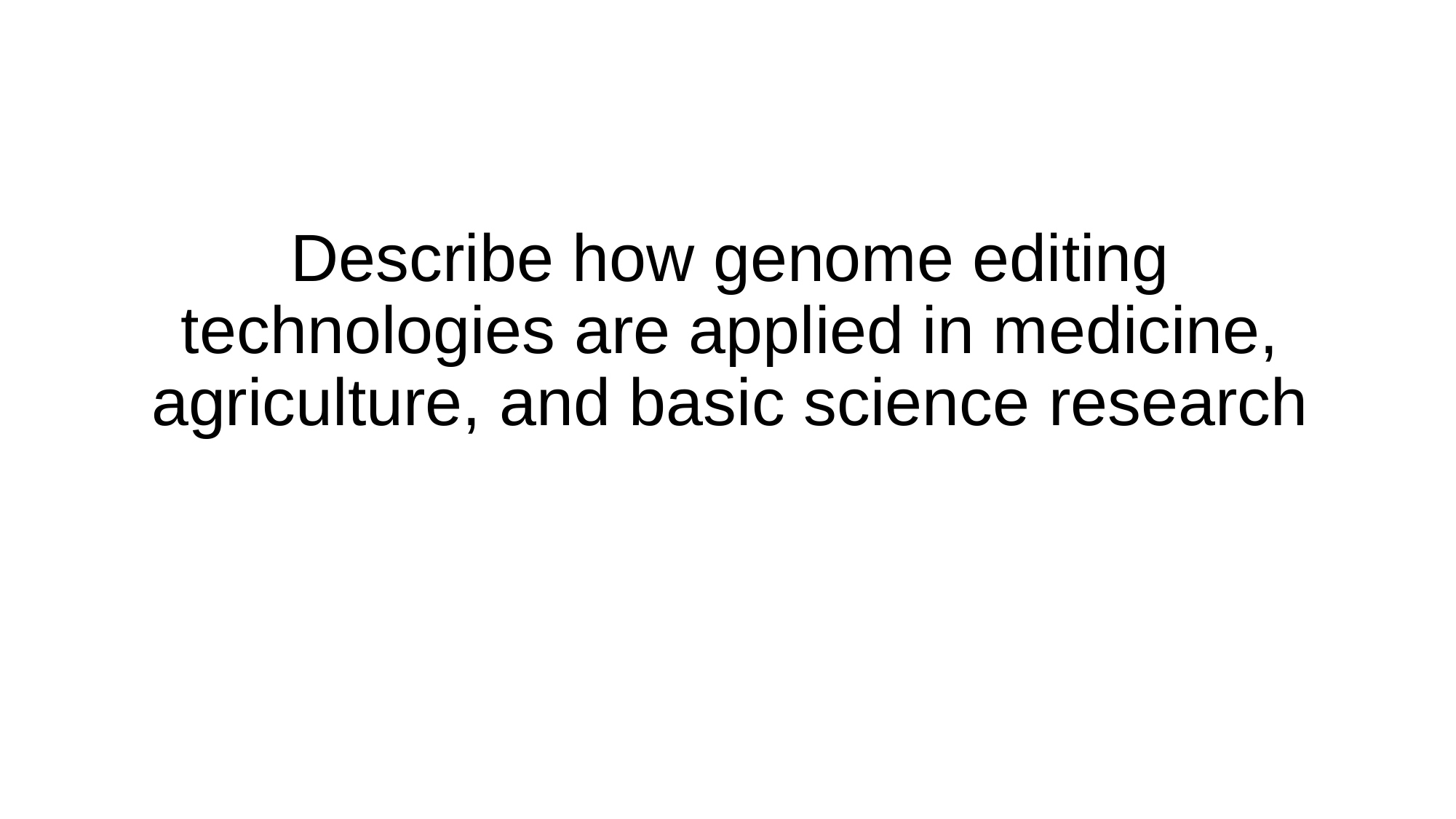

Describe how genome editing technologies are applied in medicine, agriculture, and basic science research

### Slide 5
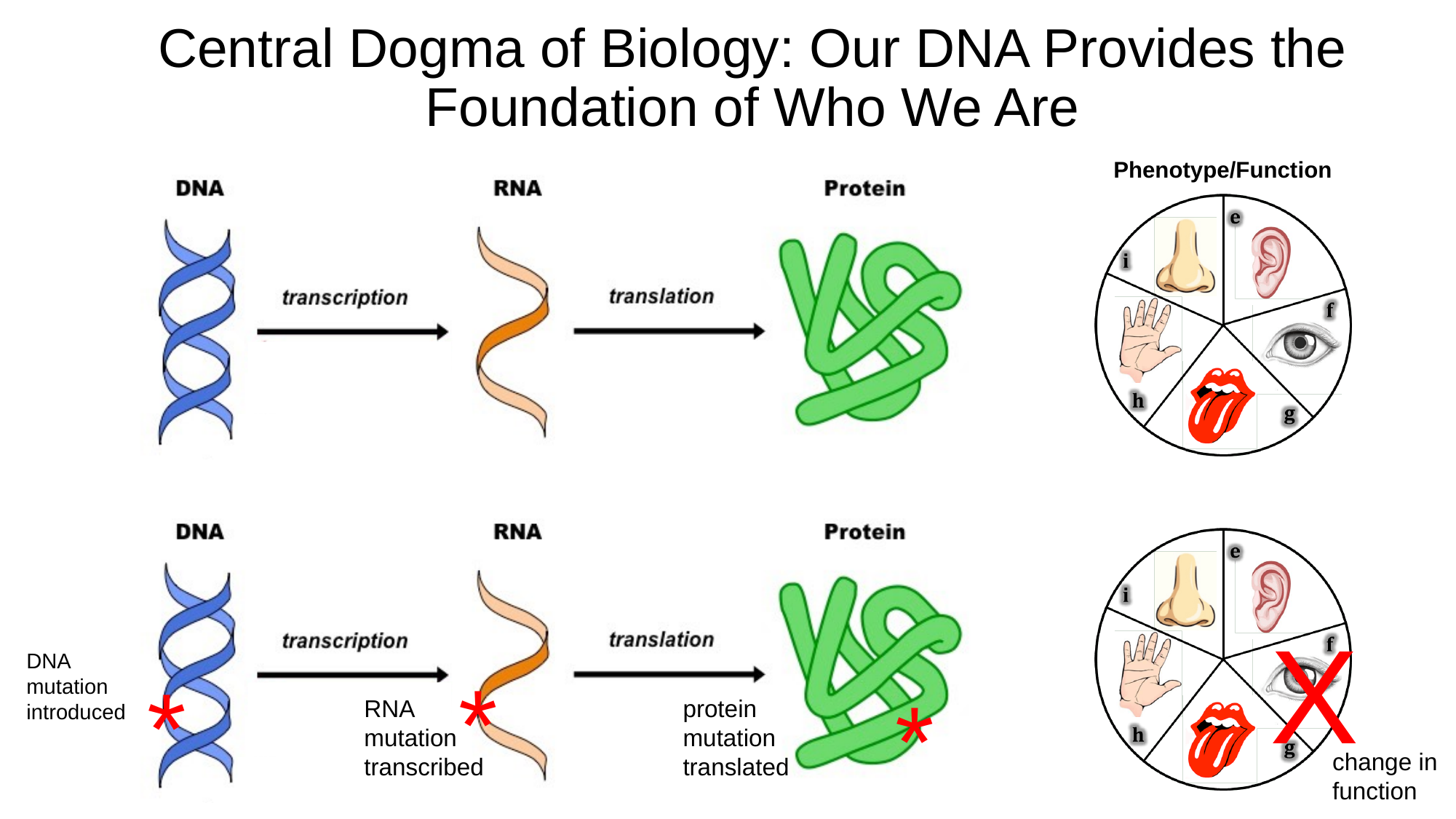

#### Central Dogma of Biology: Our DNA Provides the Foundation of Who We Are
Phenotype/Function
X
DNA
mutation introduced
*
*
*
RNA
mutation transcribed
protein mutation translated
change in function

### Slide 6
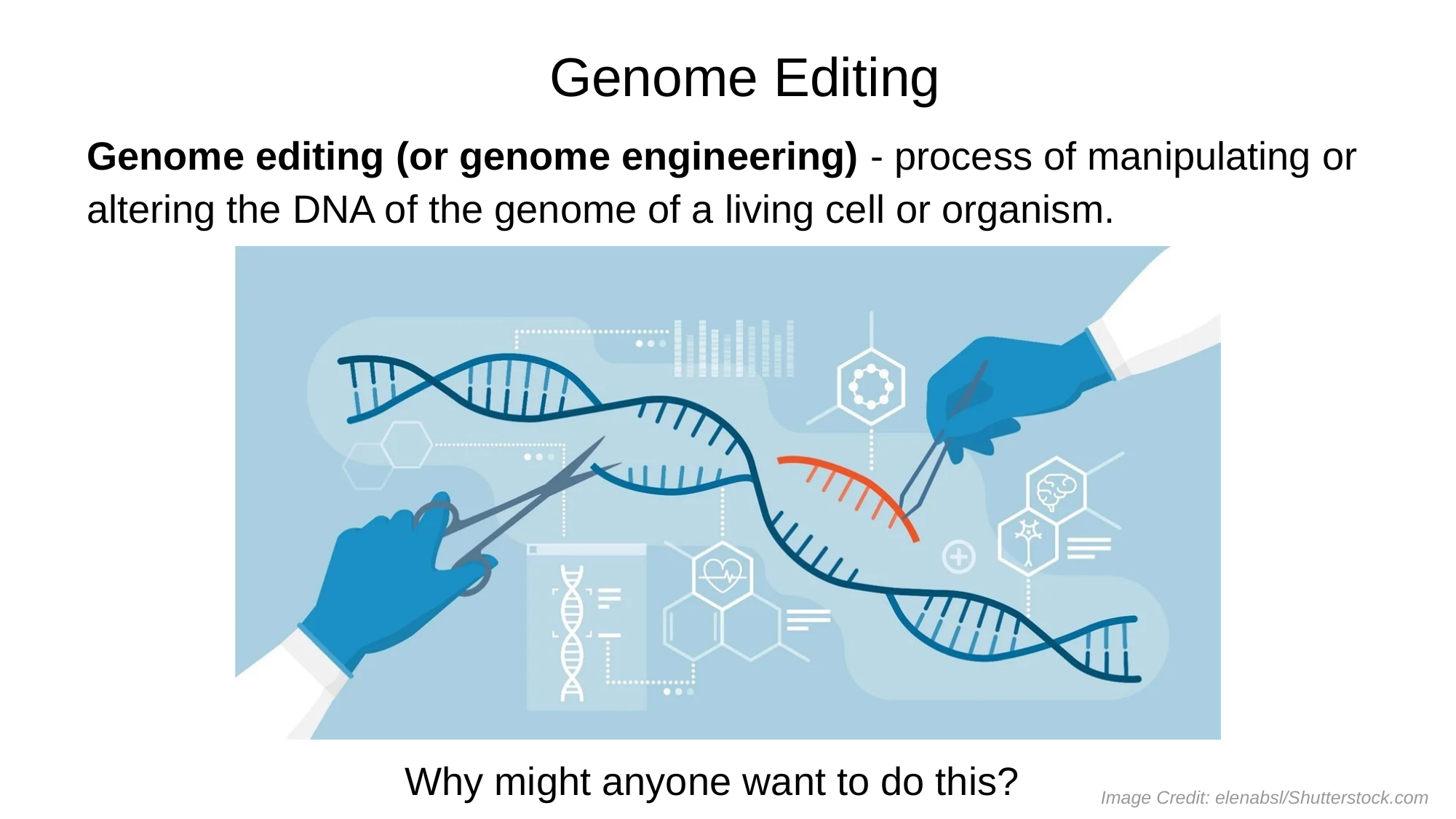

#### Genome Editing
Genome editing (or genome engineering) - process of manipulating or altering the DNA of the genome of a living cell or organism.
Why might anyone want to do this?
Image Credit: elenabsl/Shutterstock.com

### Slide 7
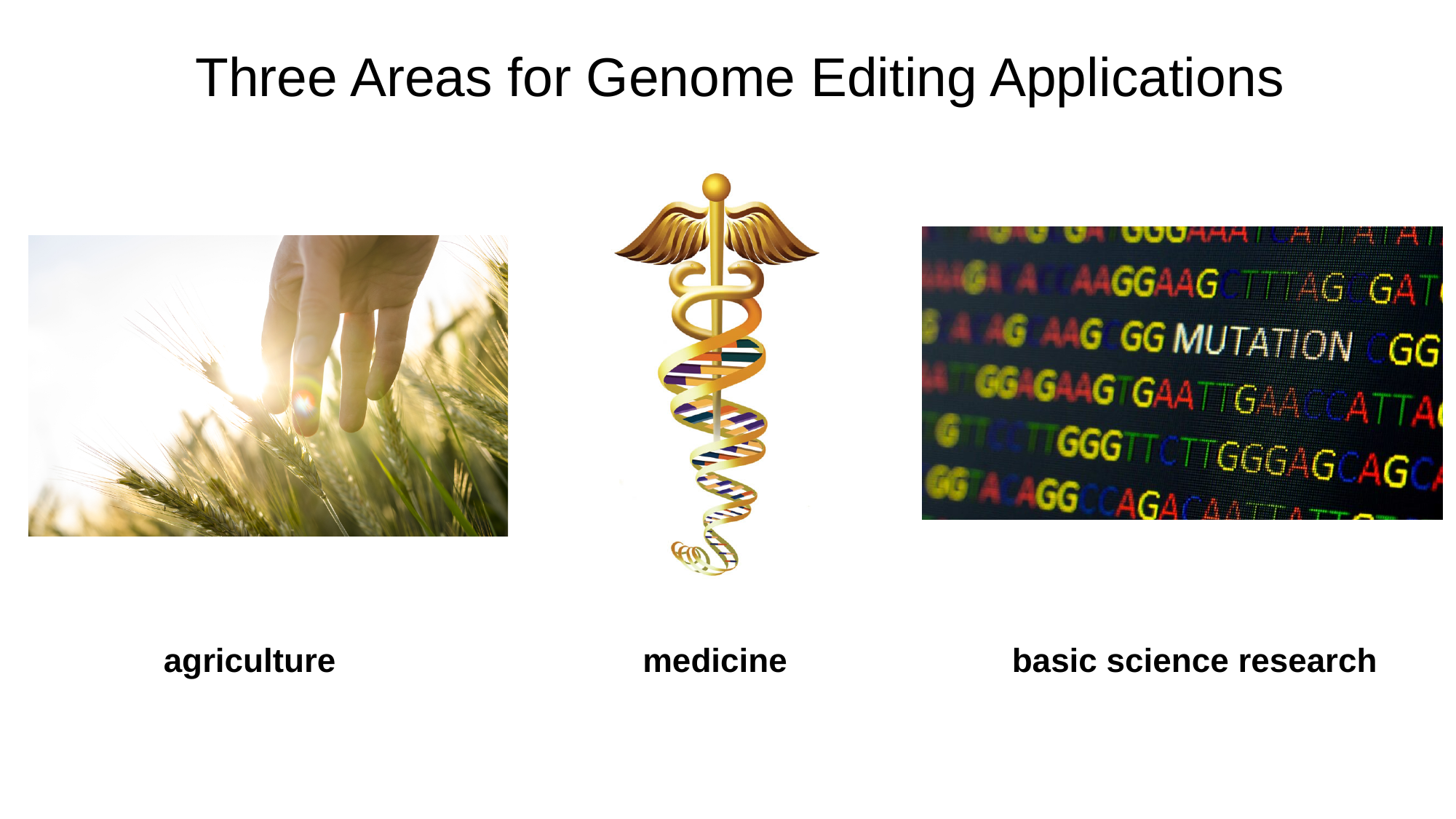

#### Three Areas for Genome Editing Applications
agriculture
medicine
basic science research

### Slide 8
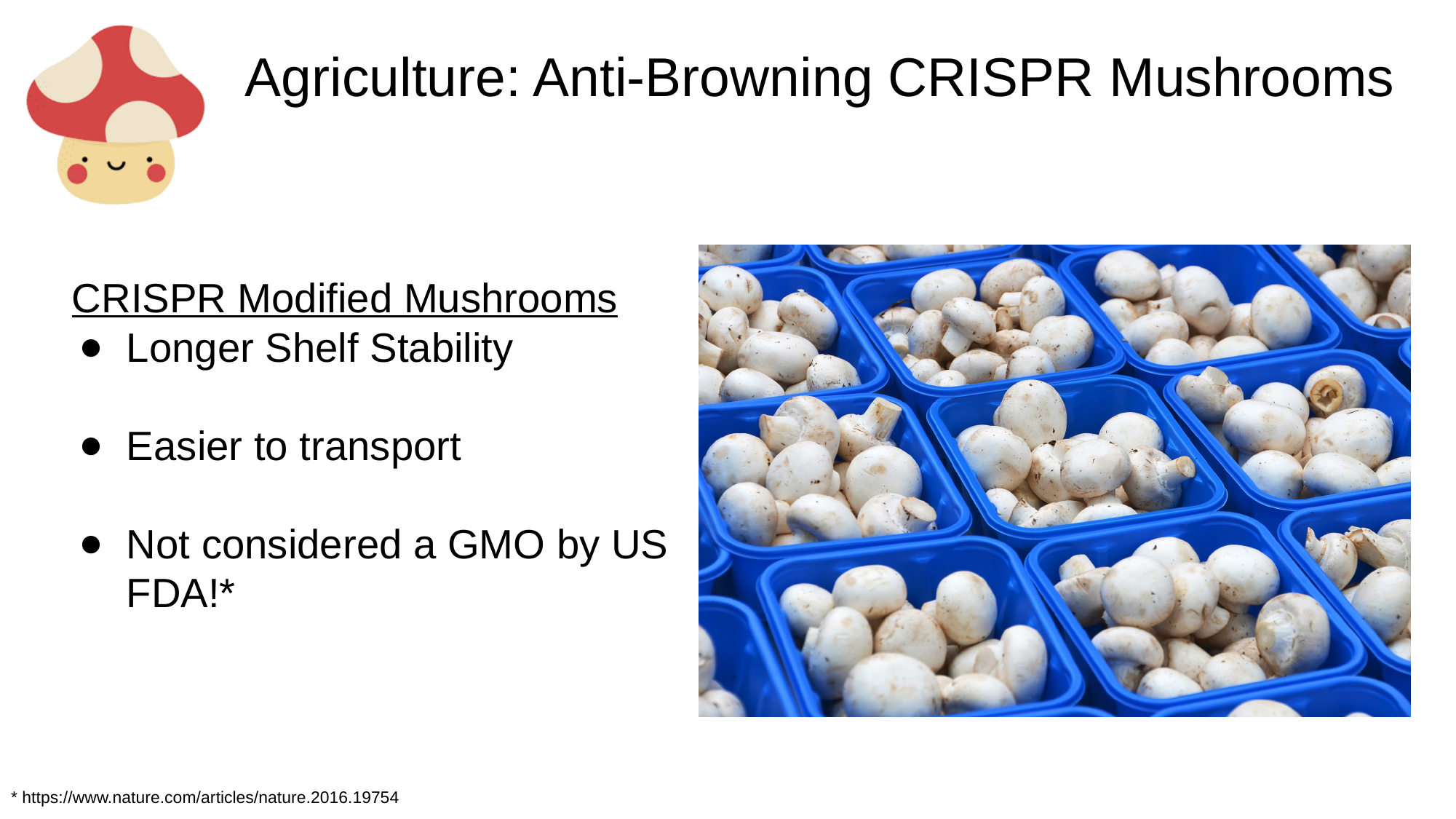

#### Agriculture: Anti-Browning CRISPR Mushrooms
CRISPR Modified Mushrooms
Longer Shelf Stability
Easier to transport
Not considered a GMO by US FDA!*
* https://www.nature.com/articles/nature.2016.19754

### Slide 9
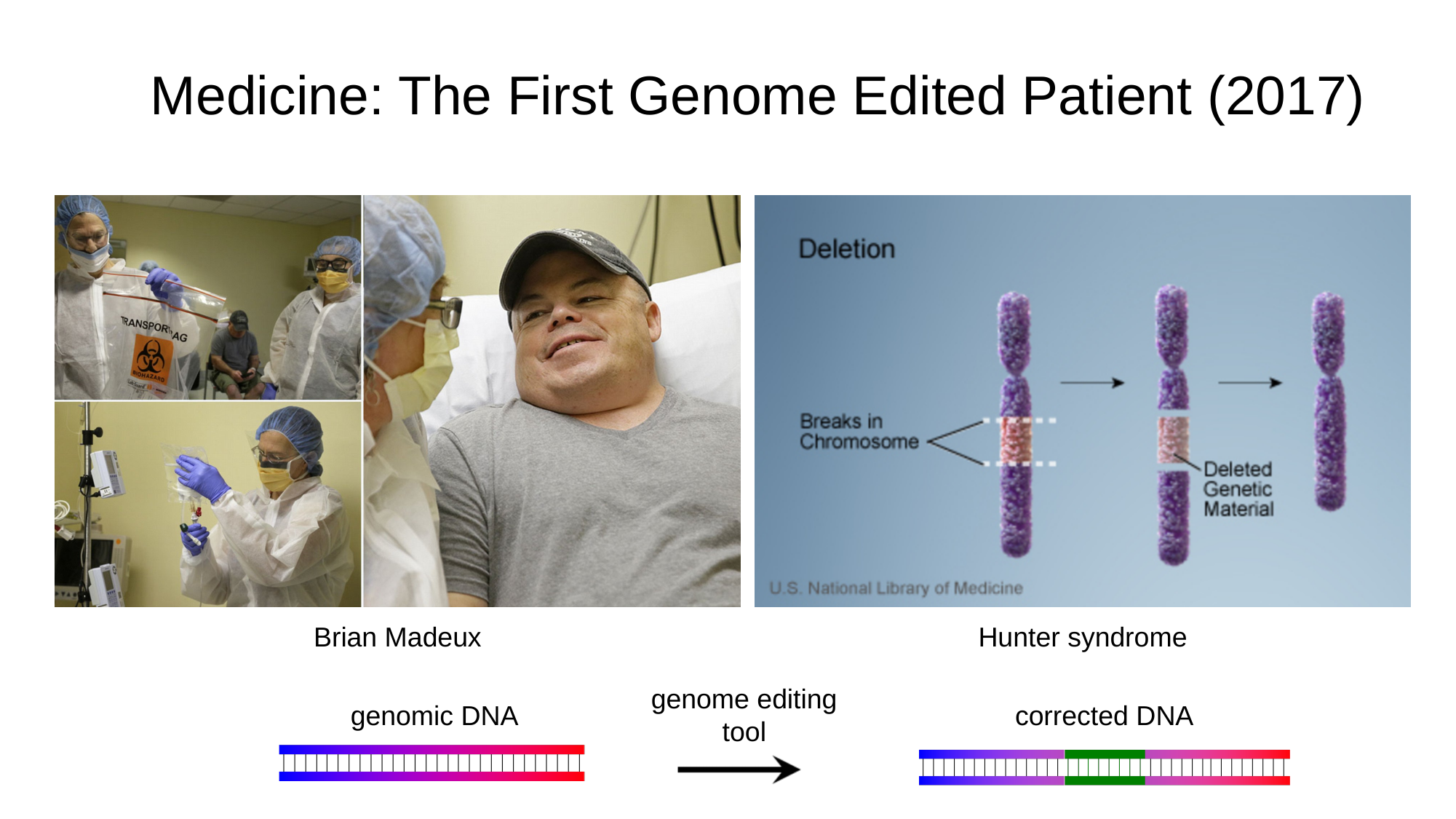

#### Medicine: The First Genome Edited Patient (2017)
Brian Madeux
Hunter syndrome
genome editing tool
genomic DNA
corrected DNA

### Slide 10
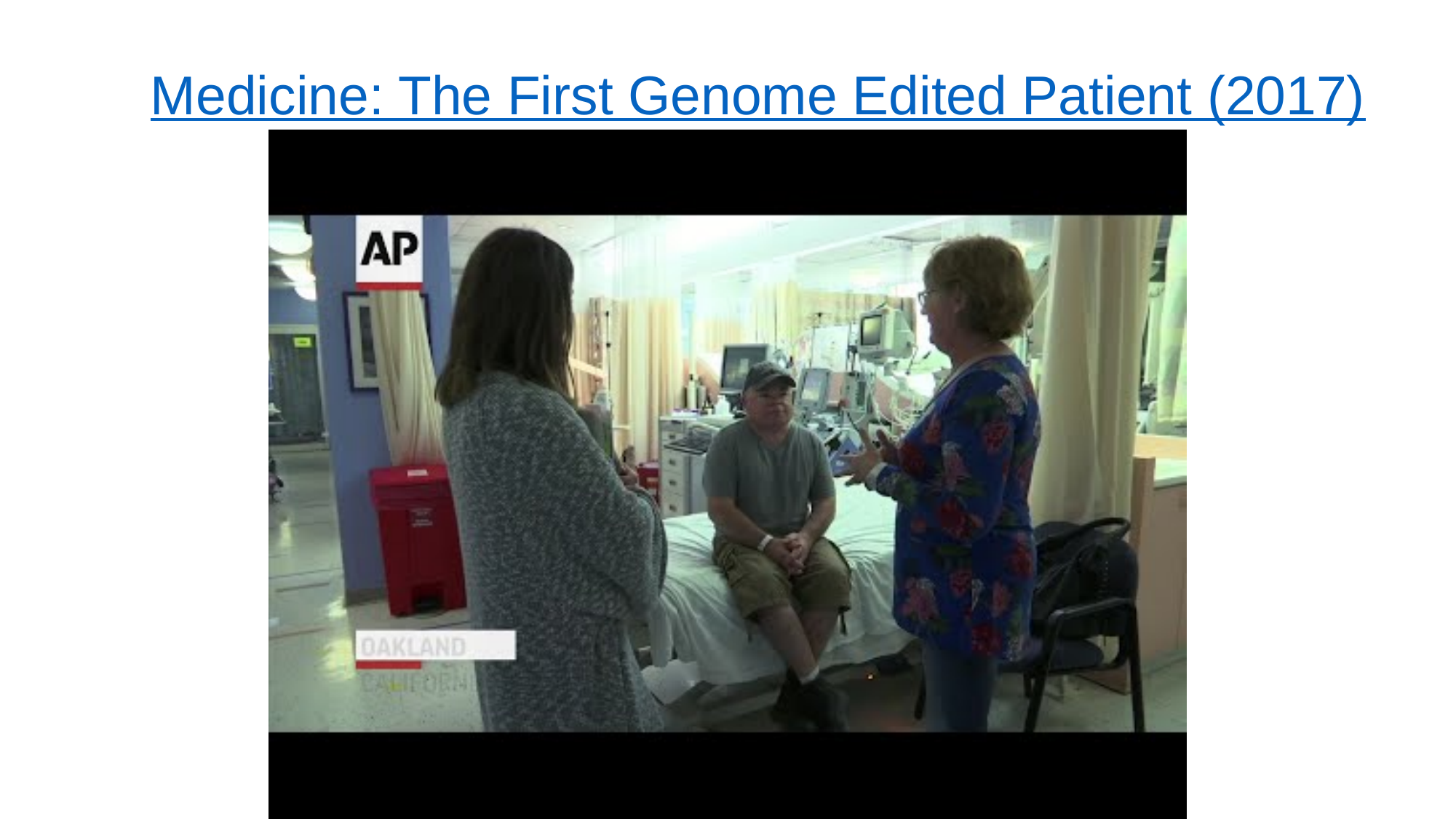

#### Medicine: The First Genome Edited Patient (2017)

### Slide 11
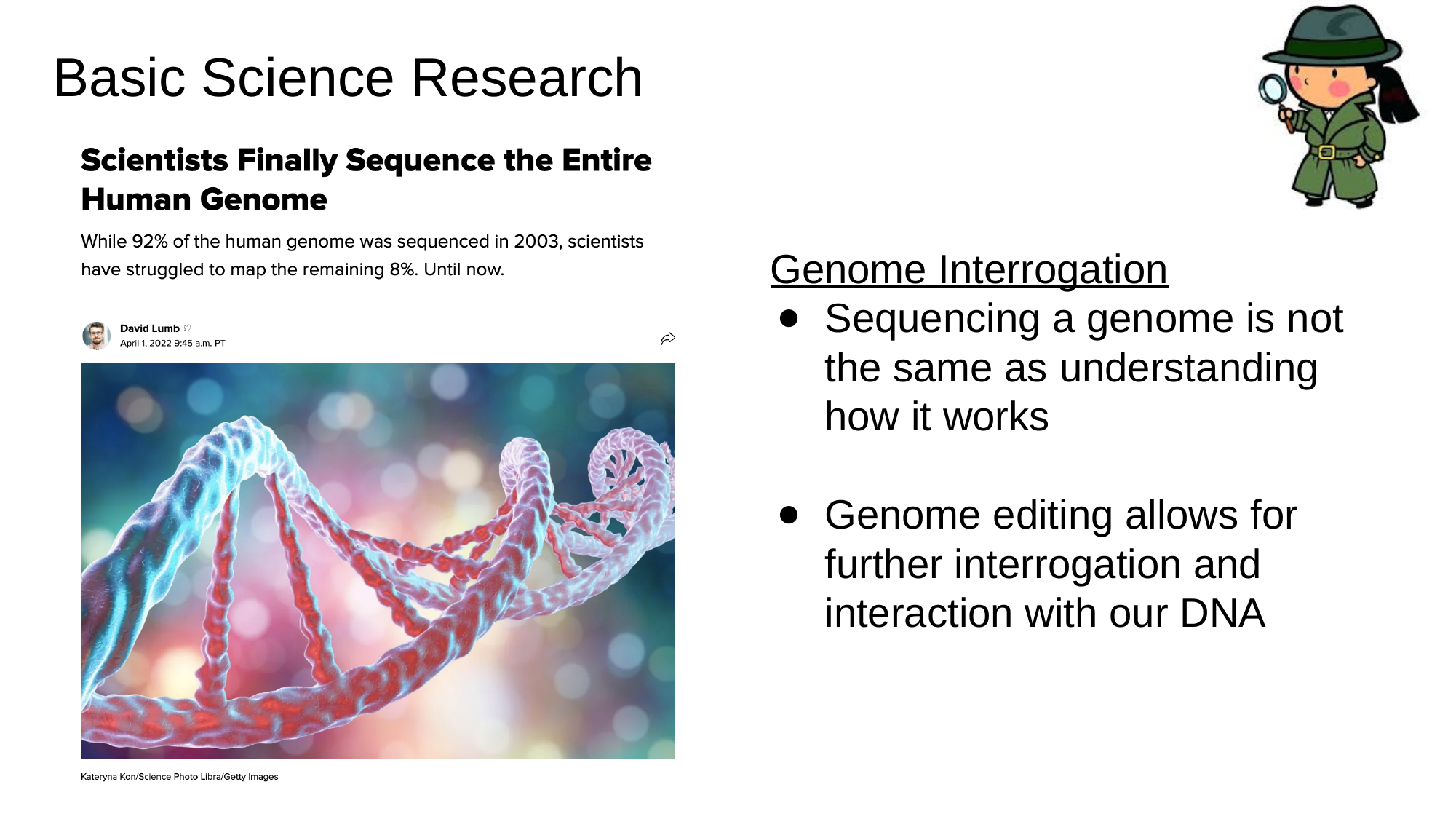

#### Basic Science Research
Genome Interrogation
Sequencing a genome is not the same as understanding how it works
Genome editing allows for further interrogation and interaction with our DNA

### Slide 12
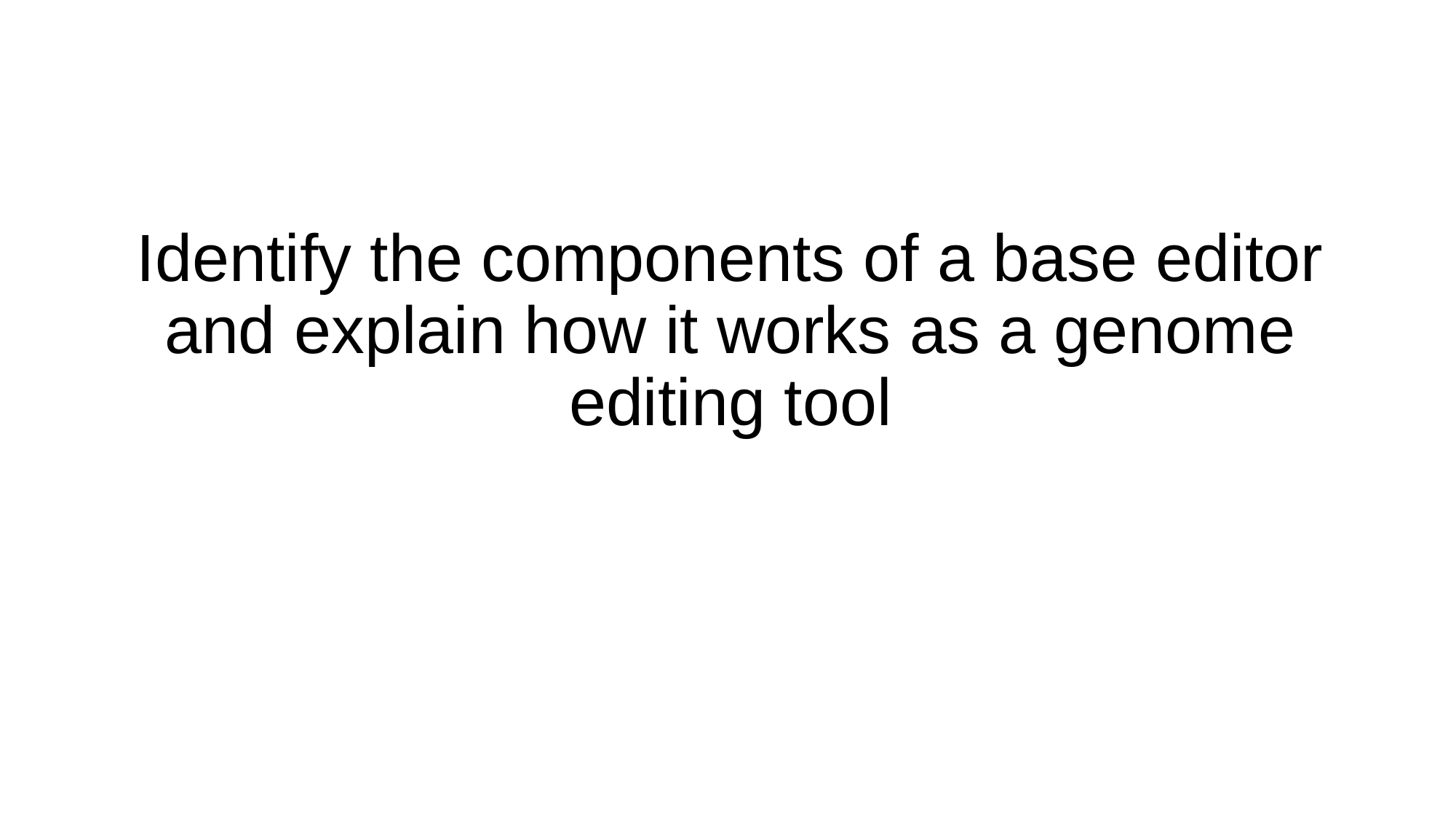

Identify the components of a base editor and explain how it works as a genome editing tool

### Slide 13
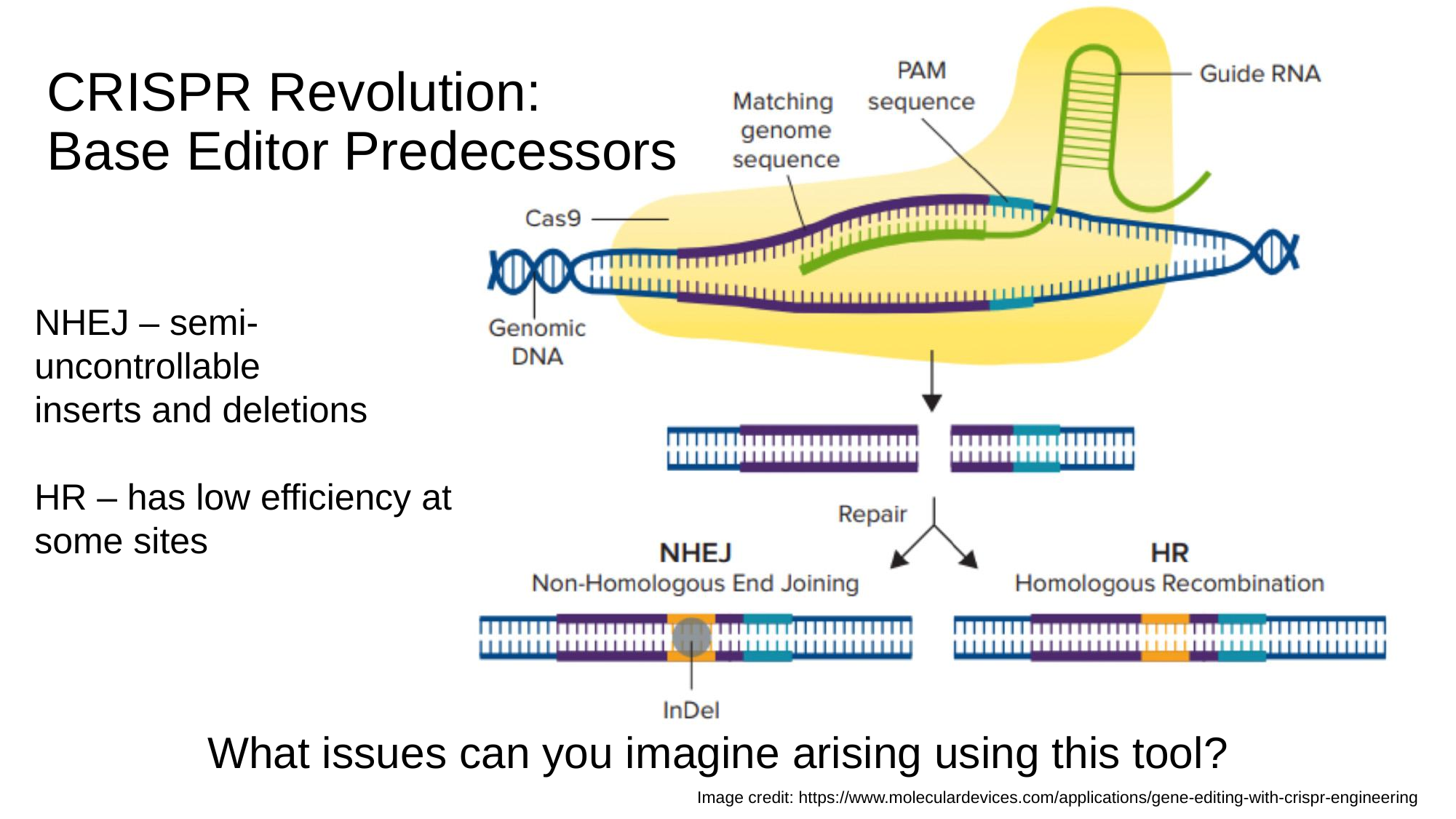

#### CRISPR Revolution: Base Editor Predecessors
NHEJ – semi-uncontrollable
inserts and deletions
HR – has low efficiency at some sites
What issues can you imagine arising using this tool?
Image credit: https://www.moleculardevices.com/applications/gene-editing-with-crispr-engineering

### Slide 14
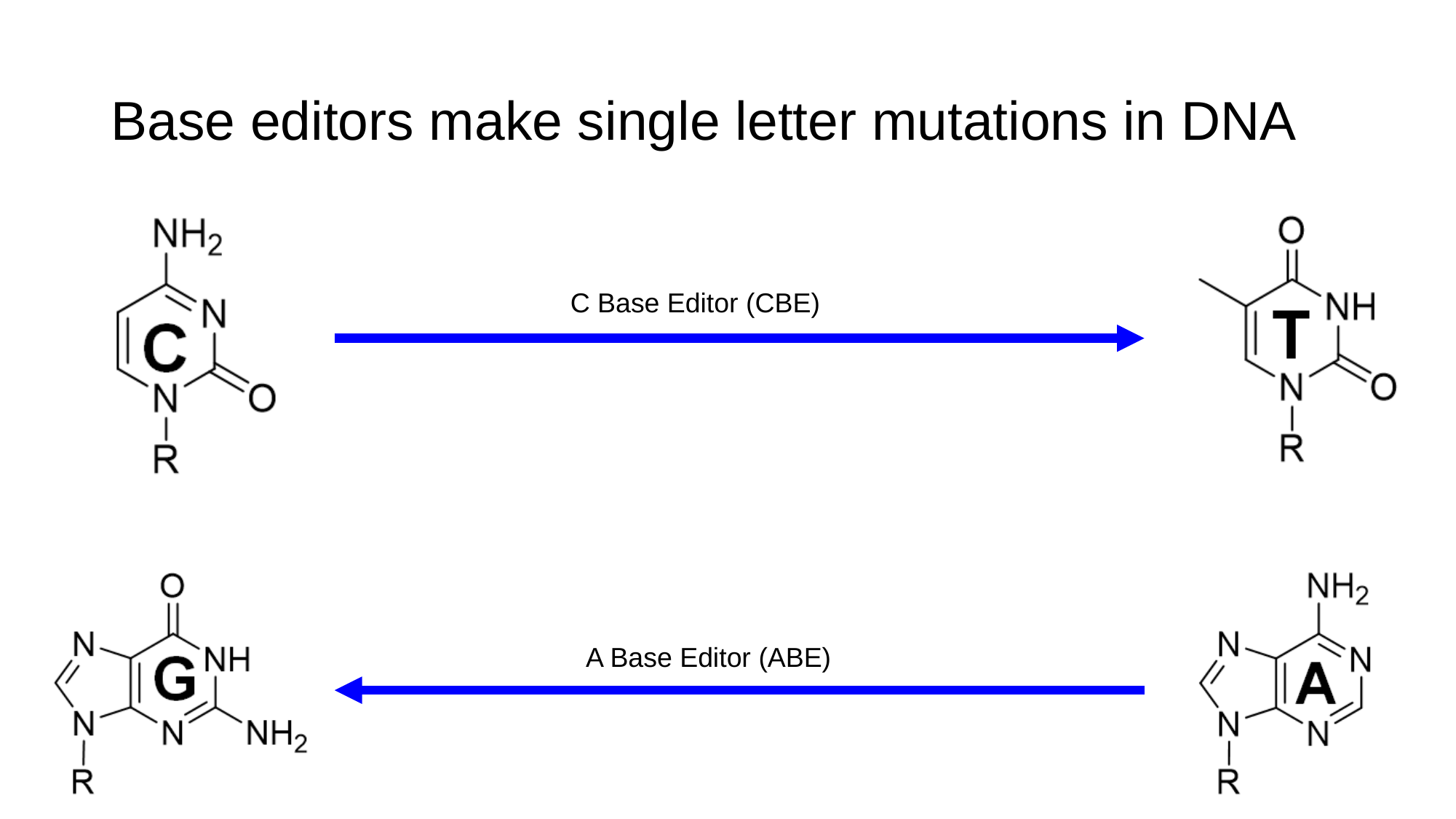

#### Base editors make single letter mutations in DNA
C Base Editor (CBE)
A Base Editor (ABE)

### Slide 15
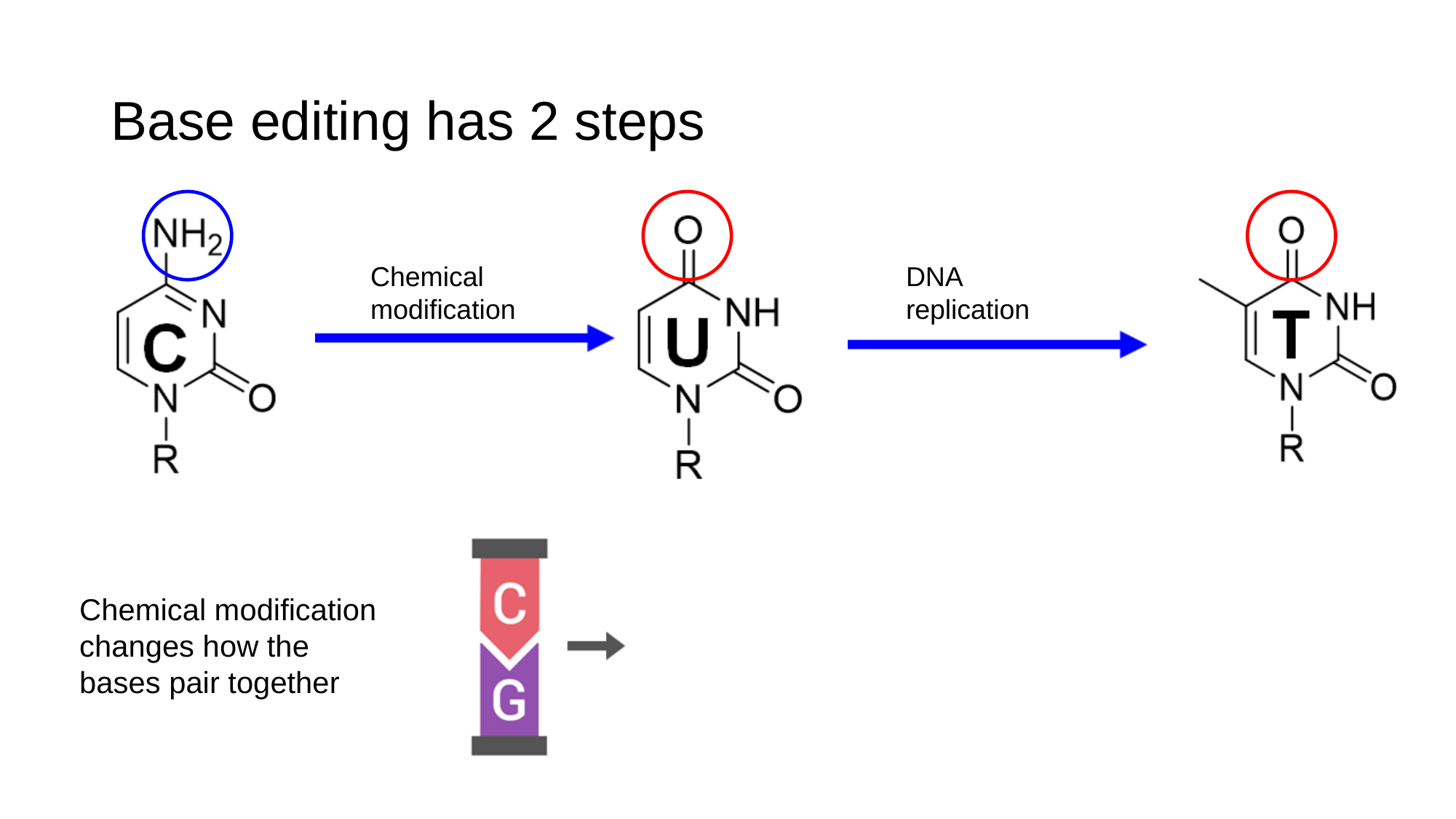

#### Base editing has 2 steps
Chemical modification
DNA replication
Chemical modification changes how the bases pair together

### Slide 16
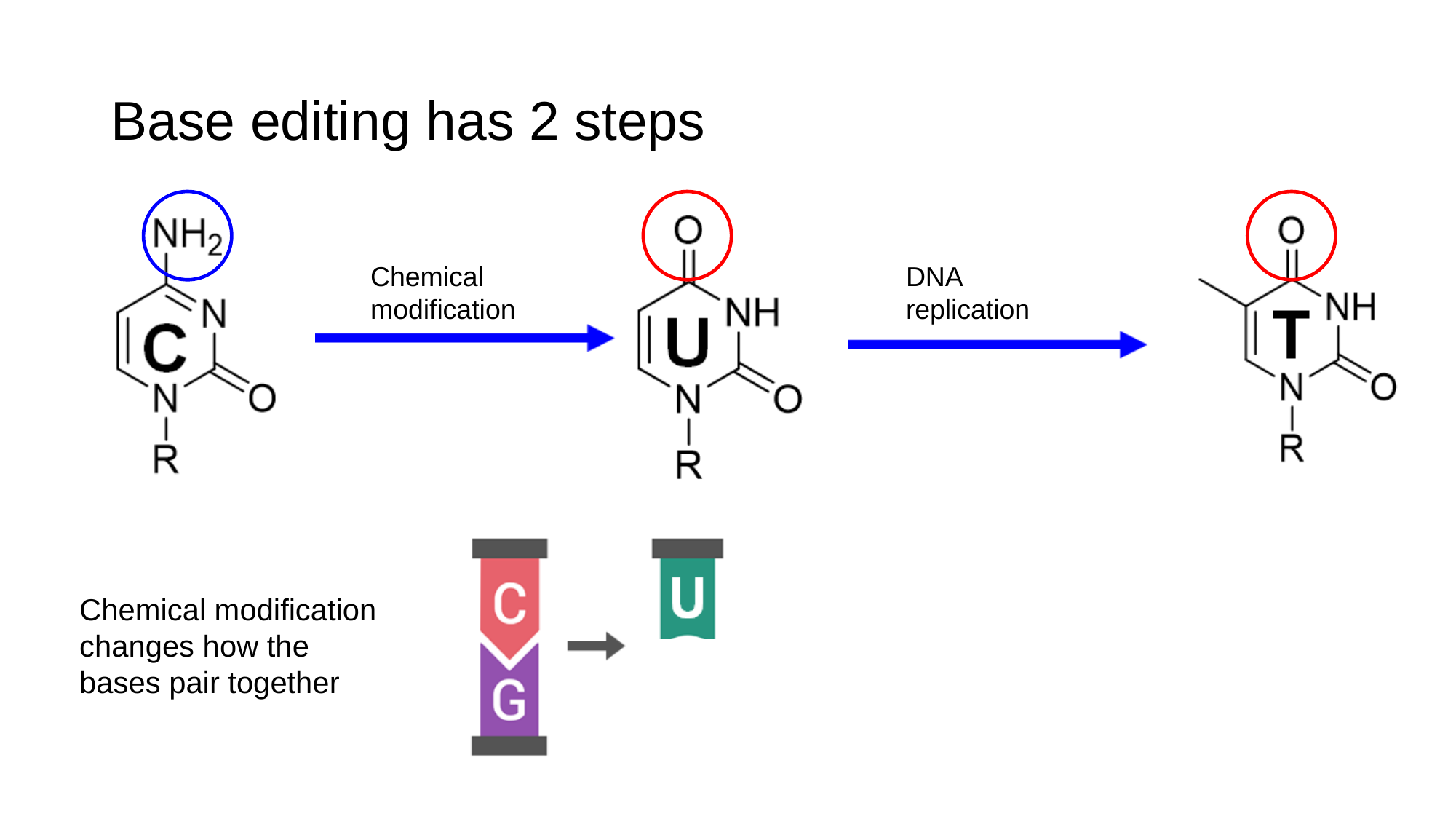

#### Base editing has 2 steps
Chemical modification
DNA replication
Chemical modification changes how the bases pair together

### Slide 17
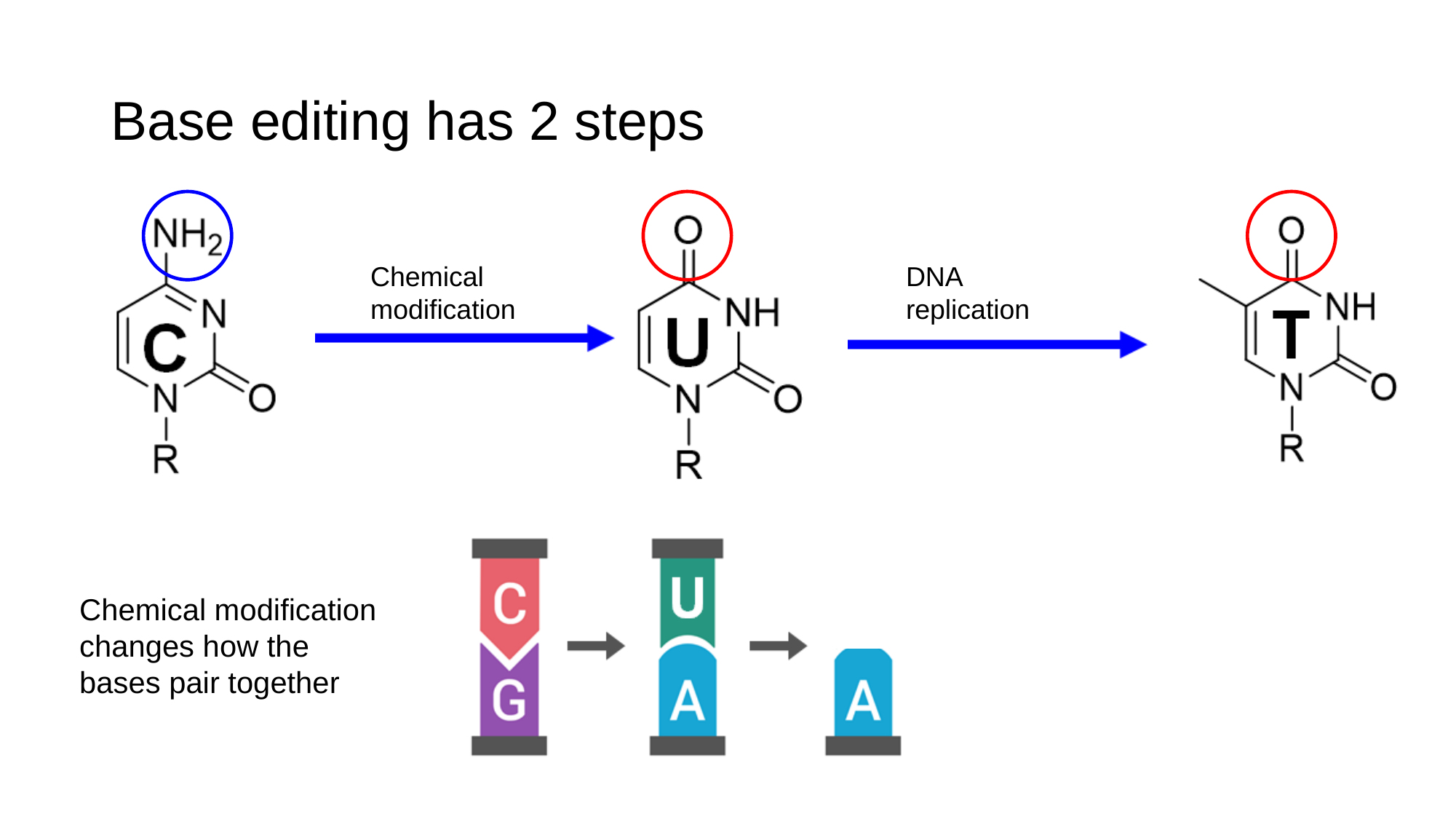

#### Base editing has 2 steps
Chemical modification
DNA replication
Chemical modification changes how the bases pair together

### Slide 18
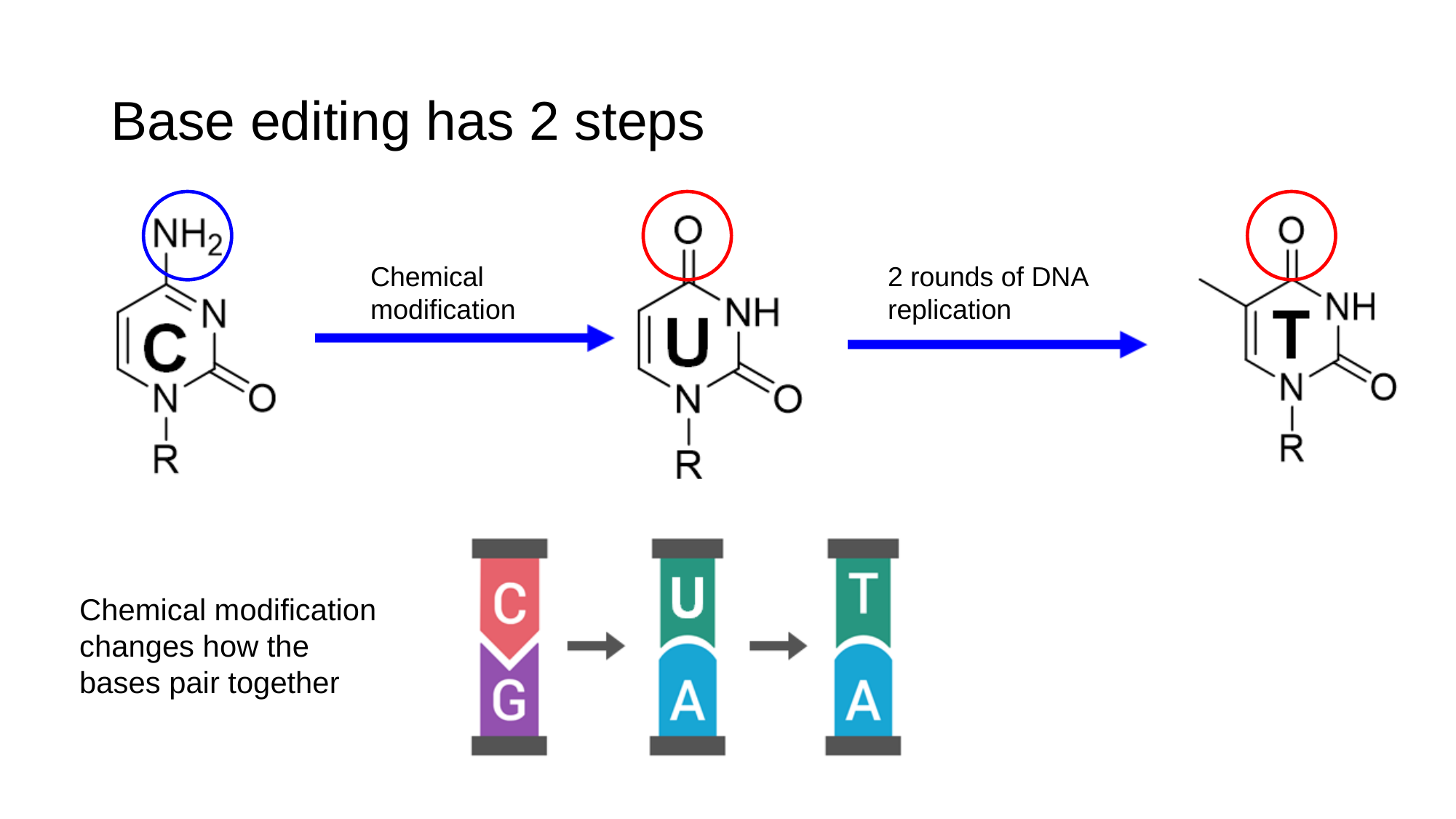

#### Base editing has 2 steps
Chemical modification
2 rounds of DNA replication
Chemical modification changes how the bases pair together

### Slide 19
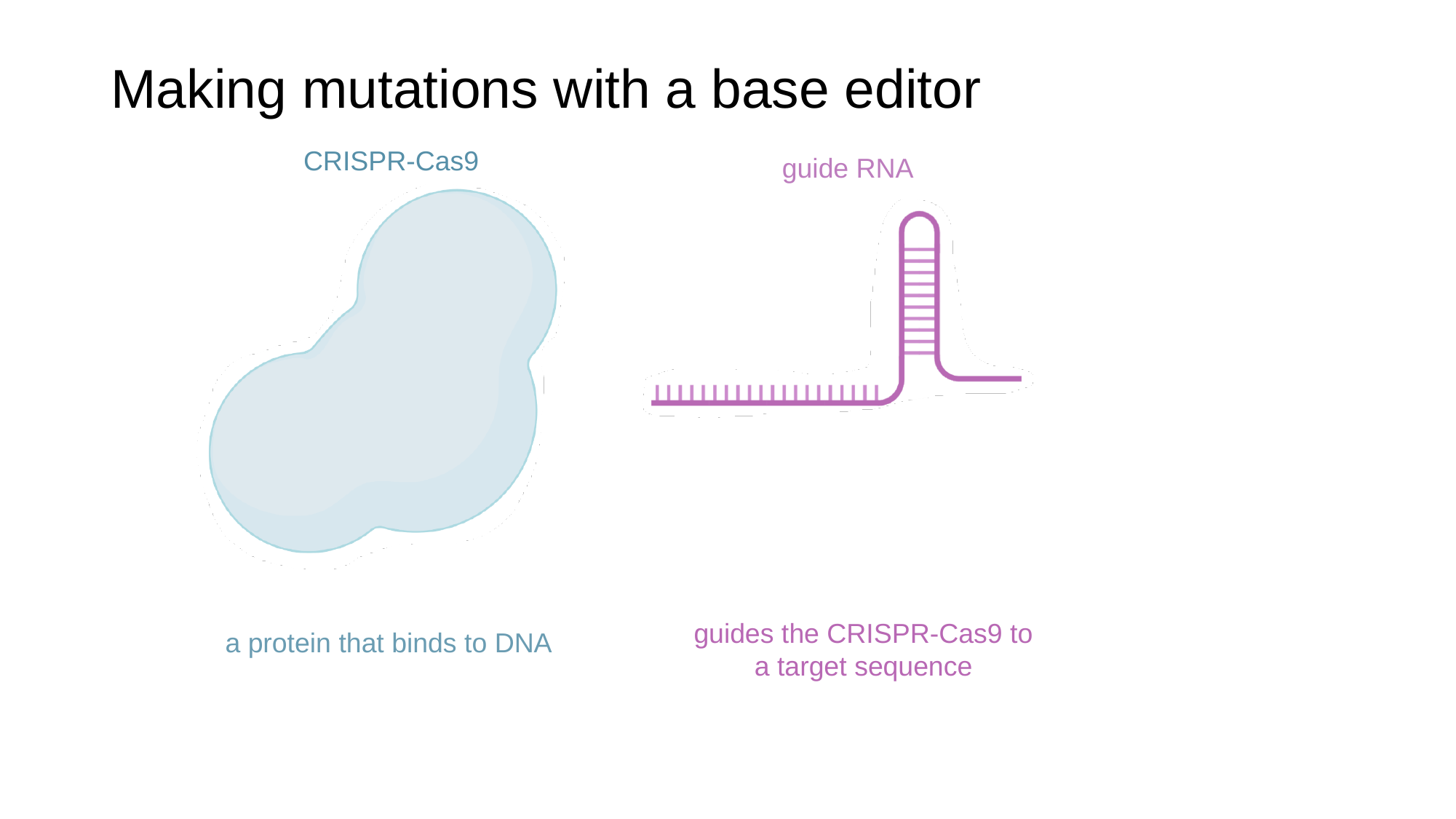

#### Making mutations with a base editor
CRISPR-Cas9
guide RNA
guides the CRISPR-Cas9 to a target sequence
a protein that binds to DNA

### Slide 20
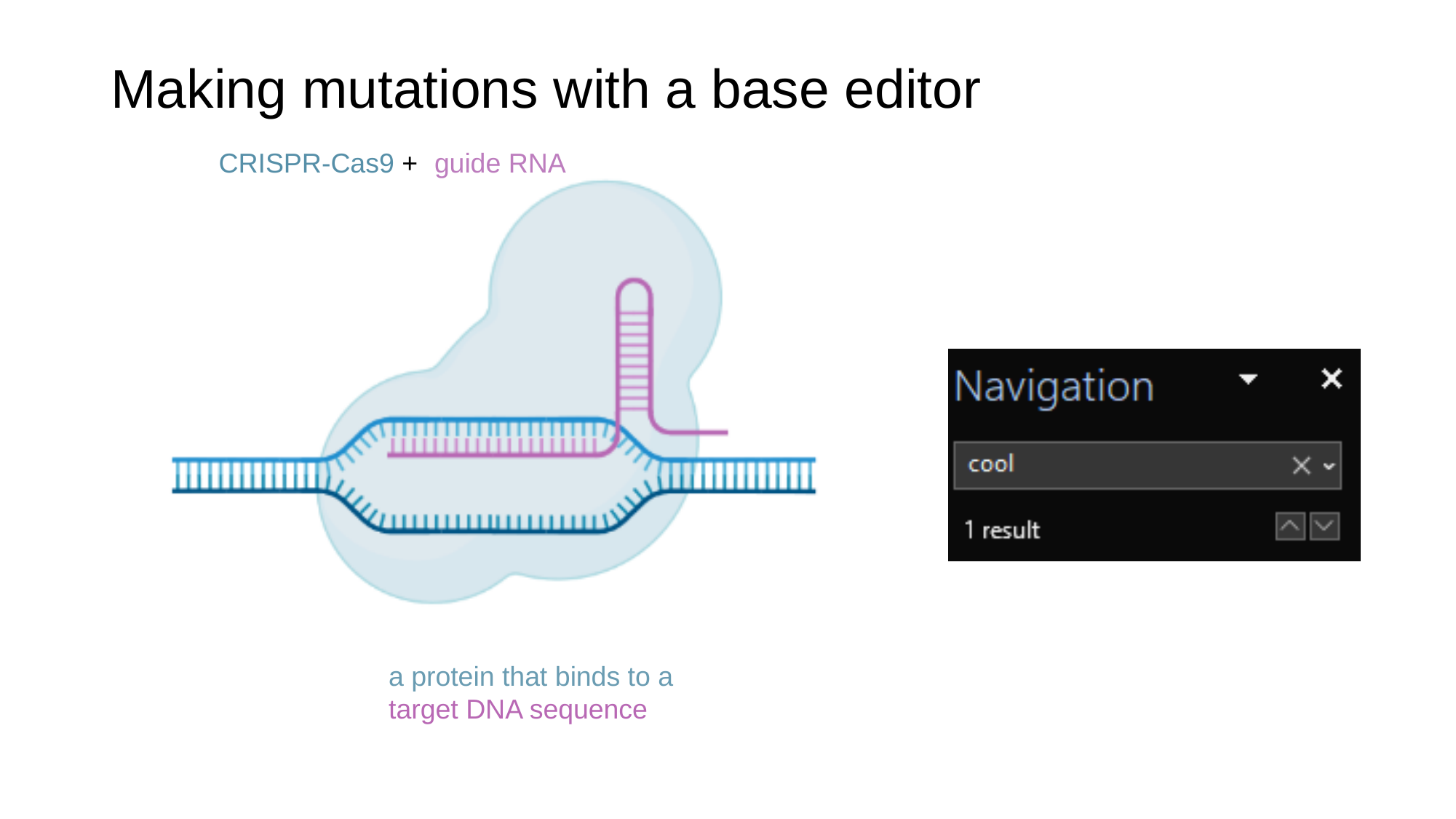

#### Making mutations with a base editor
CRISPR-Cas9 +
guide RNA
a protein that binds to a target DNA sequence

### Slide 21
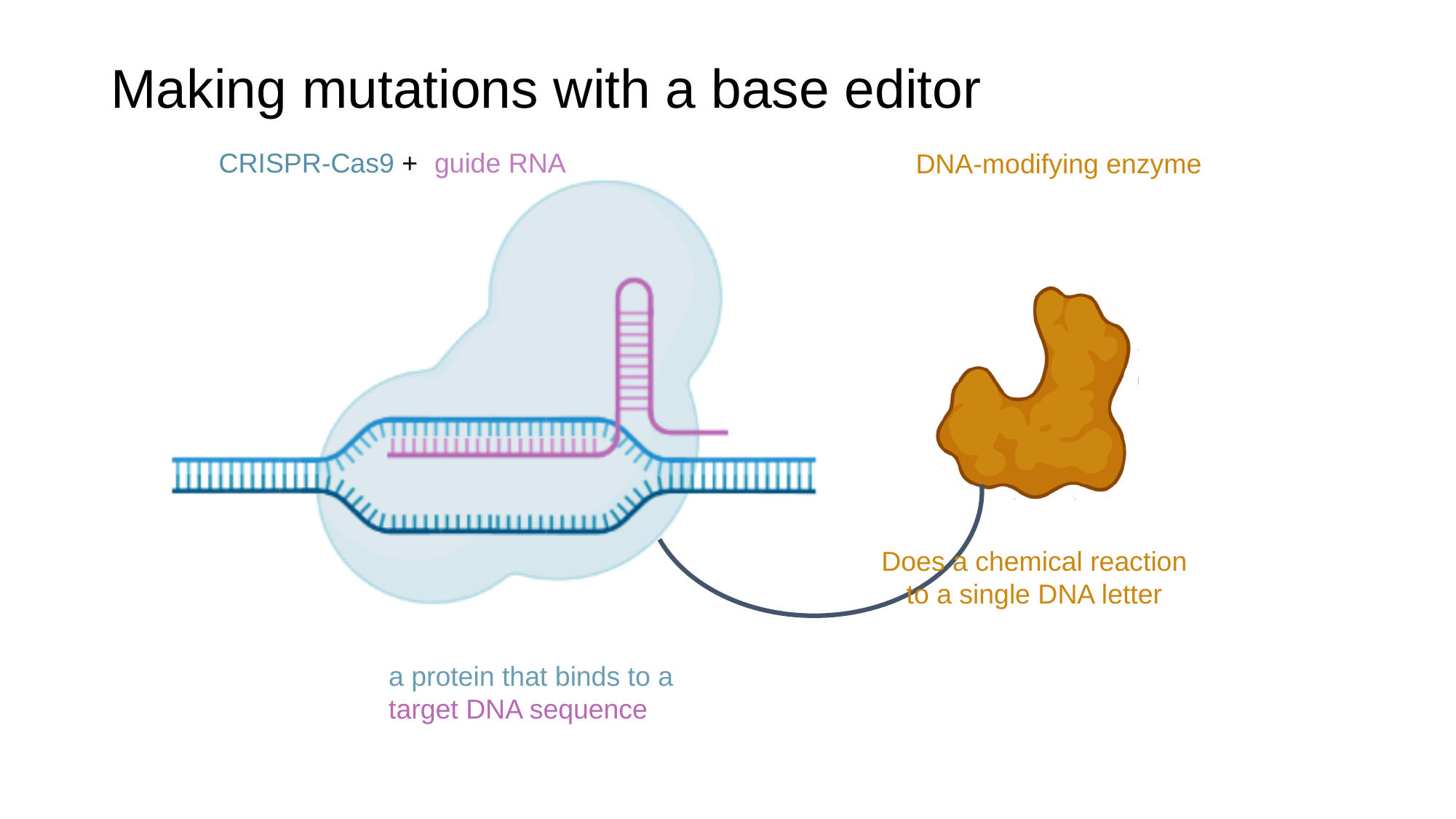

#### Making mutations with a base editor
CRISPR-Cas9 +
guide RNA
DNA-modifying enzyme
Does a chemical reaction to a single DNA letter
a protein that binds to a target DNA sequence

### Slide 22
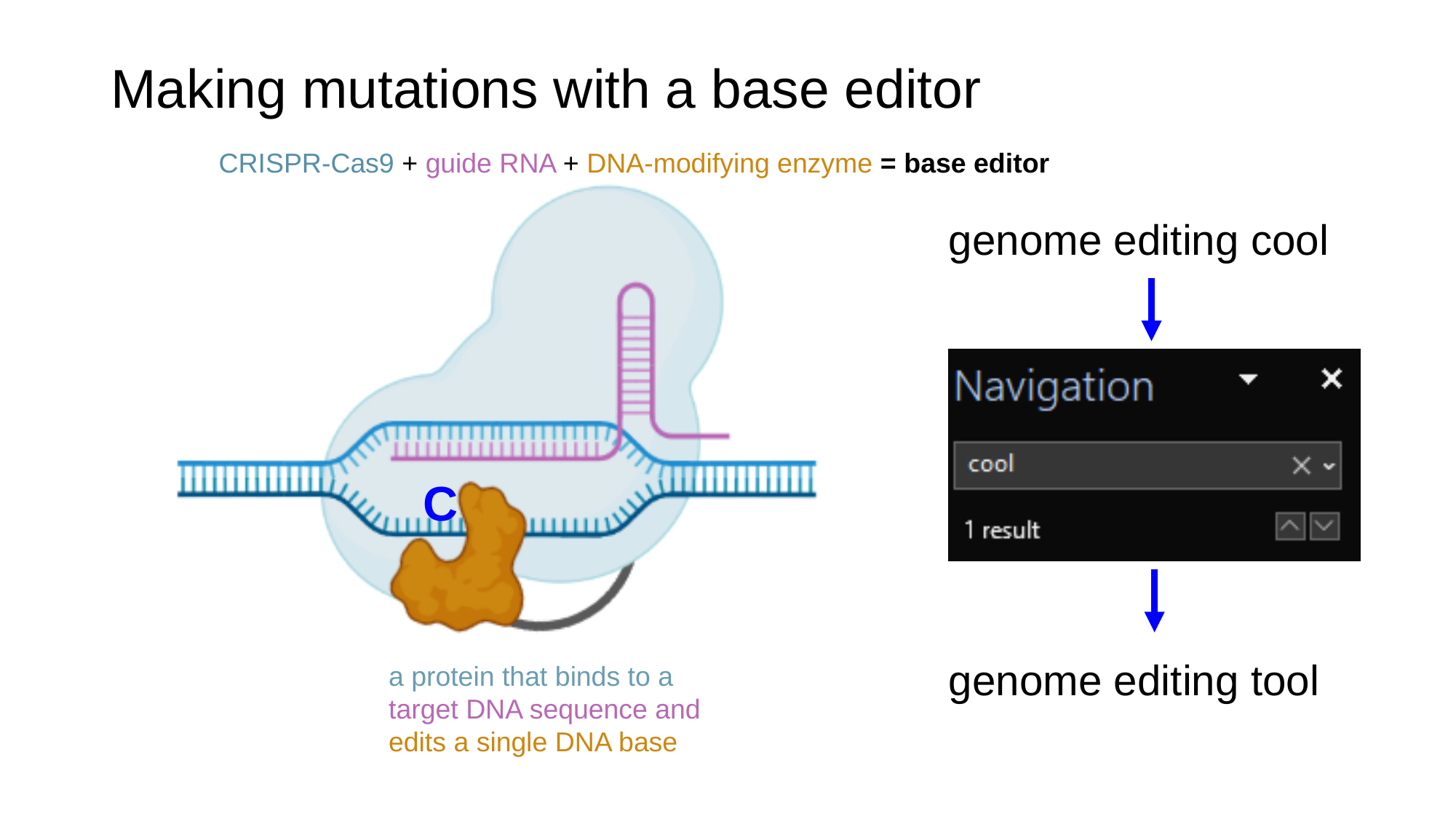

#### Making mutations with a base editor
CRISPR-Cas9 + guide RNA + DNA-modifying enzyme = base editor
C
genome editing cool
genome editing tool
a protein that binds to a target DNA sequence and
edits a single DNA base

### Slide 23
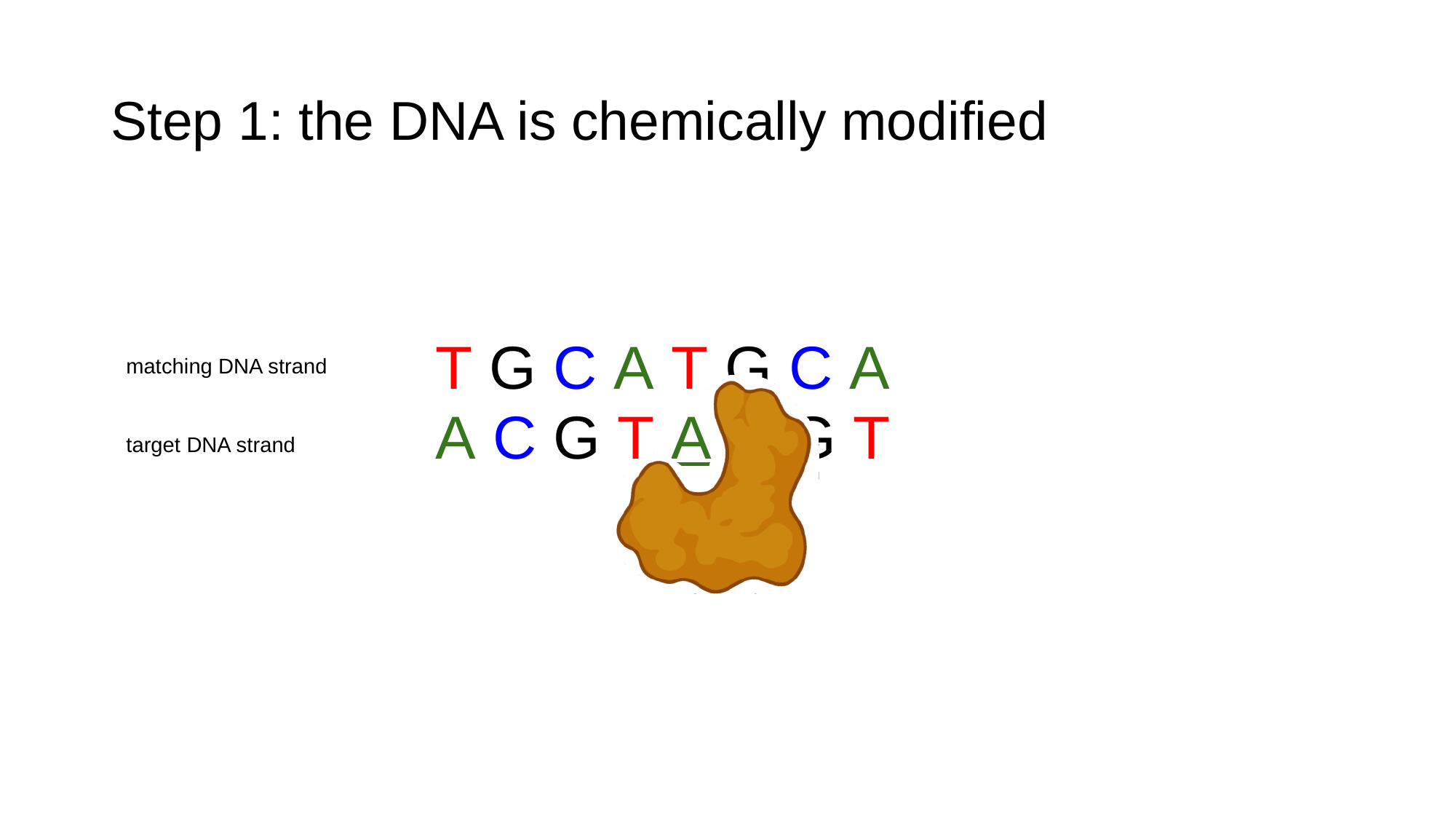

#### Step 1: the DNA is chemically modified
T G C A T G C A
matching DNA strand
A C G T A C G T
target DNA strand

### Slide 24
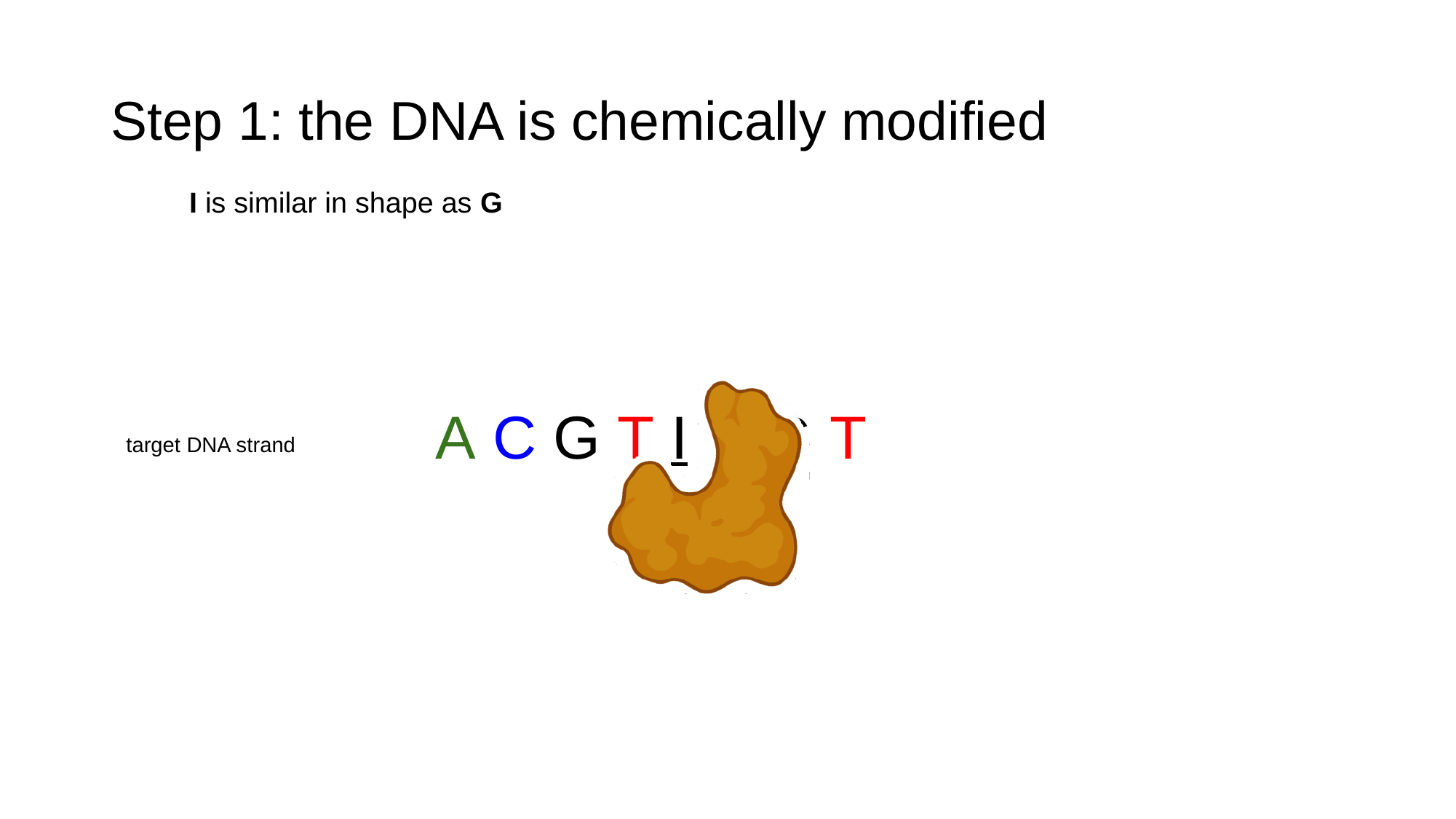

#### Step 1: the DNA is chemically modified
I is similar in shape as G
A C G T I C G T
target DNA strand

### Slide 25
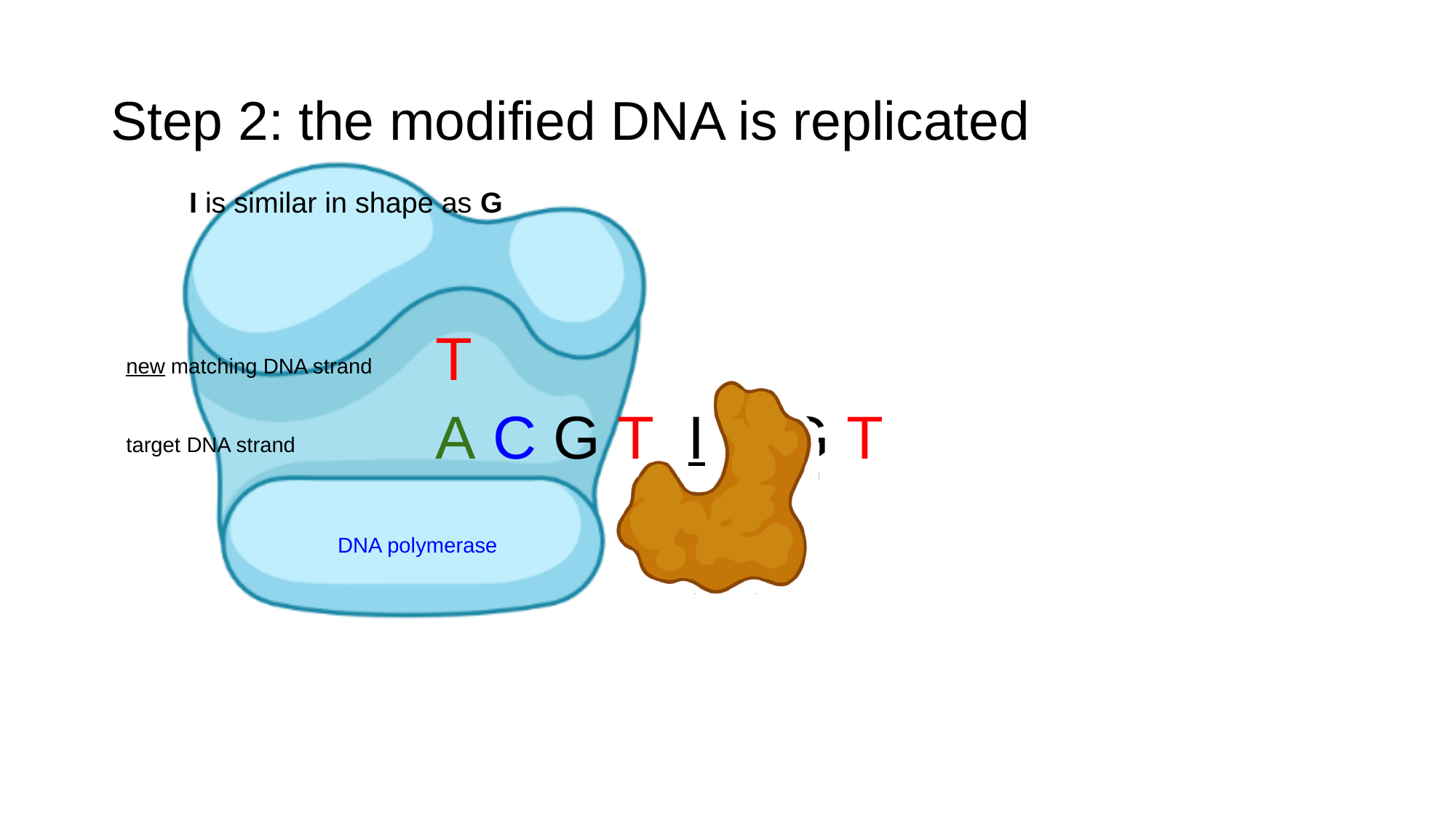

#### Step 2: the modified DNA is replicated
I is similar in shape as G
T
new matching DNA strand
A C G T I C G T
target DNA strand
DNA polymerase

### Slide 26
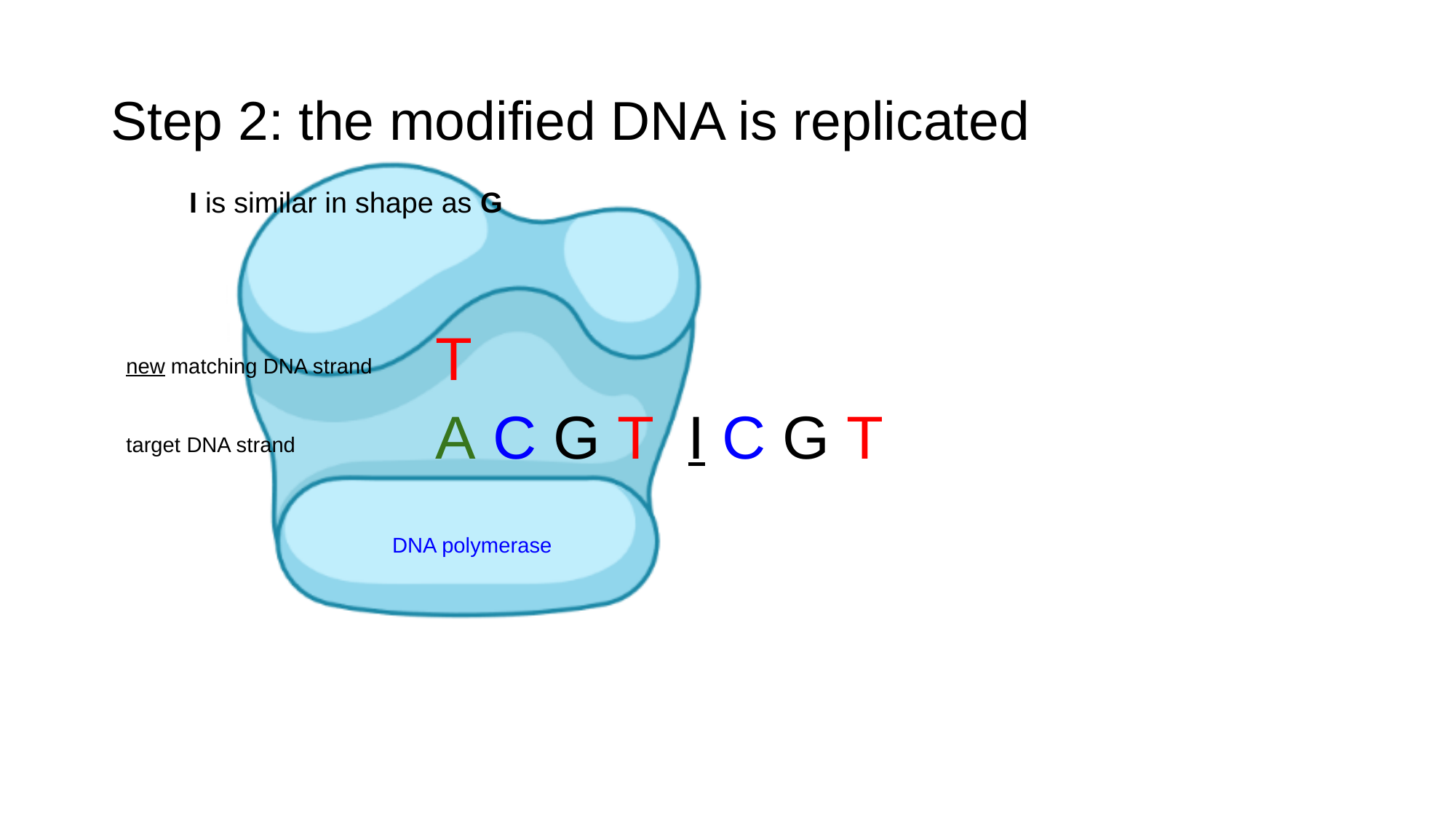

#### Step 2: the modified DNA is replicated
I is similar in shape as G
T
new matching DNA strand
A C G T I C G T
target DNA strand
DNA polymerase

### Slide 27
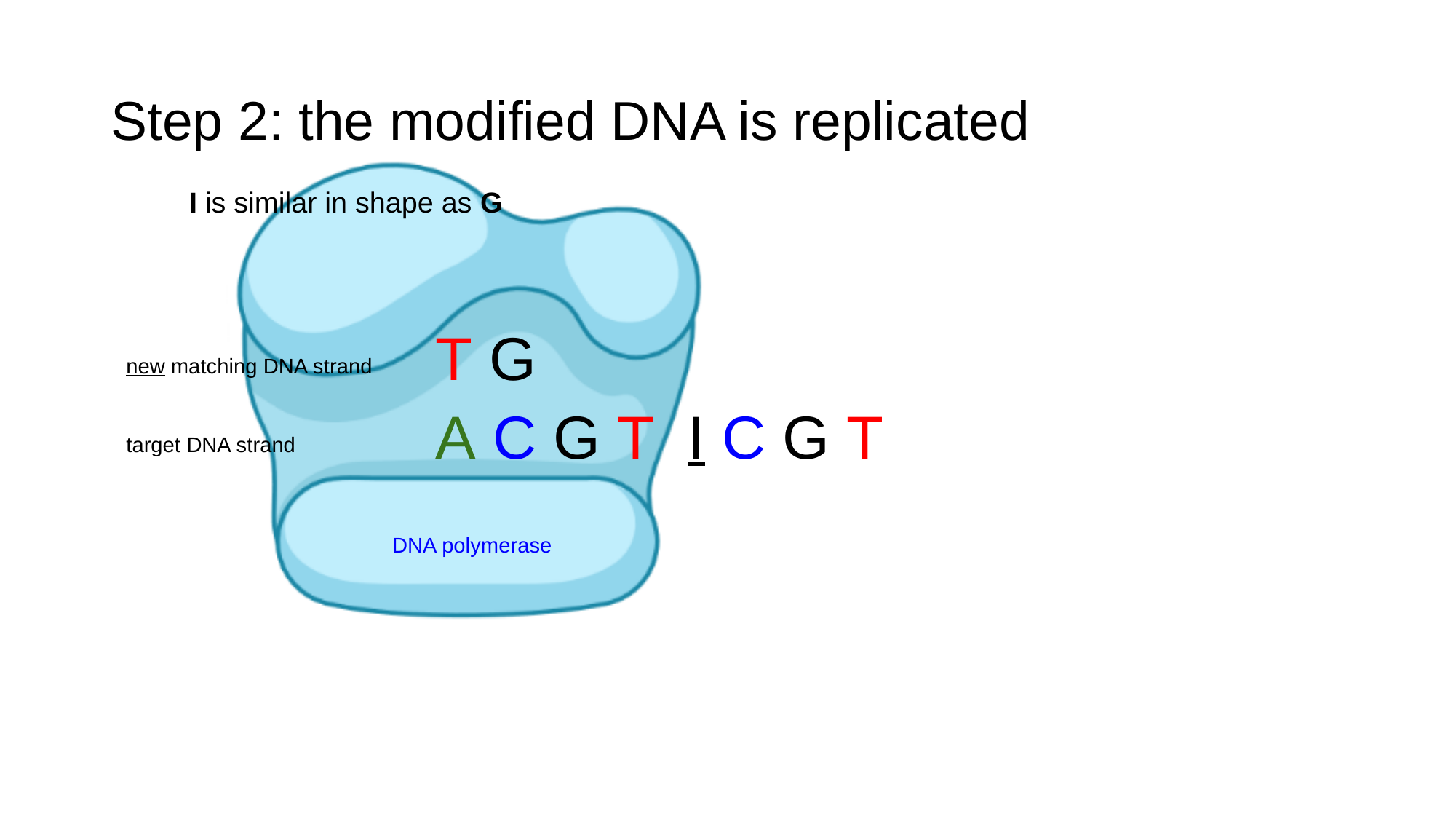

#### Step 2: the modified DNA is replicated
I is similar in shape as G
T G
new matching DNA strand
A C G T I C G T
target DNA strand
DNA polymerase

### Slide 28
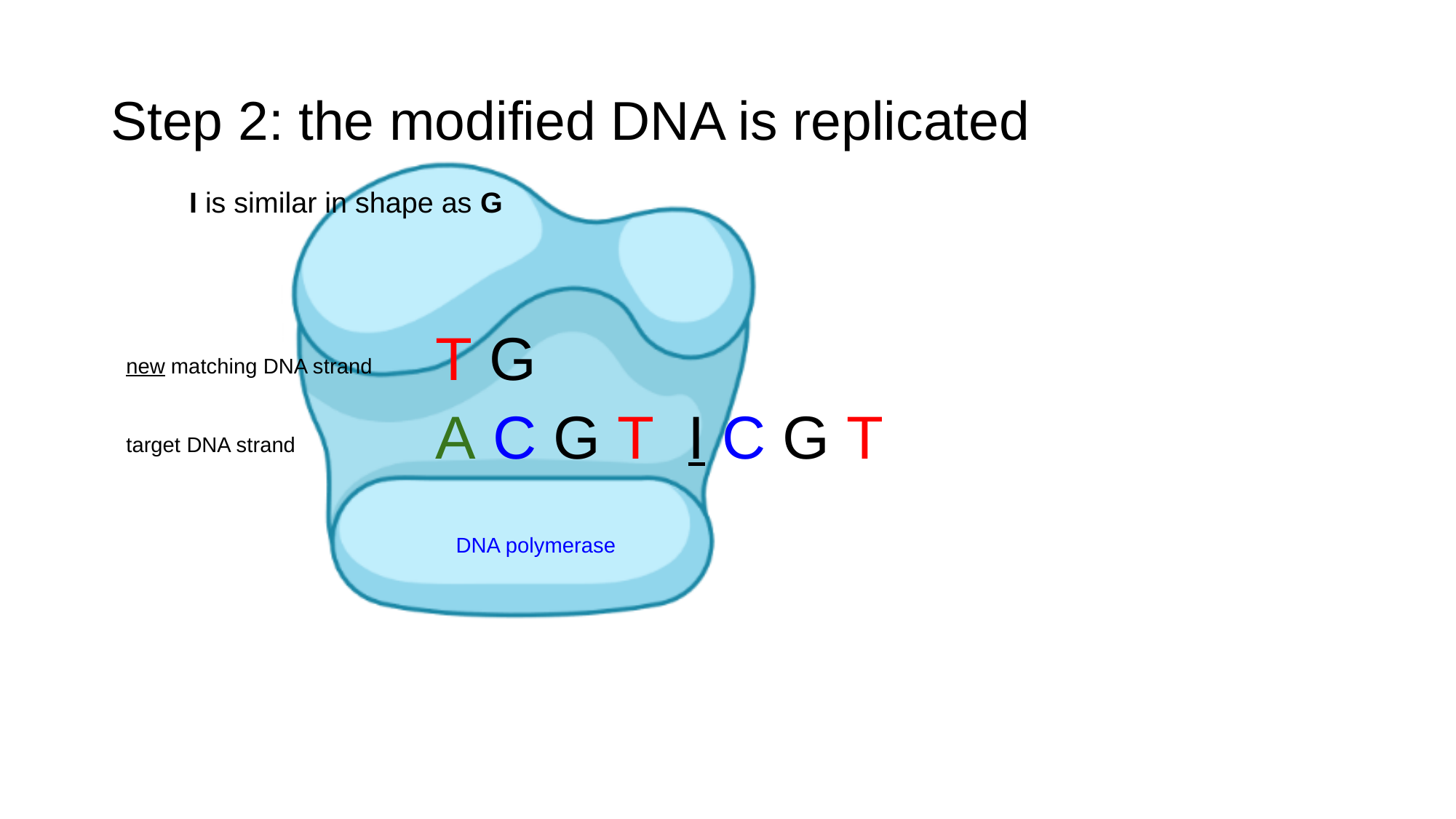

#### Step 2: the modified DNA is replicated
I is similar in shape as G
T G
new matching DNA strand
A C G T I C G T
target DNA strand
DNA polymerase

### Slide 29
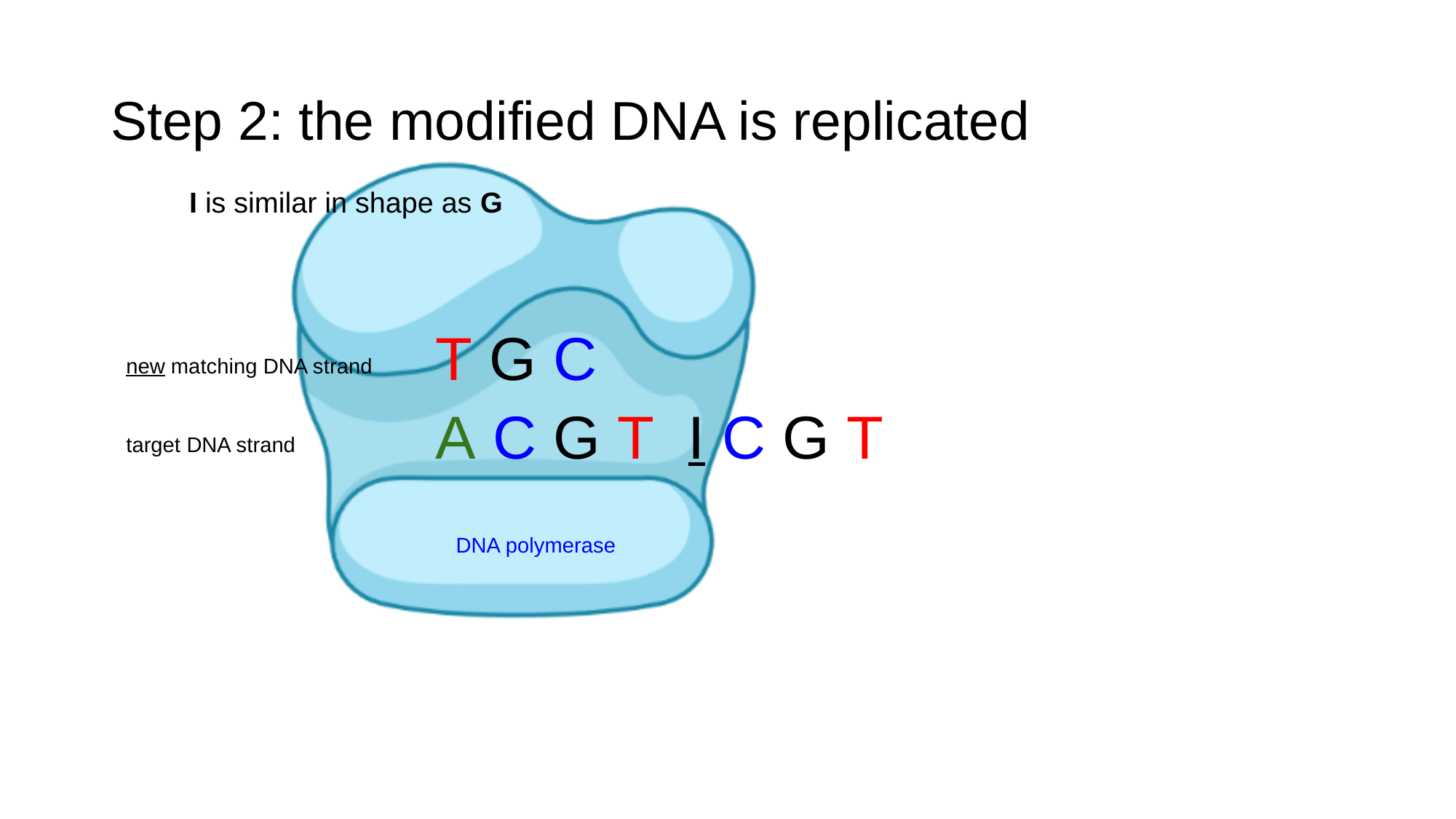

#### Step 2: the modified DNA is replicated
I is similar in shape as G
T G C
new matching DNA strand
A C G T I C G T
target DNA strand
DNA polymerase

### Slide 30
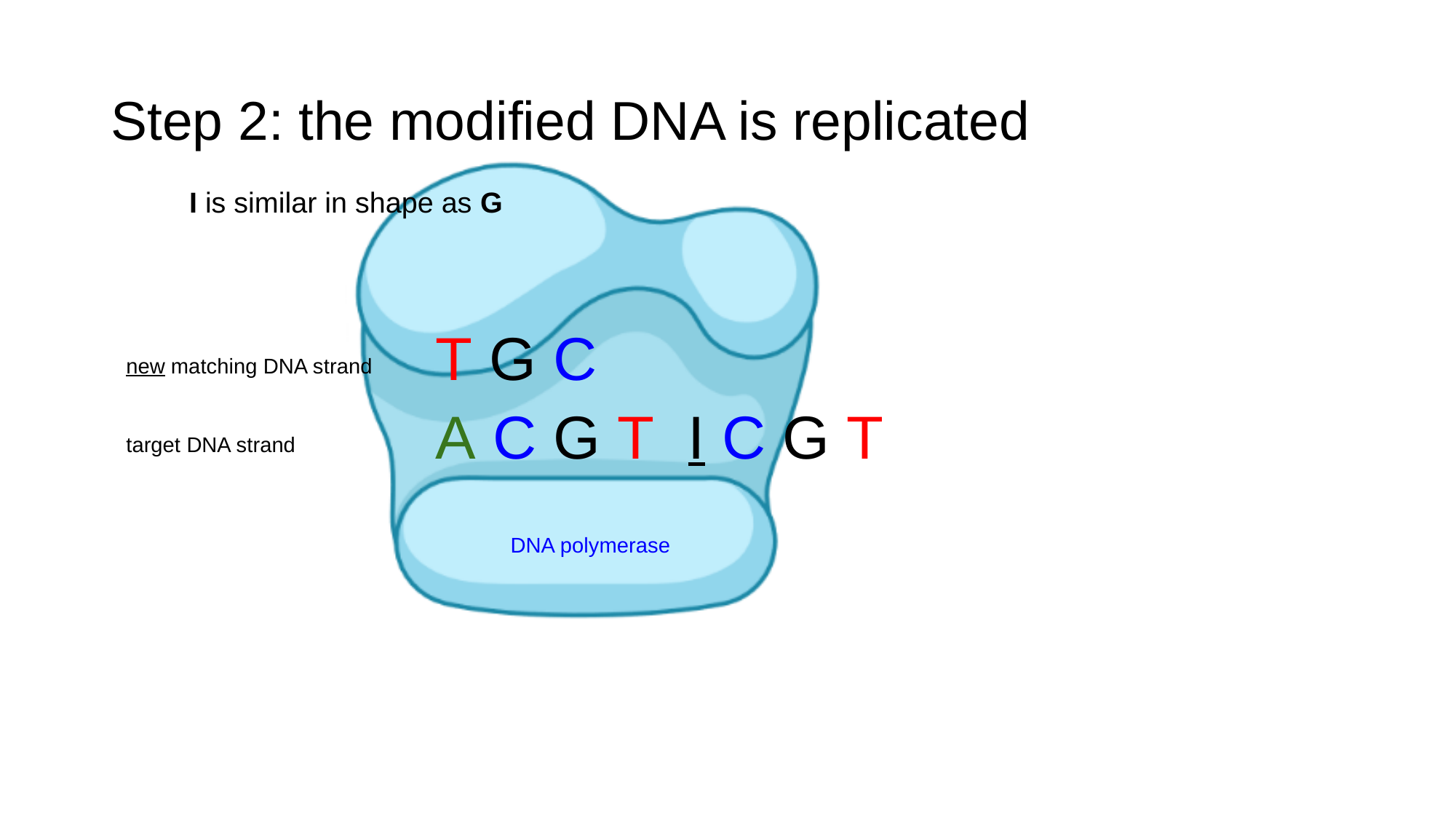

#### Step 2: the modified DNA is replicated
I is similar in shape as G
T G C
new matching DNA strand
A C G T I C G T
target DNA strand
DNA polymerase

### Slide 31
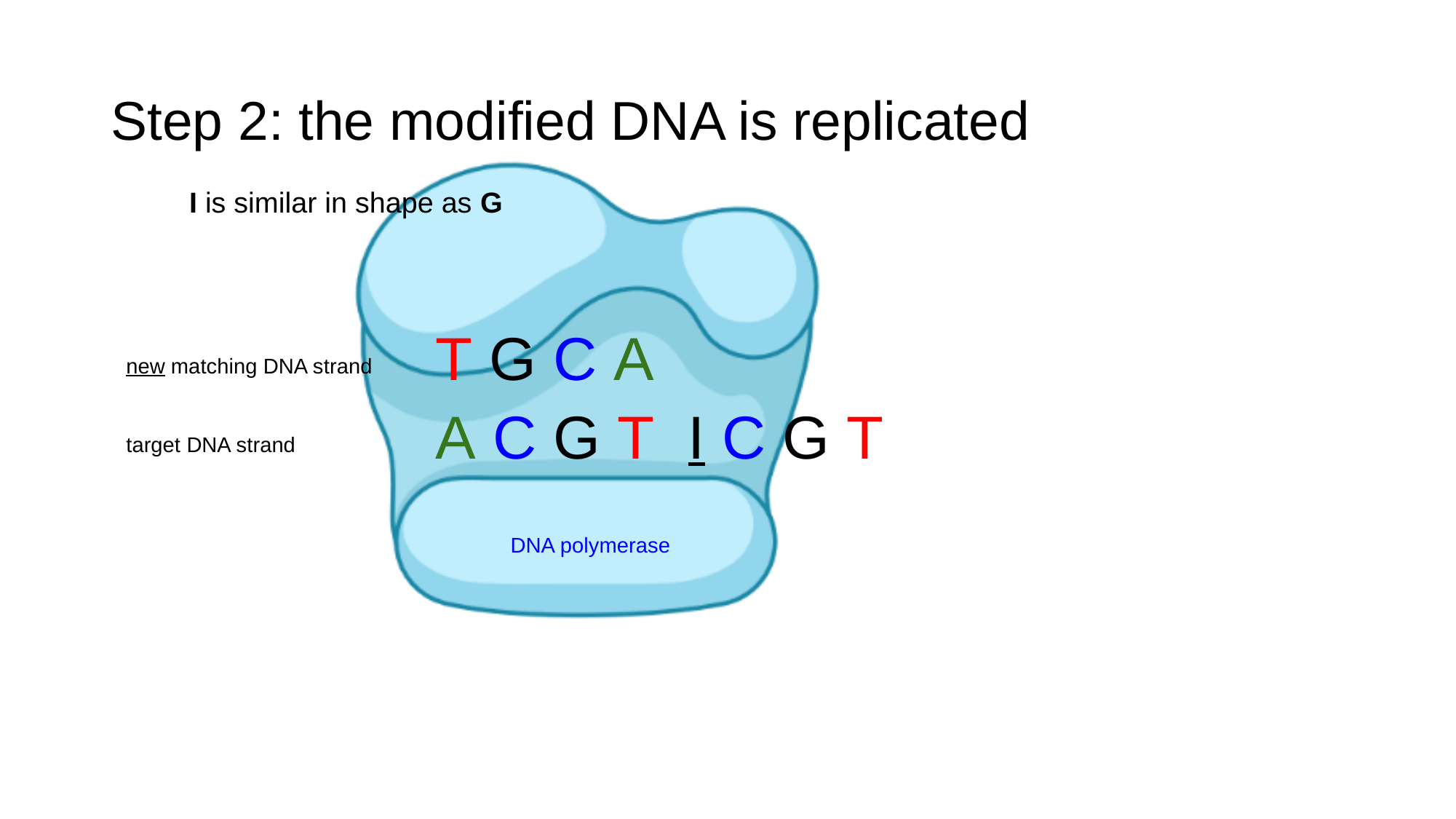

#### Step 2: the modified DNA is replicated
I is similar in shape as G
T G C A
new matching DNA strand
A C G T I C G T
target DNA strand
DNA polymerase

### Slide 32
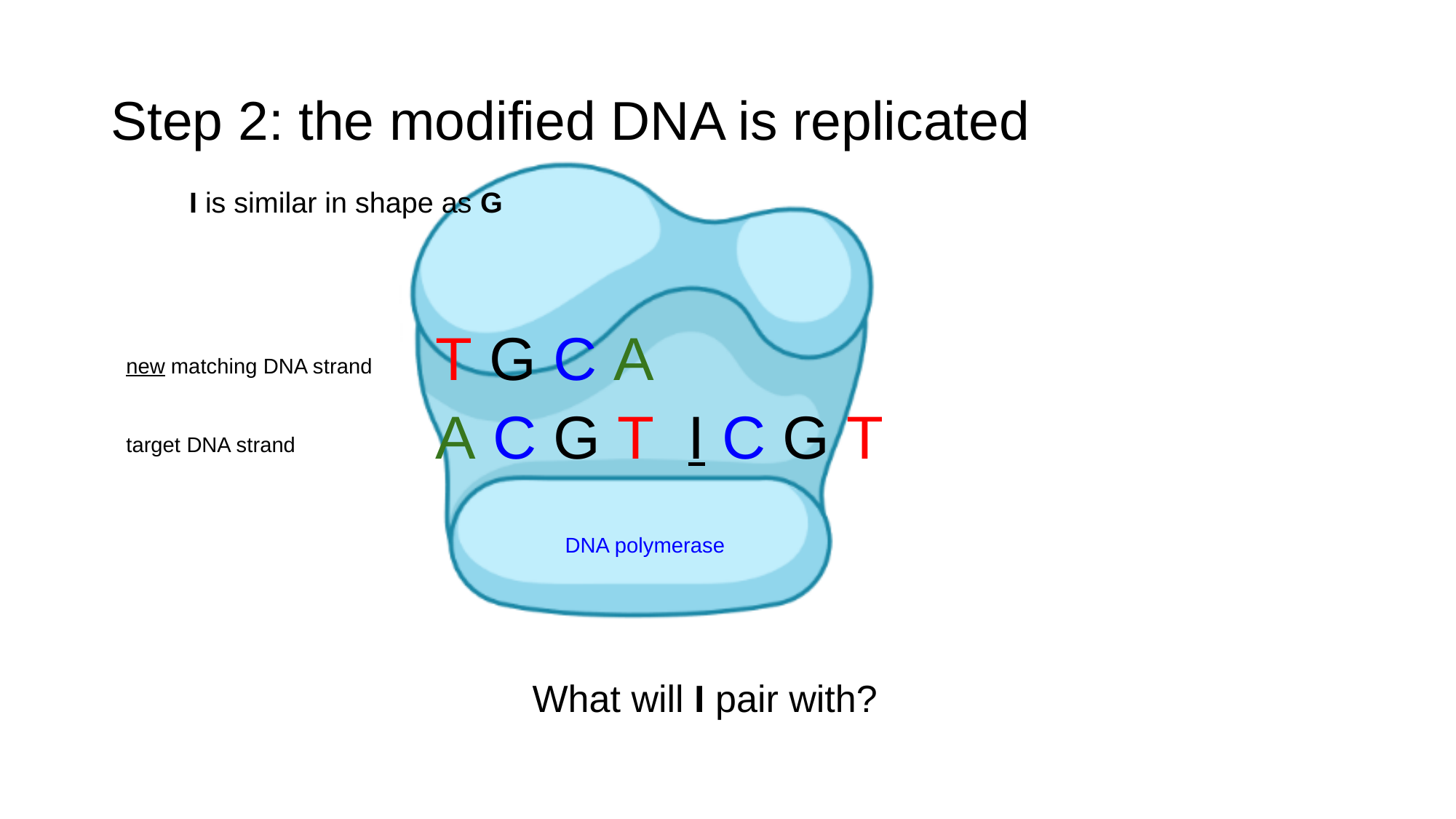

#### Step 2: the modified DNA is replicated
I is similar in shape as G
T G C A
new matching DNA strand
A C G T I C G T
target DNA strand
DNA polymerase
What will I pair with?

### Slide 33
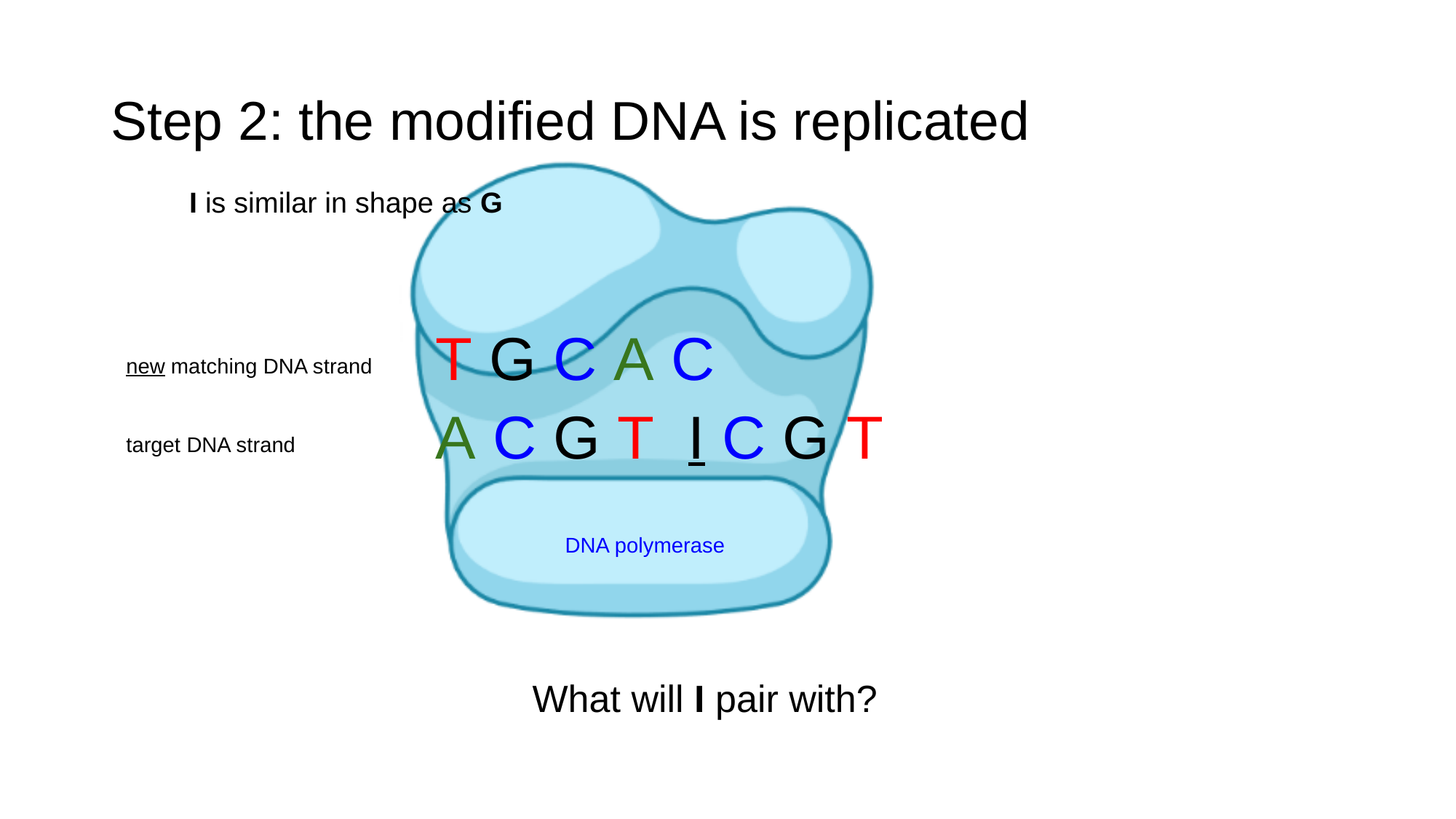

#### Step 2: the modified DNA is replicated
I is similar in shape as G
T G C A C
new matching DNA strand
A C G T I C G T
target DNA strand
DNA polymerase
What will I pair with?

### Slide 34
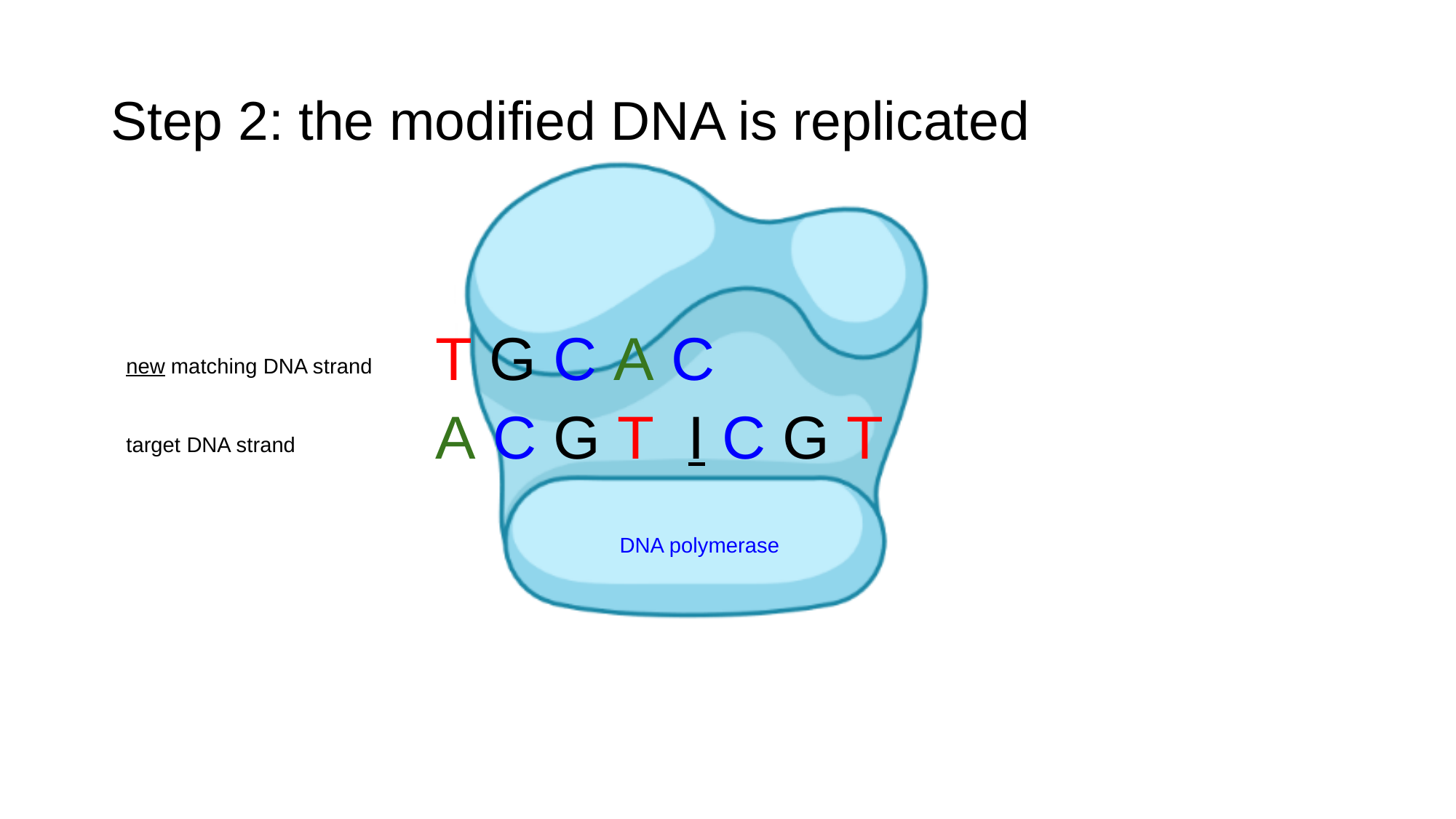

#### Step 2: the modified DNA is replicated
T G C A C
new matching DNA strand
A C G T I C G T
target DNA strand
DNA polymerase

### Slide 35
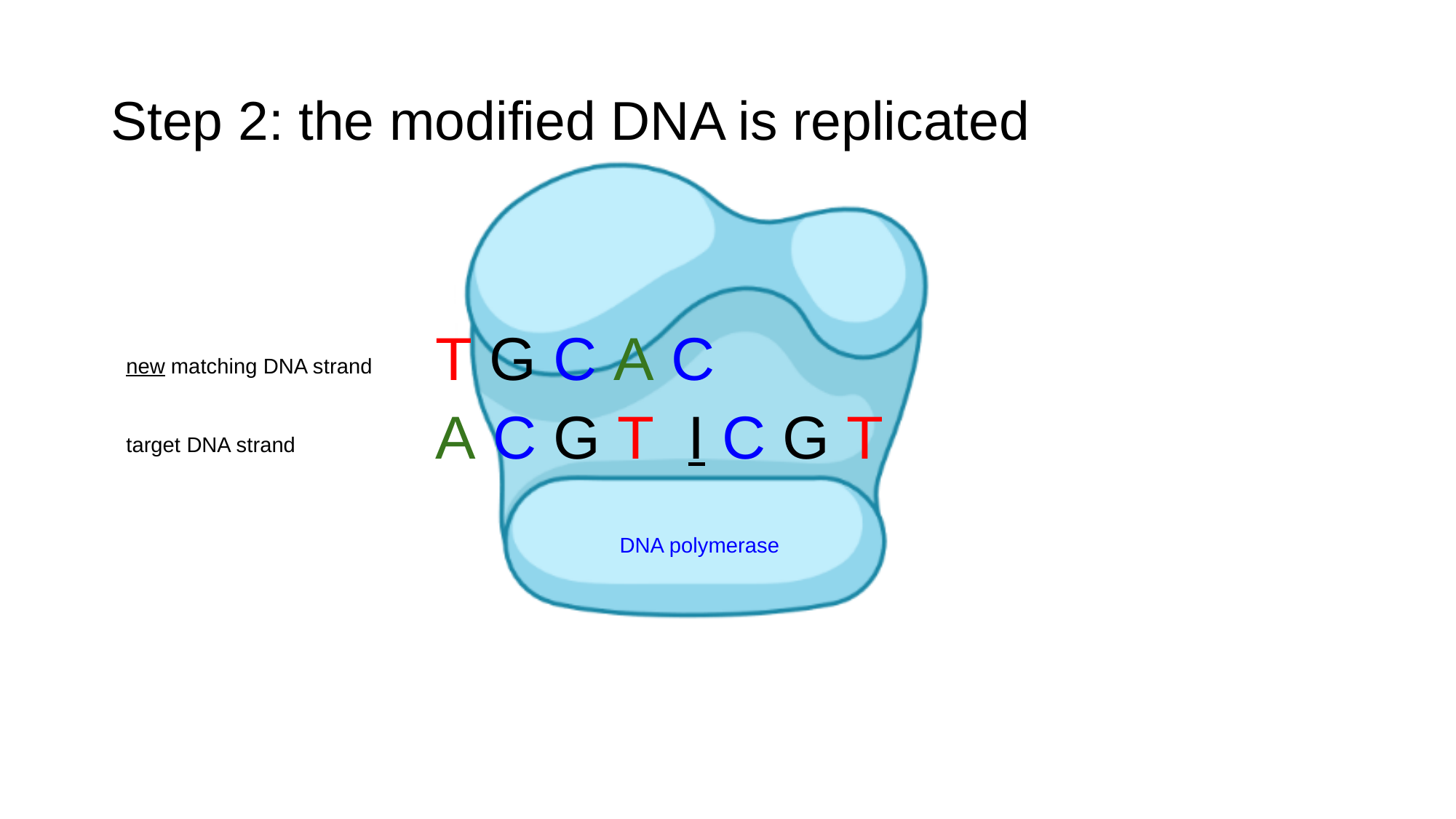

#### Step 2: the modified DNA is replicated
T G C A C
new matching DNA strand
A C G T I C G T
target DNA strand
DNA polymerase

### Slide 36
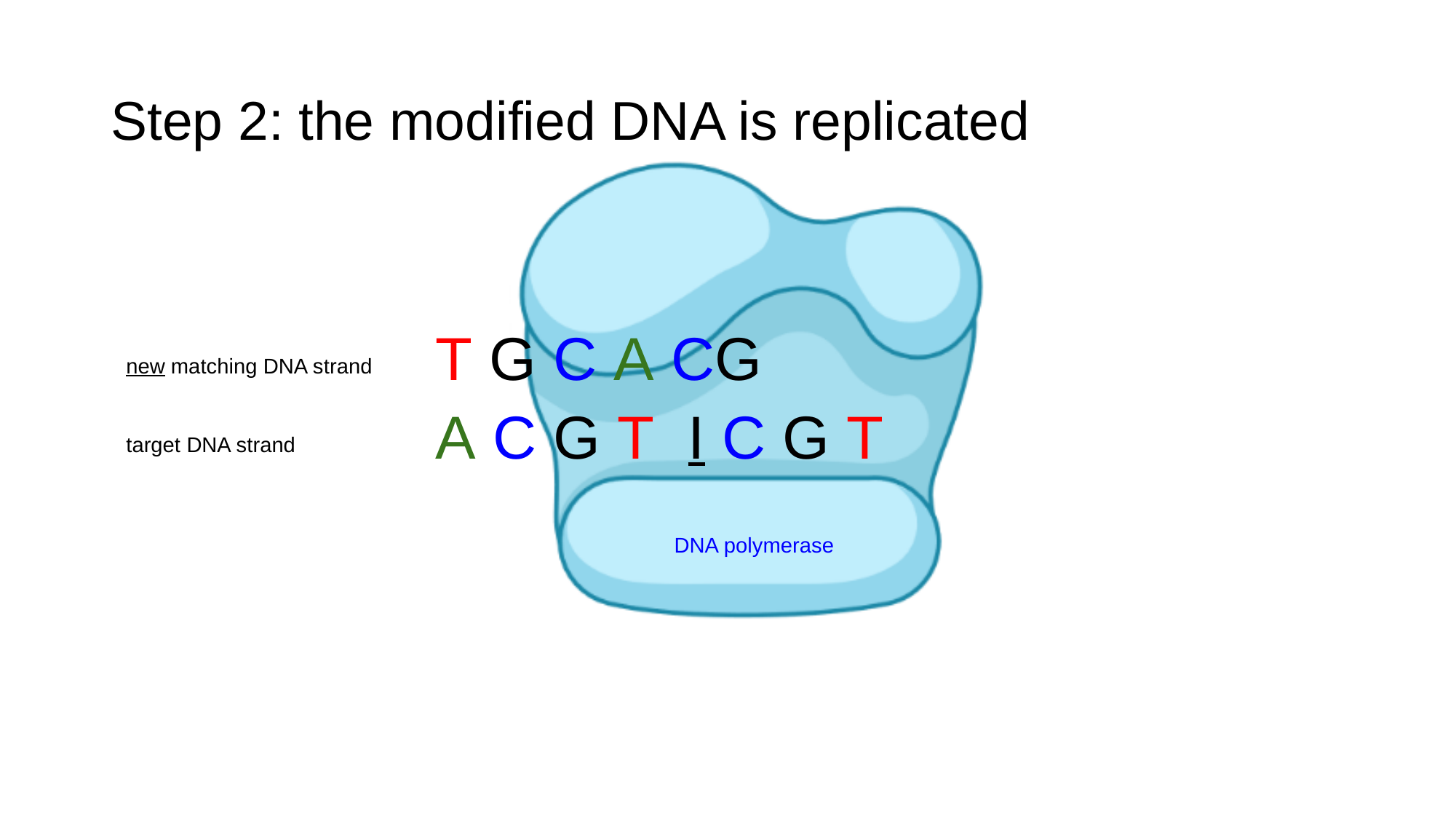

#### Step 2: the modified DNA is replicated
T G C A CG
new matching DNA strand
A C G T I C G T
target DNA strand
DNA polymerase

### Slide 37
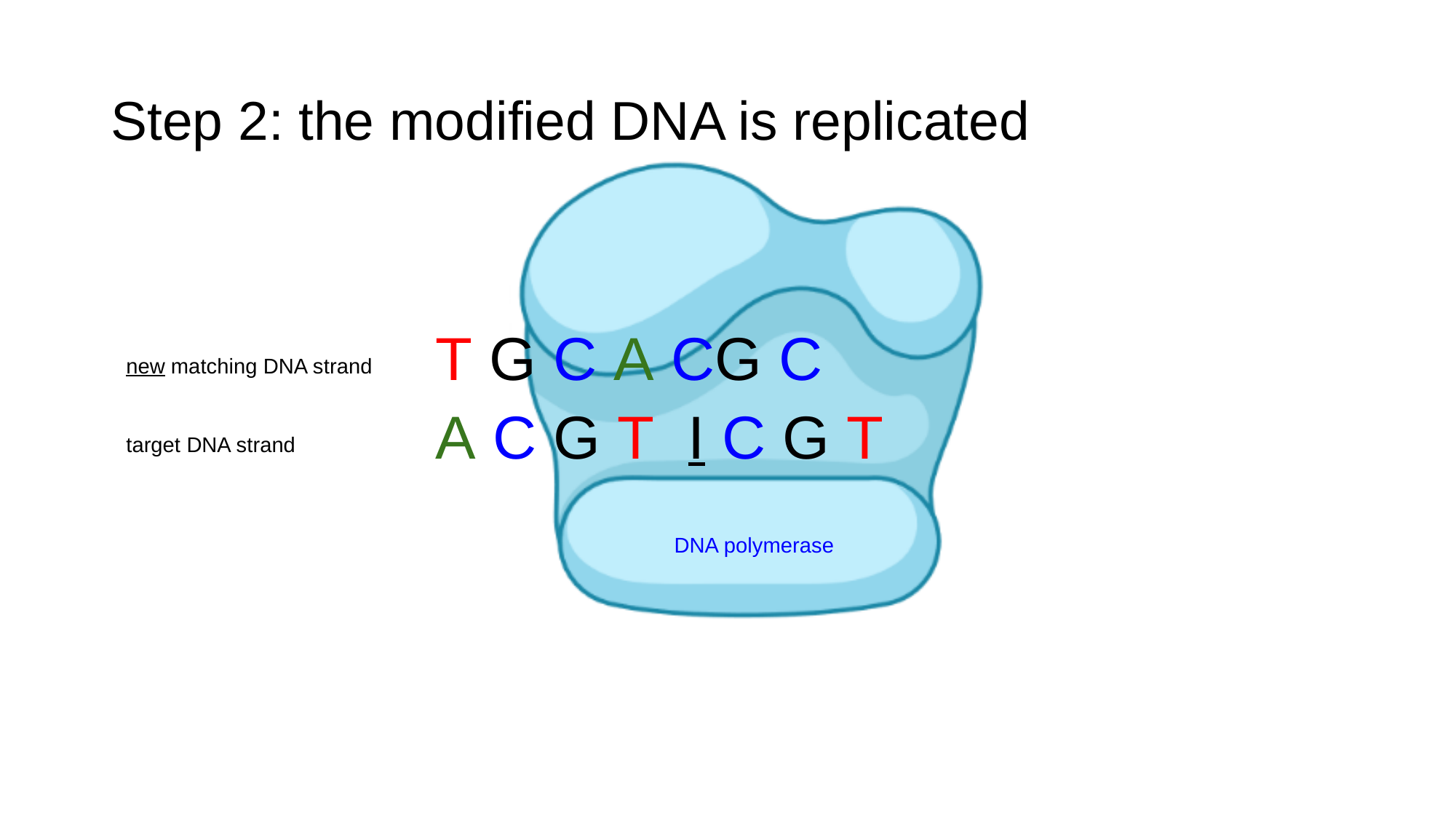

#### Step 2: the modified DNA is replicated
T G C A CG C
new matching DNA strand
A C G T I C G T
target DNA strand
DNA polymerase

### Slide 38
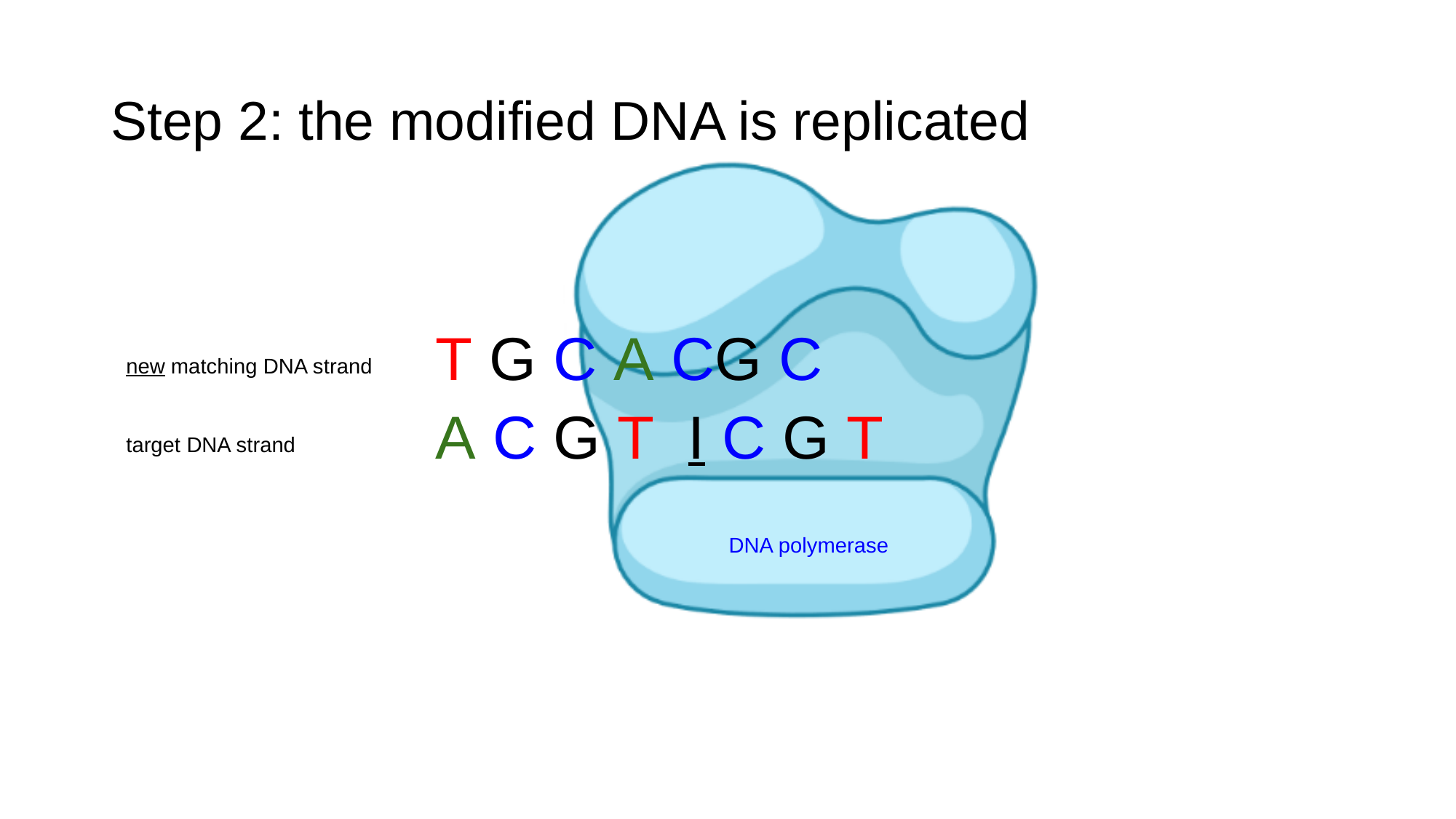

#### Step 2: the modified DNA is replicated
T G C A CG C
new matching DNA strand
A C G T I C G T
target DNA strand
DNA polymerase

### Slide 39
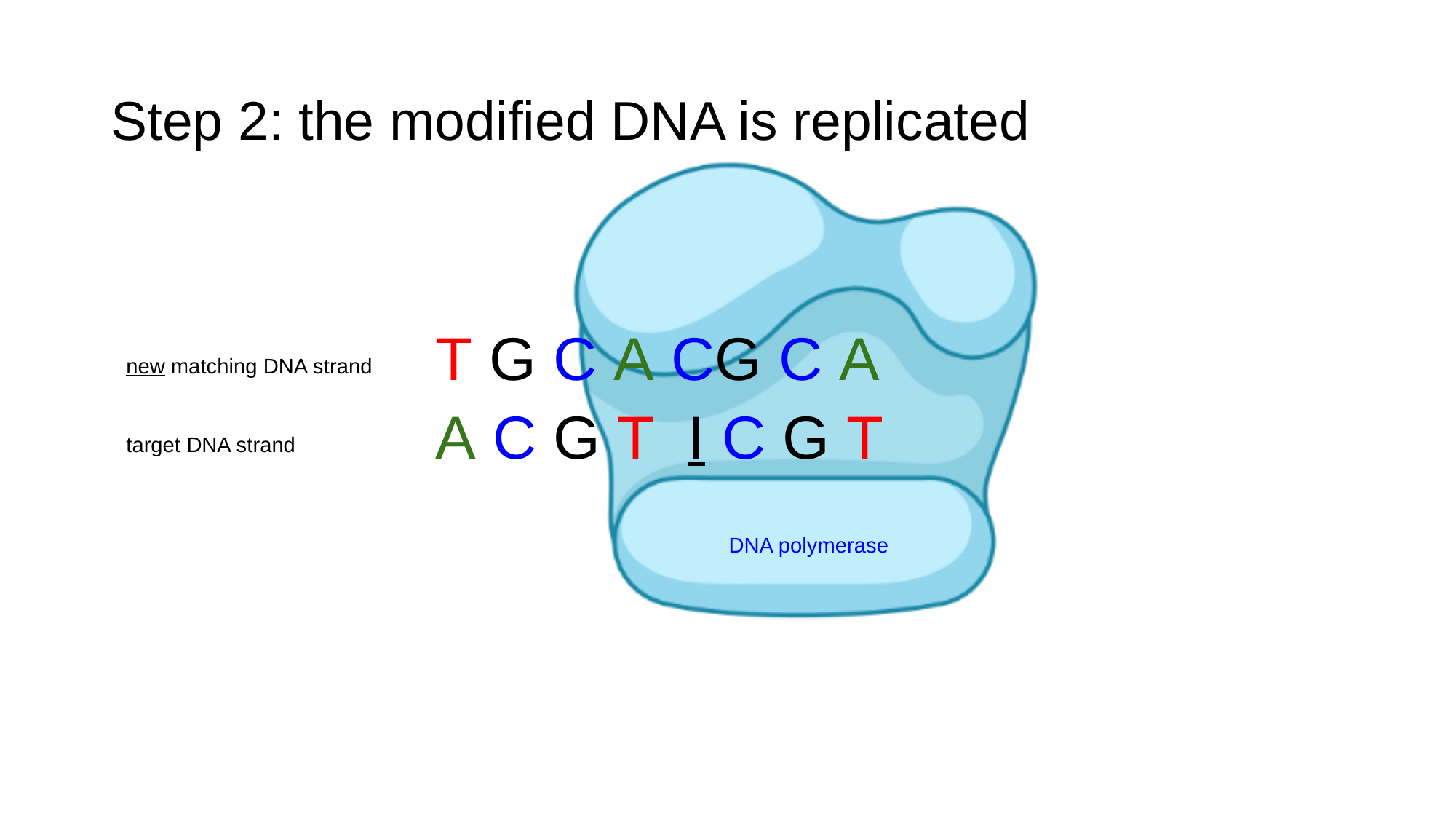

#### Step 2: the modified DNA is replicated
T G C A CG C A
new matching DNA strand
A C G T I C G T
target DNA strand
DNA polymerase

### Slide 40
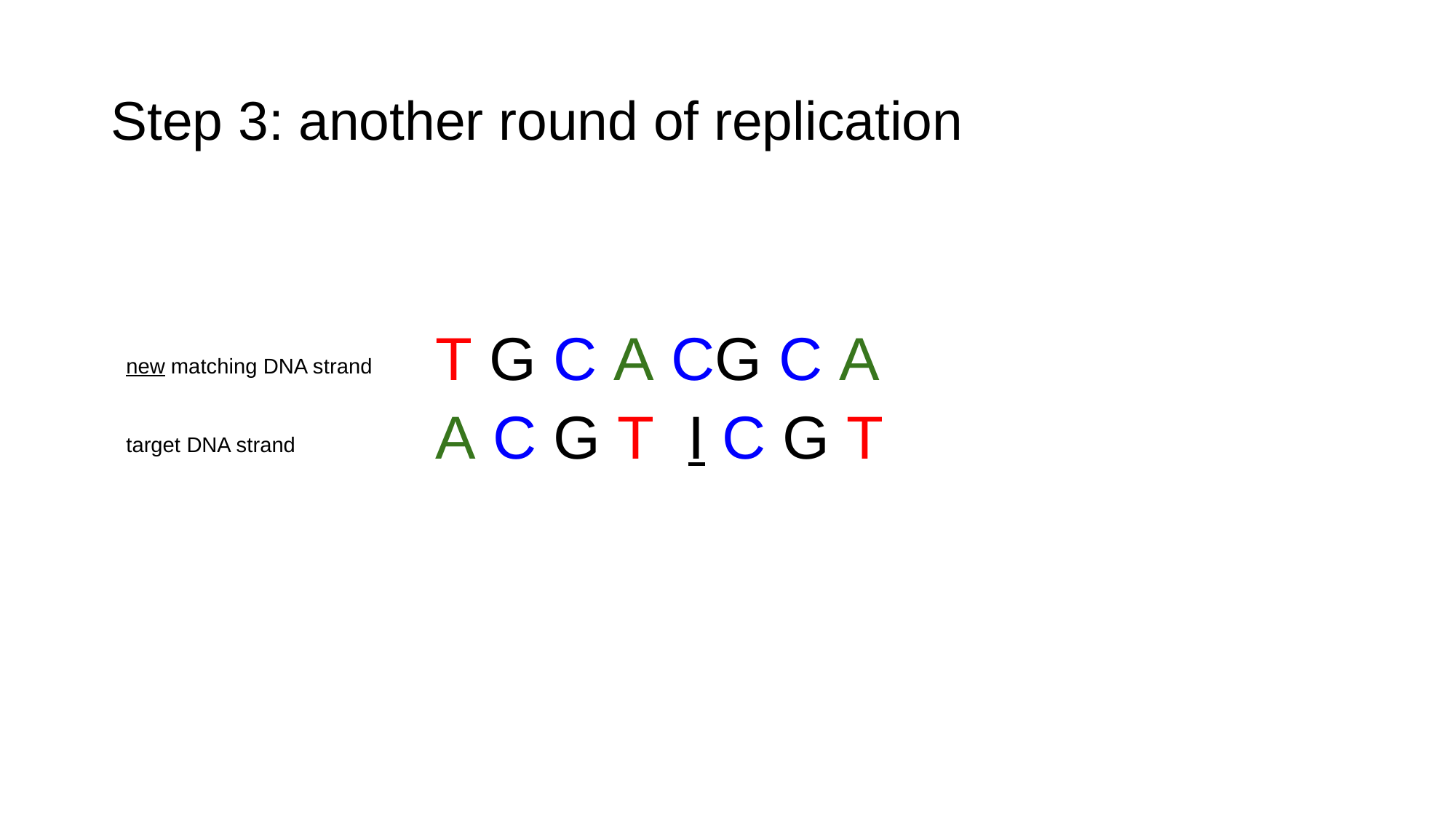

#### Step 3: another round of replication
T G C A CG C A
new matching DNA strand
A C G T I C G T
target DNA strand

### Slide 41
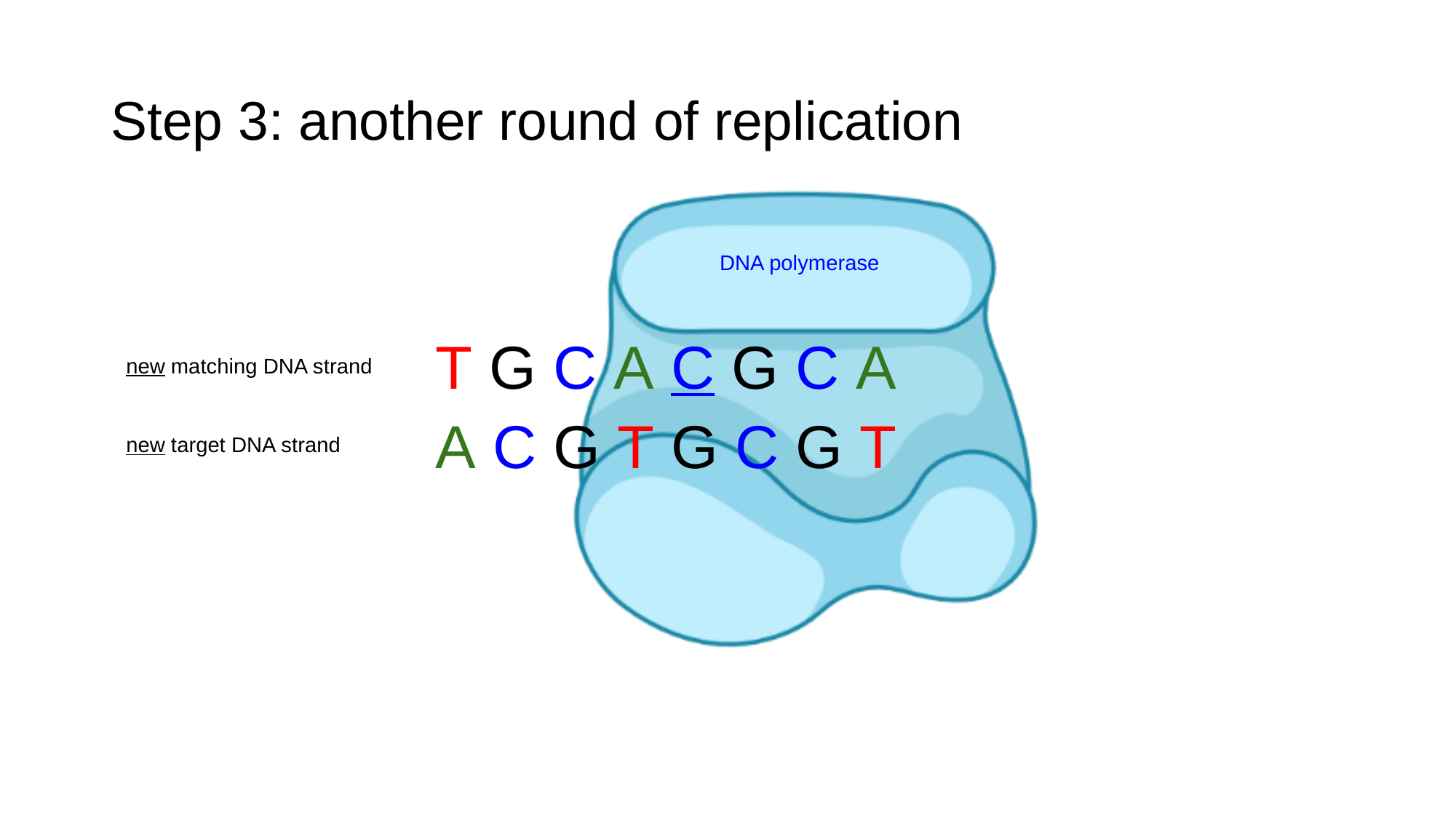

#### Step 3: another round of replication
DNA polymerase
T G C A C G C A
new matching DNA strand
A C G T G C G T
new target DNA strand

### Slide 42
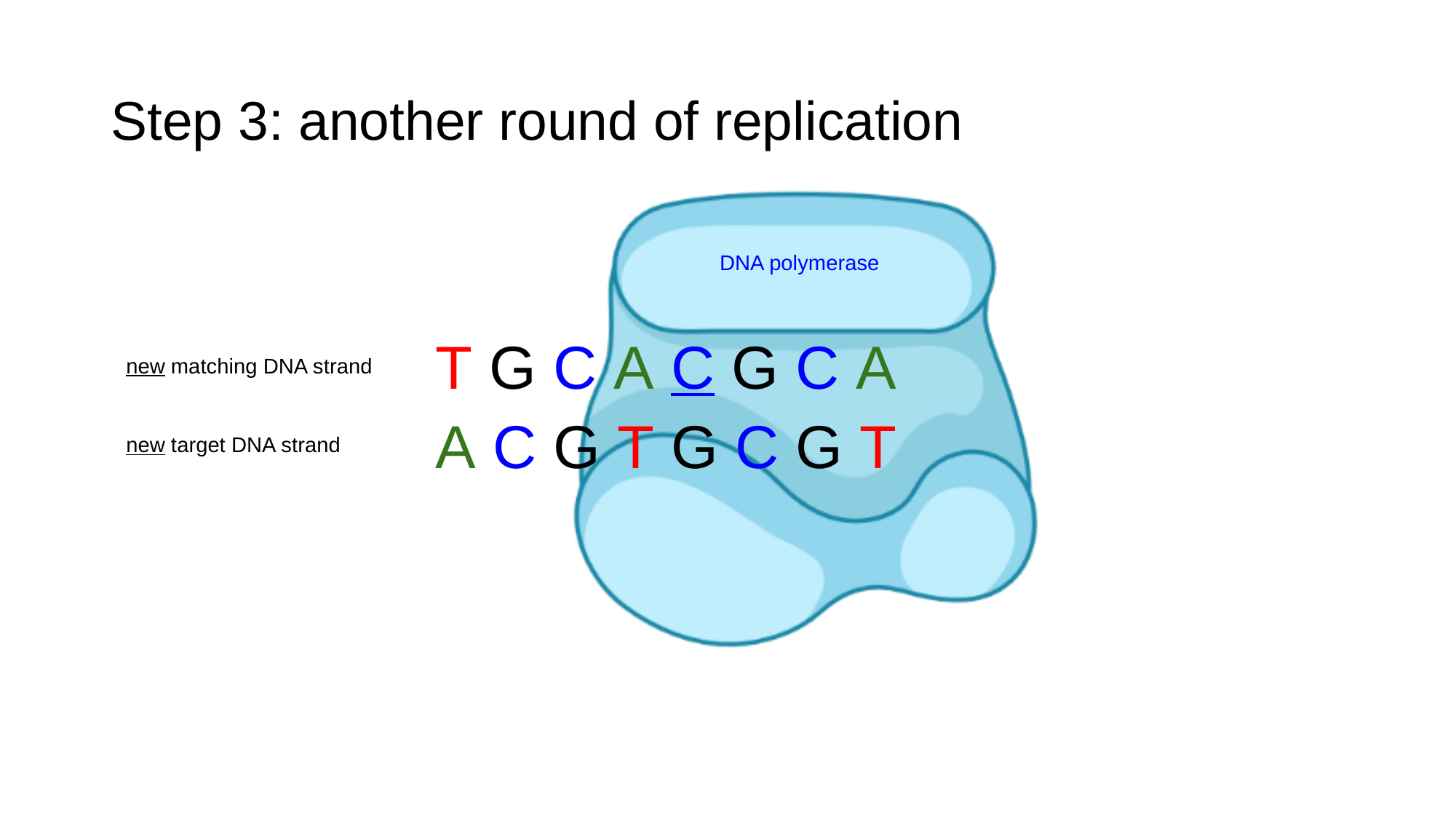

#### Step 3: another round of replication
DNA polymerase
T G C A C G C A
new matching DNA strand
A C G T G C G T
new target DNA strand

### Slide 43
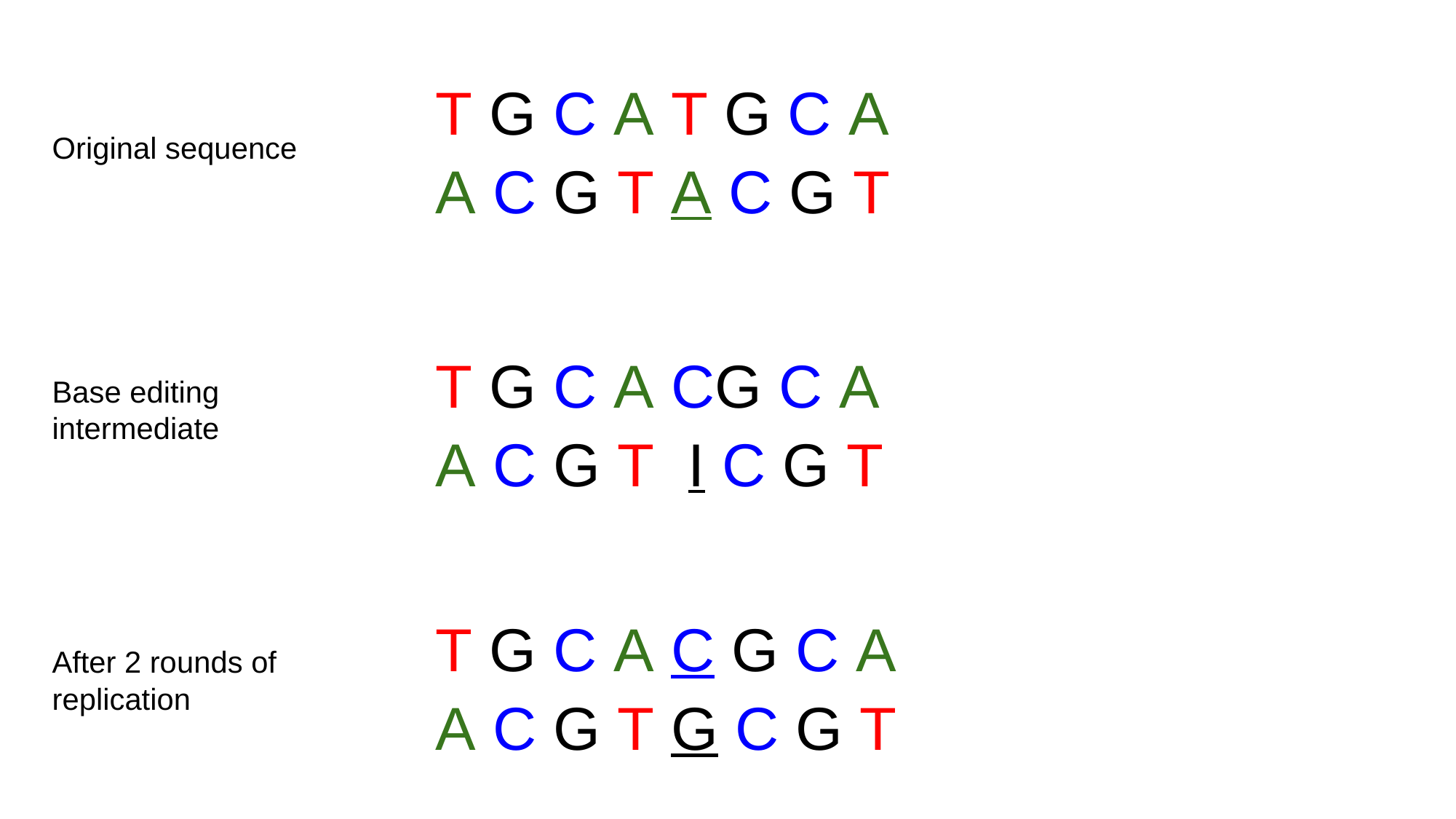

T G C A T G C A
Original sequence
A C G T A C G T
T G C A CG C A
Base editing intermediate
A C G T I C G T
T G C A C G C A
After 2 rounds of replication
A C G T G C G T

### Slide 44
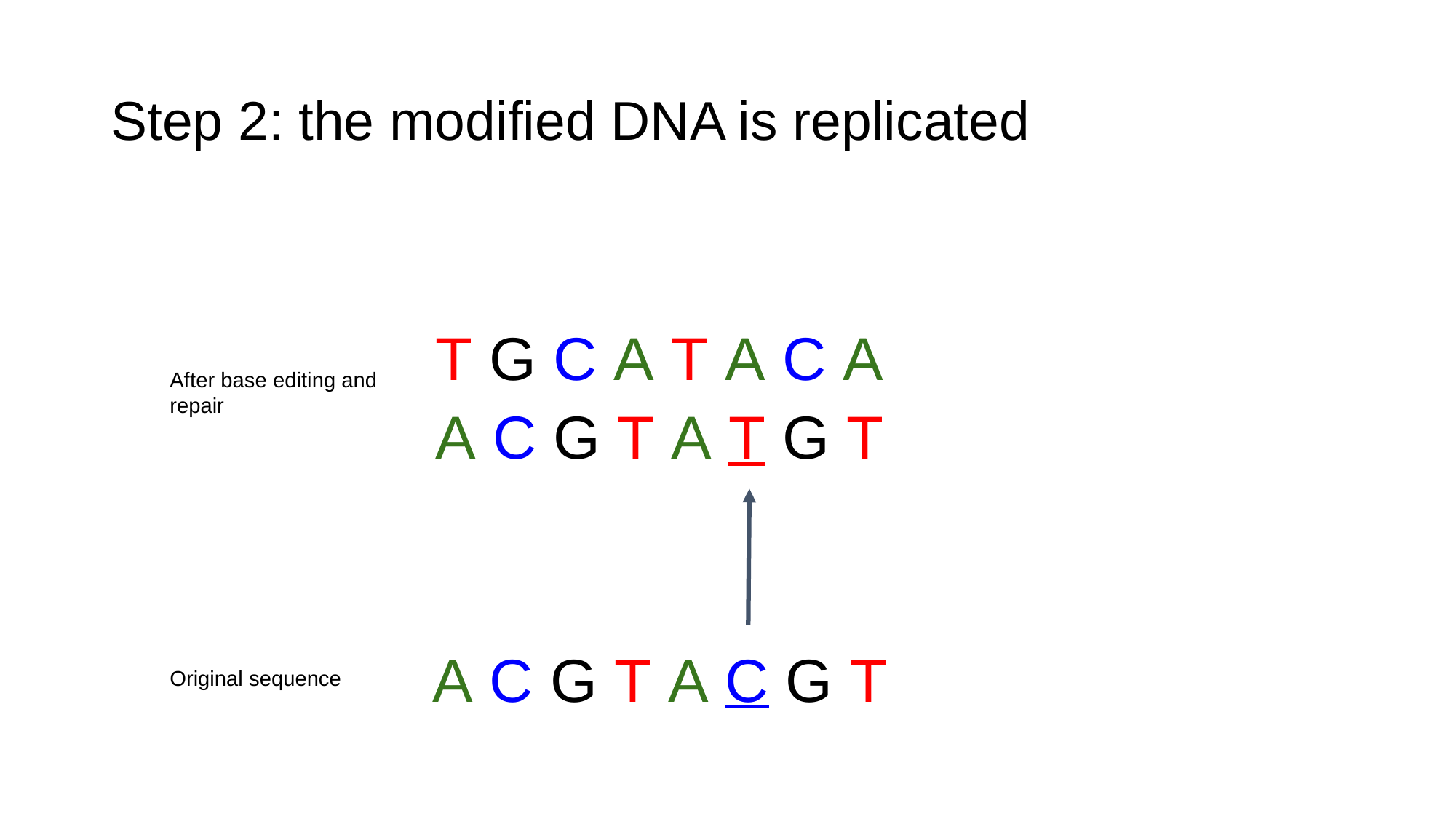

#### Step 2: the modified DNA is replicated
T G C A T A C A
After base editing and repair
A C G T A T G T
A C G T A C G T
Original sequence

### Slide 45
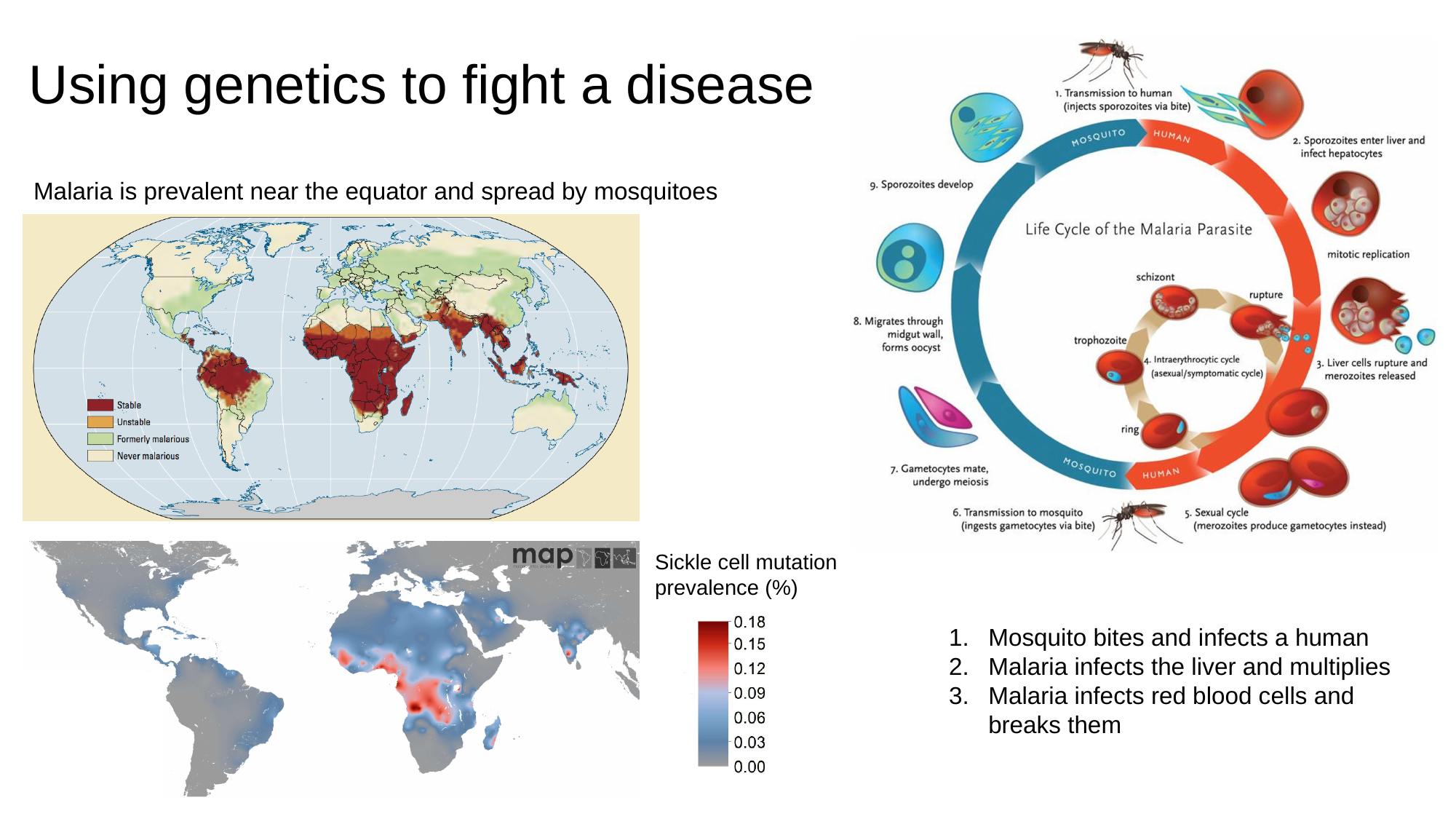

#### Using genetics to fight a disease
Malaria is prevalent near the equator and spread by mosquitoes
Sickle cell mutation prevalence (%)
Mosquito bites and infects a human
Malaria infects the liver and multiplies
Malaria infects red blood cells and breaks them

### Slide 46
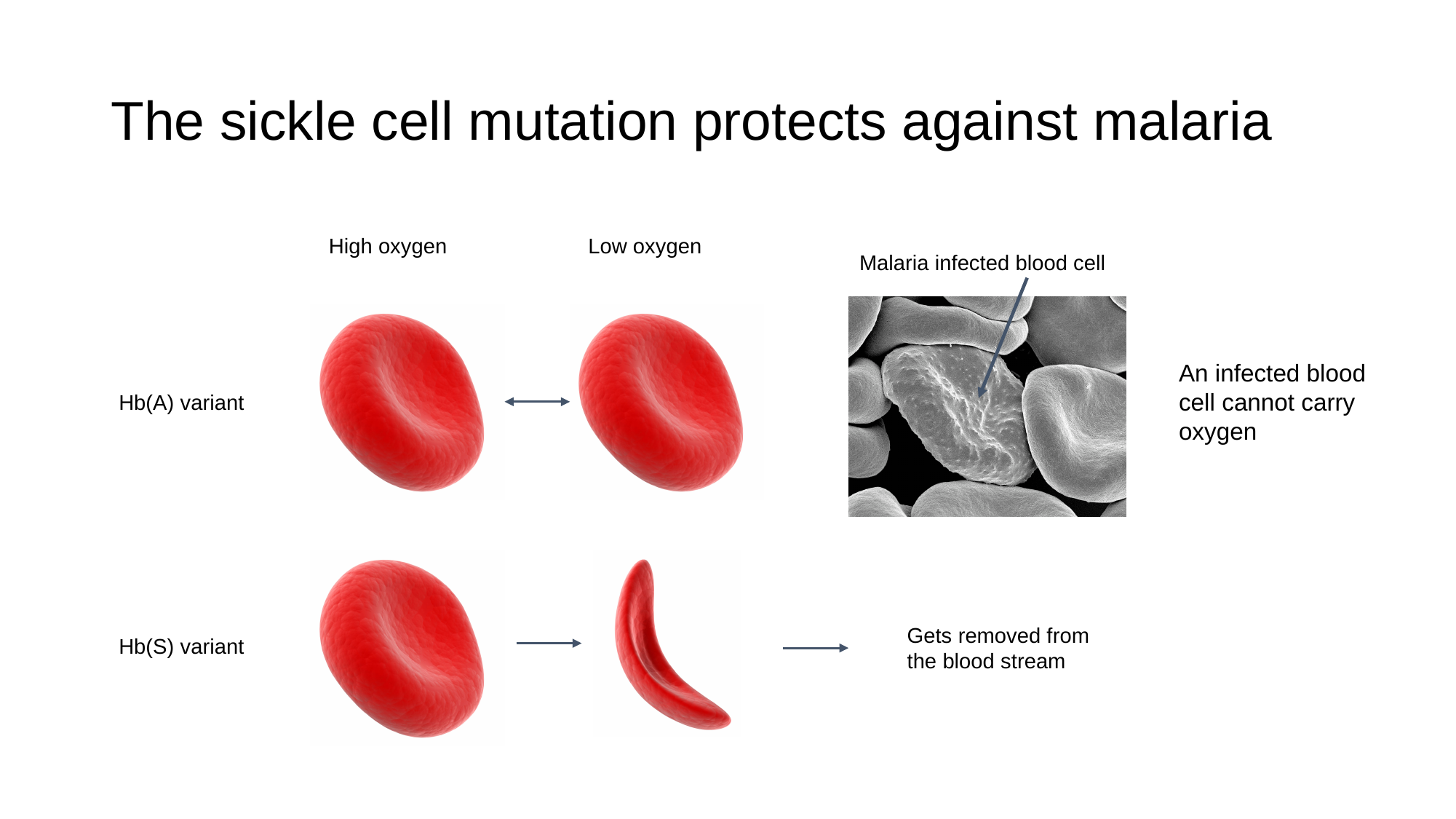

#### The sickle cell mutation protects against malaria
High oxygen
Low oxygen
Malaria infected blood cell
An infected blood cell cannot carry oxygen
Hb(A) variant
Gets removed from the blood stream
Hb(S) variant

### Slide 47
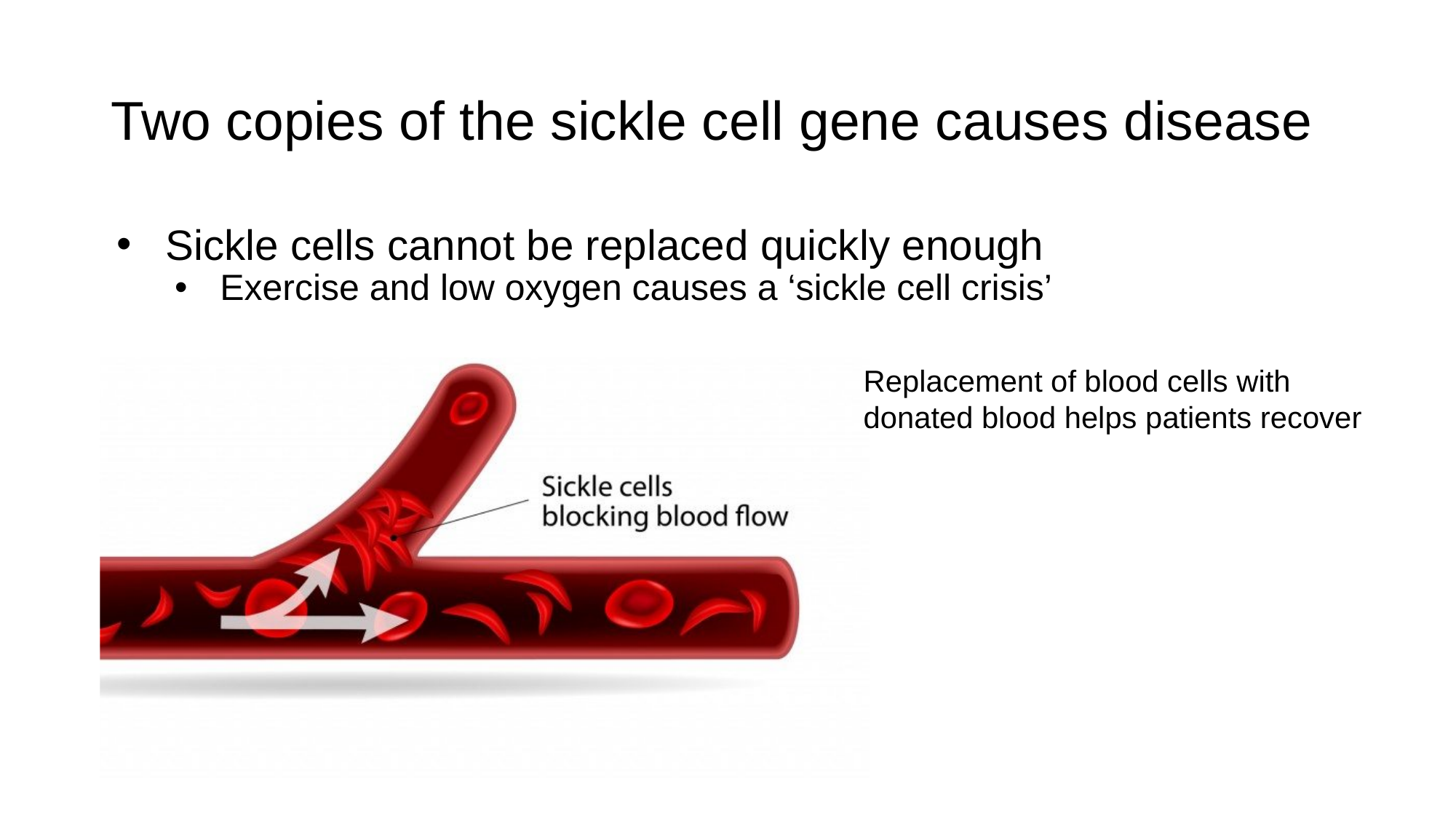

#### Two copies of the sickle cell gene causes disease
Sickle cells cannot be replaced quickly enough
Exercise and low oxygen causes a ‘sickle cell crisis’
Replacement of blood cells with donated blood helps patients recover

### Slide 48
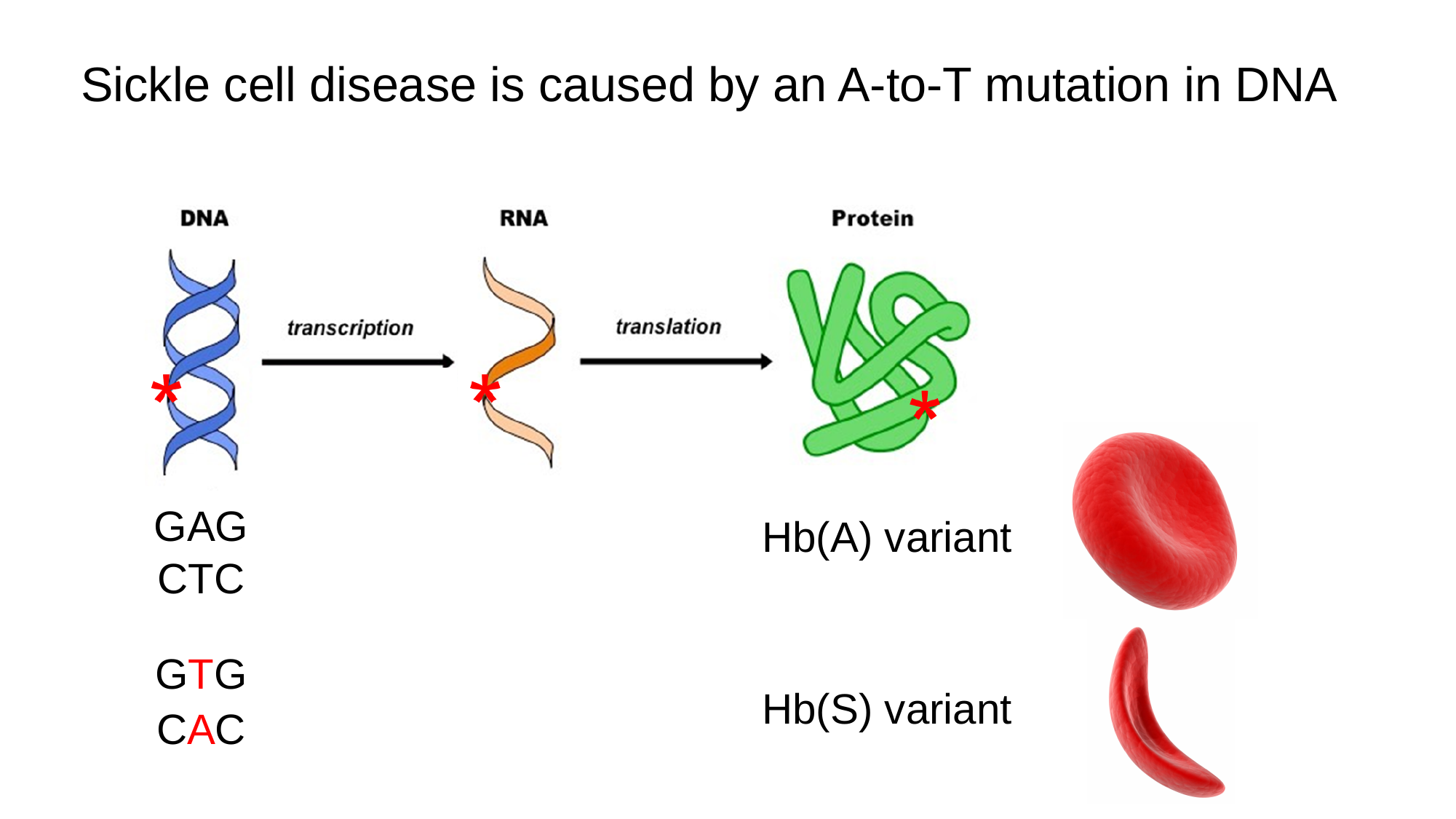

#### Sickle cell disease is caused by an A-to-T mutation in DNA
GAG
Hb(A) variant
CTC
GTG
Hb(S) variant
CAC

### Slide 49
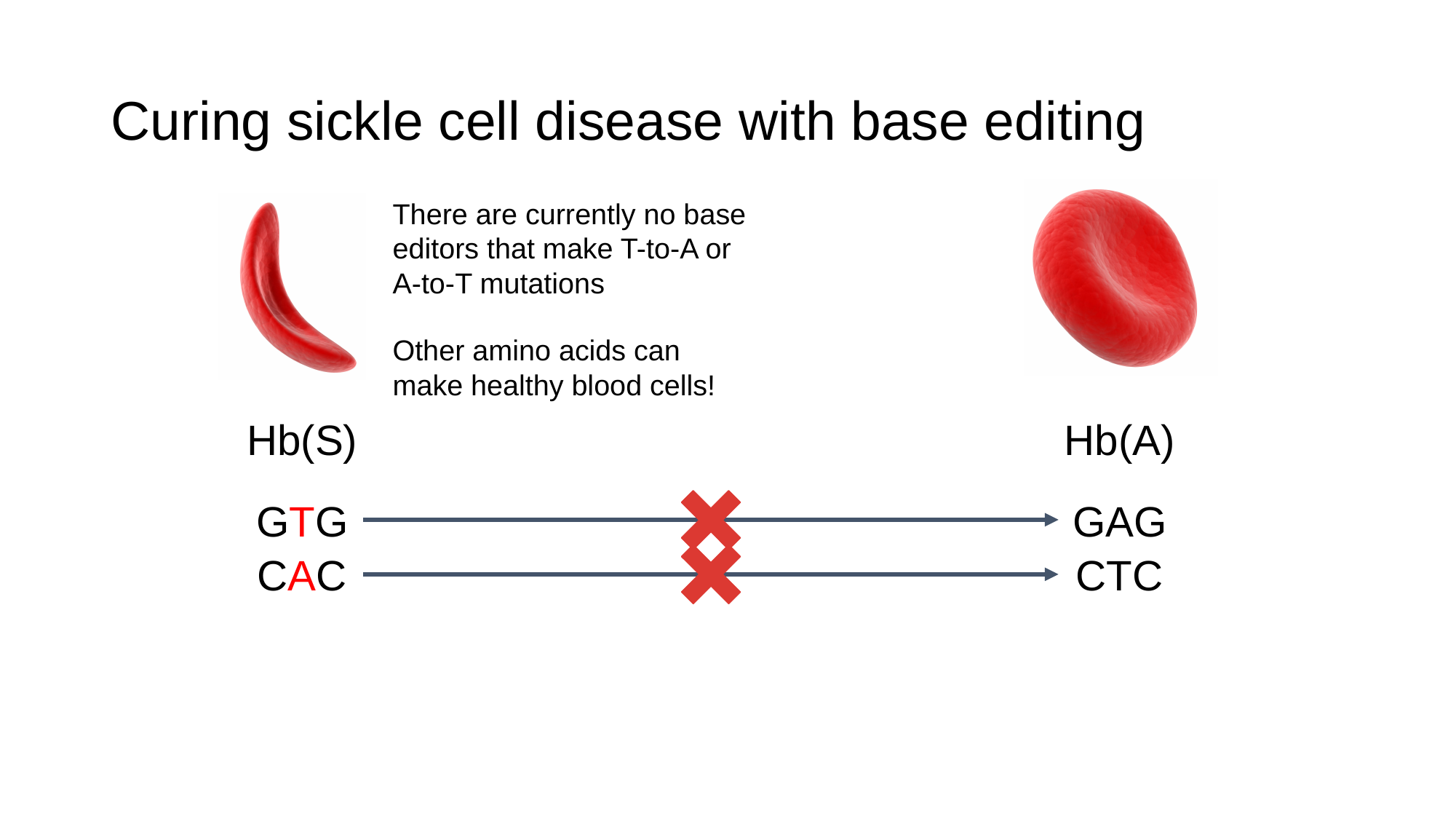

#### Curing sickle cell disease with base editing
There are currently no base editors that make T-to-A or A-to-T mutations
Other amino acids can make healthy blood cells!
Hb(S)
Hb(A)
GTG
GAG
CAC
CTC

### Slide 50
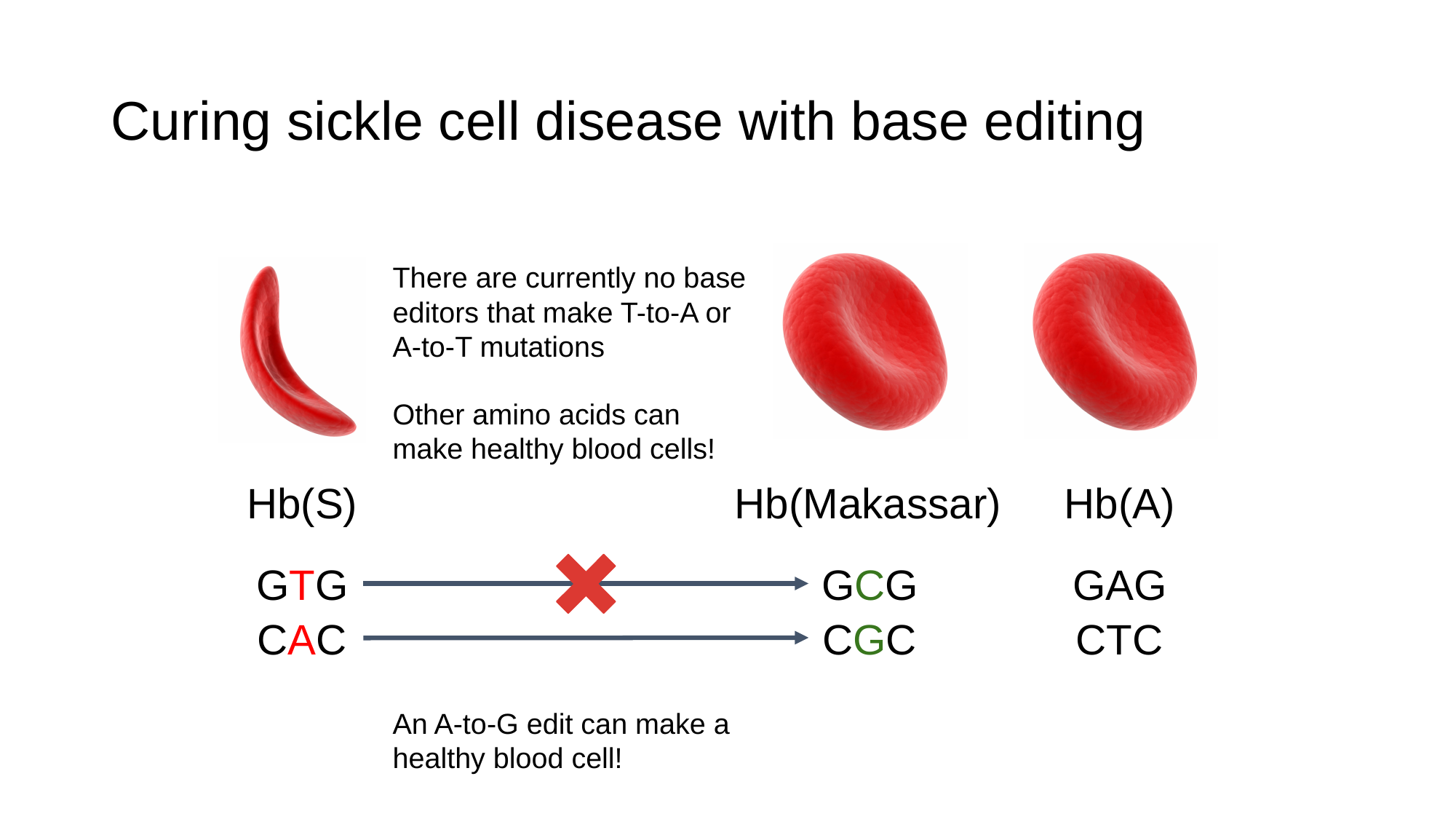

#### Curing sickle cell disease with base editing
There are currently no base editors that make T-to-A or A-to-T mutations
Other amino acids can make healthy blood cells!
Hb(S)
Hb(A)
Hb(Makassar)
GTG
GAG
GCG
CAC
CTC
CGC
An A-to-G edit can make a healthy blood cell!

### Slide 51
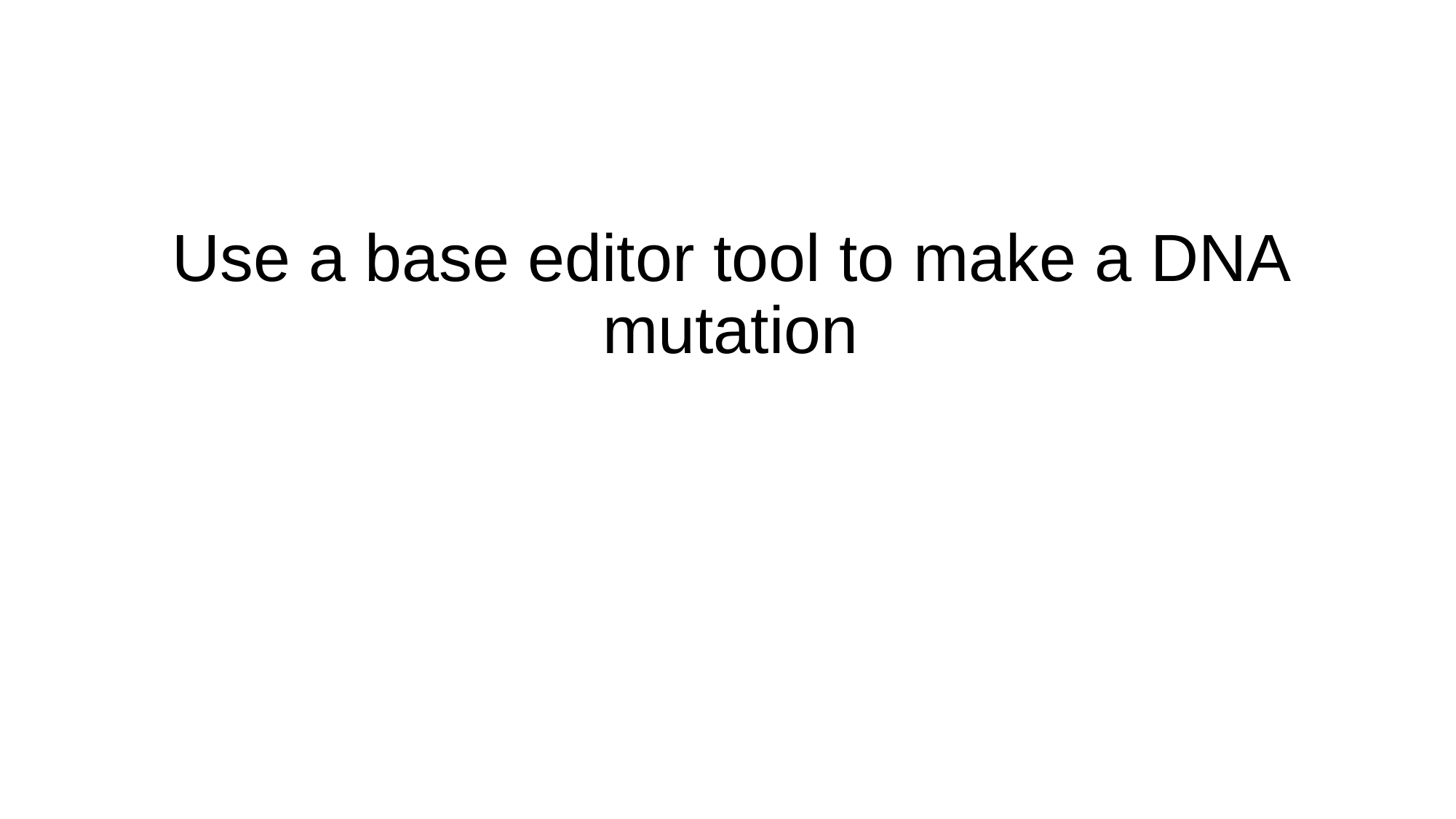

Use a base editor tool to make a DNA mutation

### Slide 52
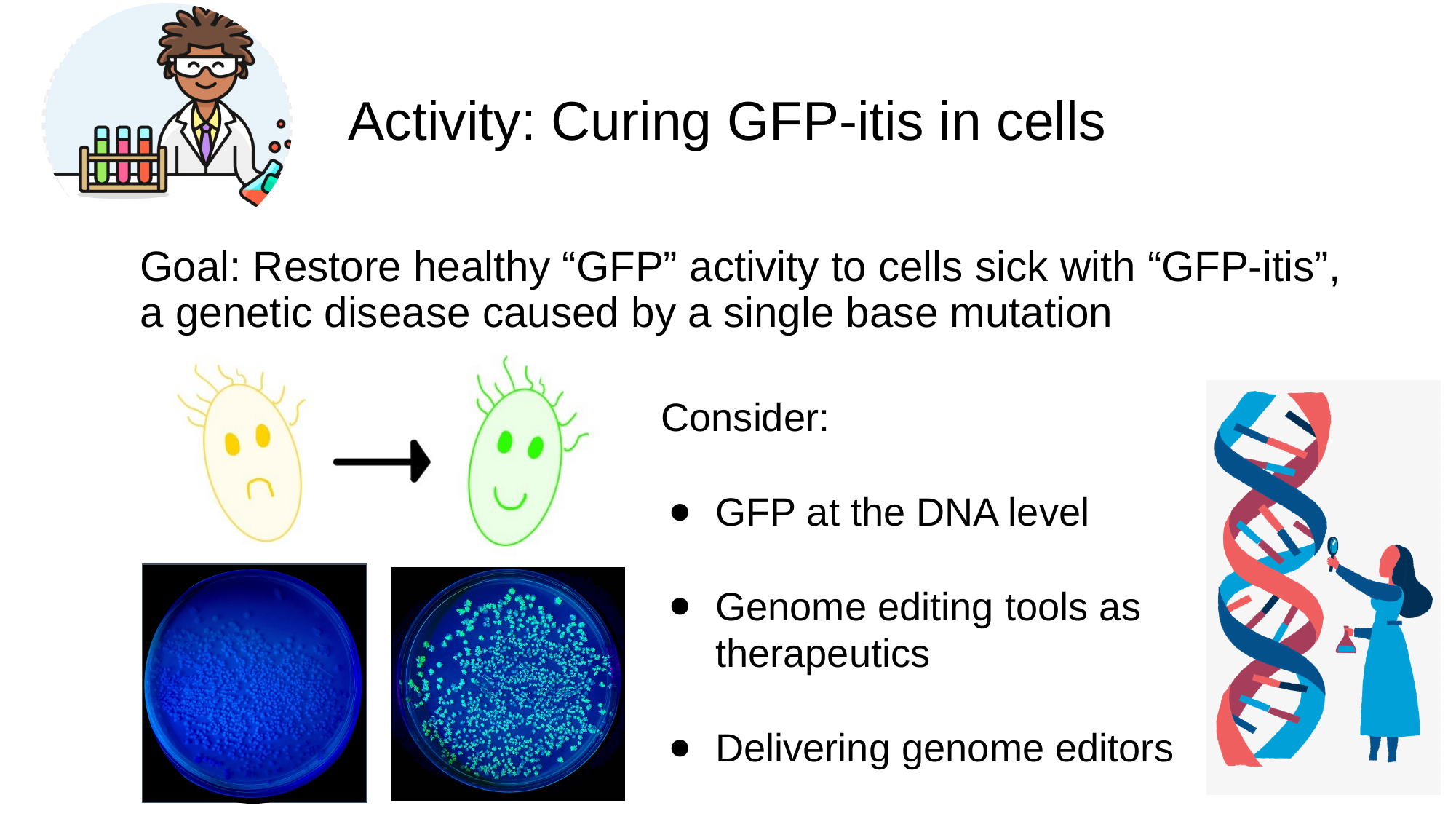

#### Activity: Curing GFP-itis in cells
Goal: Restore healthy “GFP” activity to cells sick with “GFP-itis”, a genetic disease caused by a single base mutation
Consider:
GFP at the DNA level
Genome editing tools as therapeutics
Delivering genome editors

### Slide 53
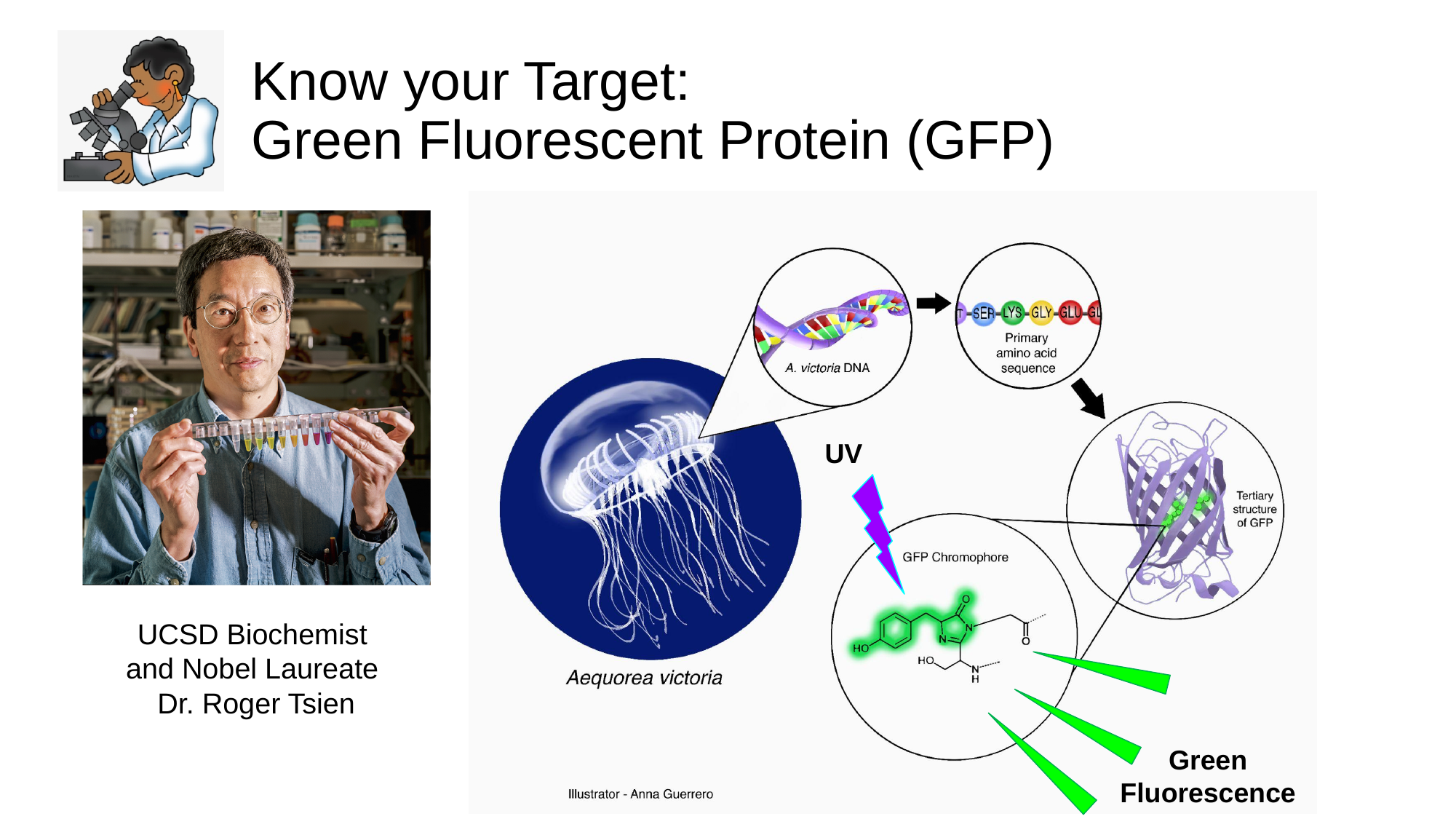

#### Know your Target: Green Fluorescent Protein (GFP)
UV
UCSD Biochemist
and Nobel Laureate
Dr. Roger Tsien
Green
Fluorescence

### Slide 54
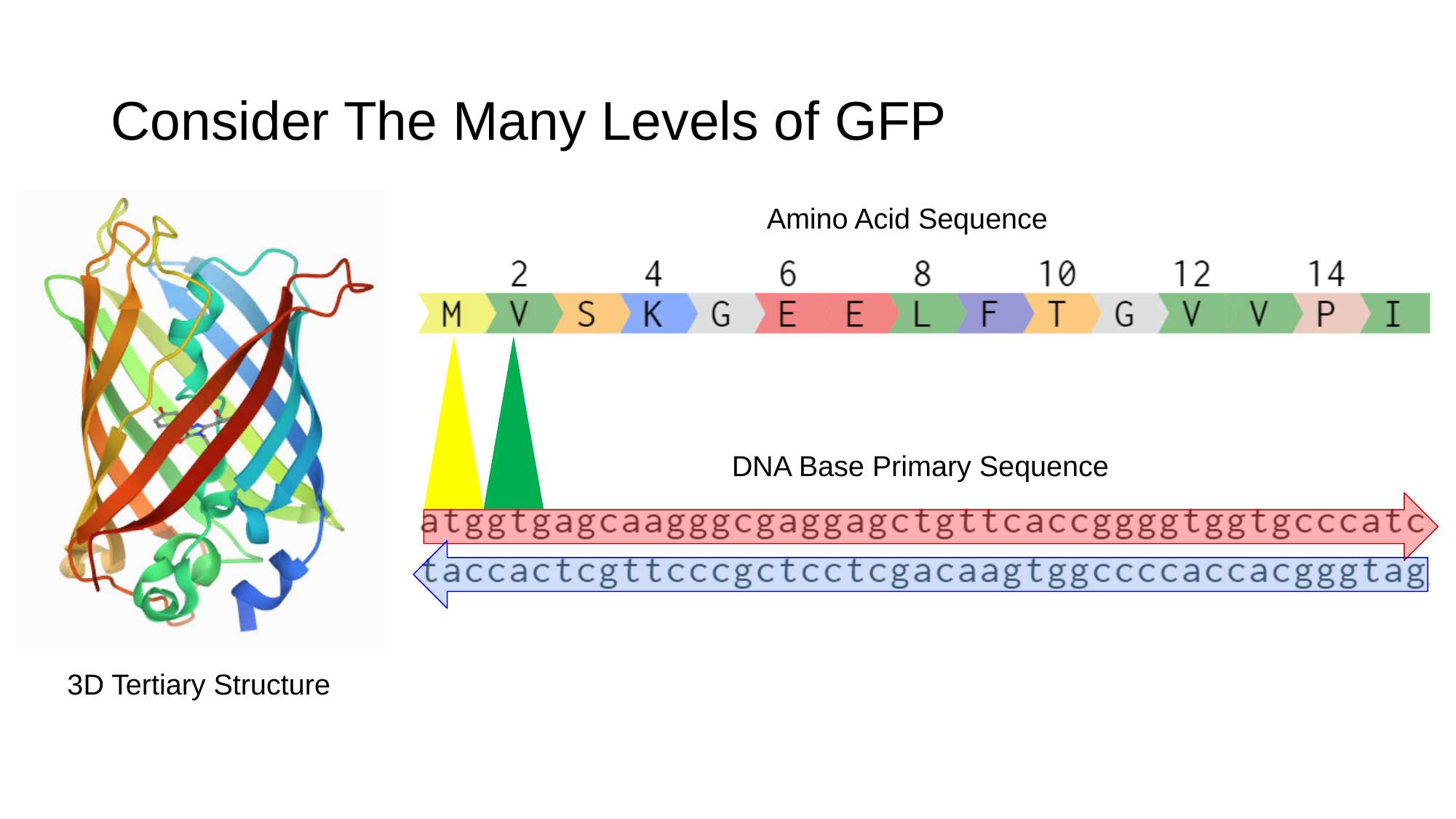

#### Consider The Many Levels of GFP
Amino Acid Sequence
DNA Base Primary Sequence
3D Tertiary Structure

### Slide 55
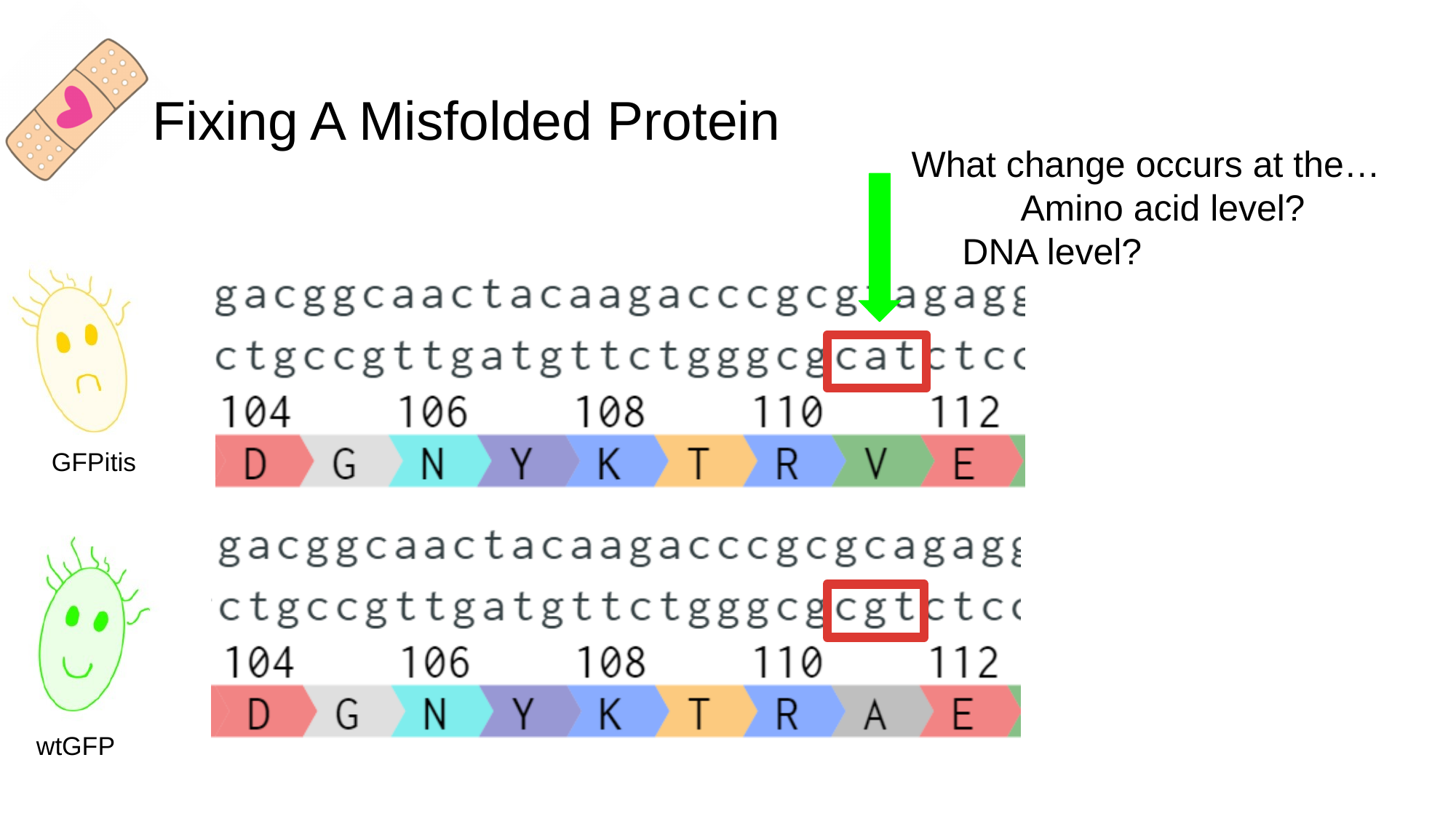

#### Fixing A Misfolded Protein
What change occurs at the…
	Amino acid level?
 DNA level?
GFPitis
wtGFP

### Slide 56
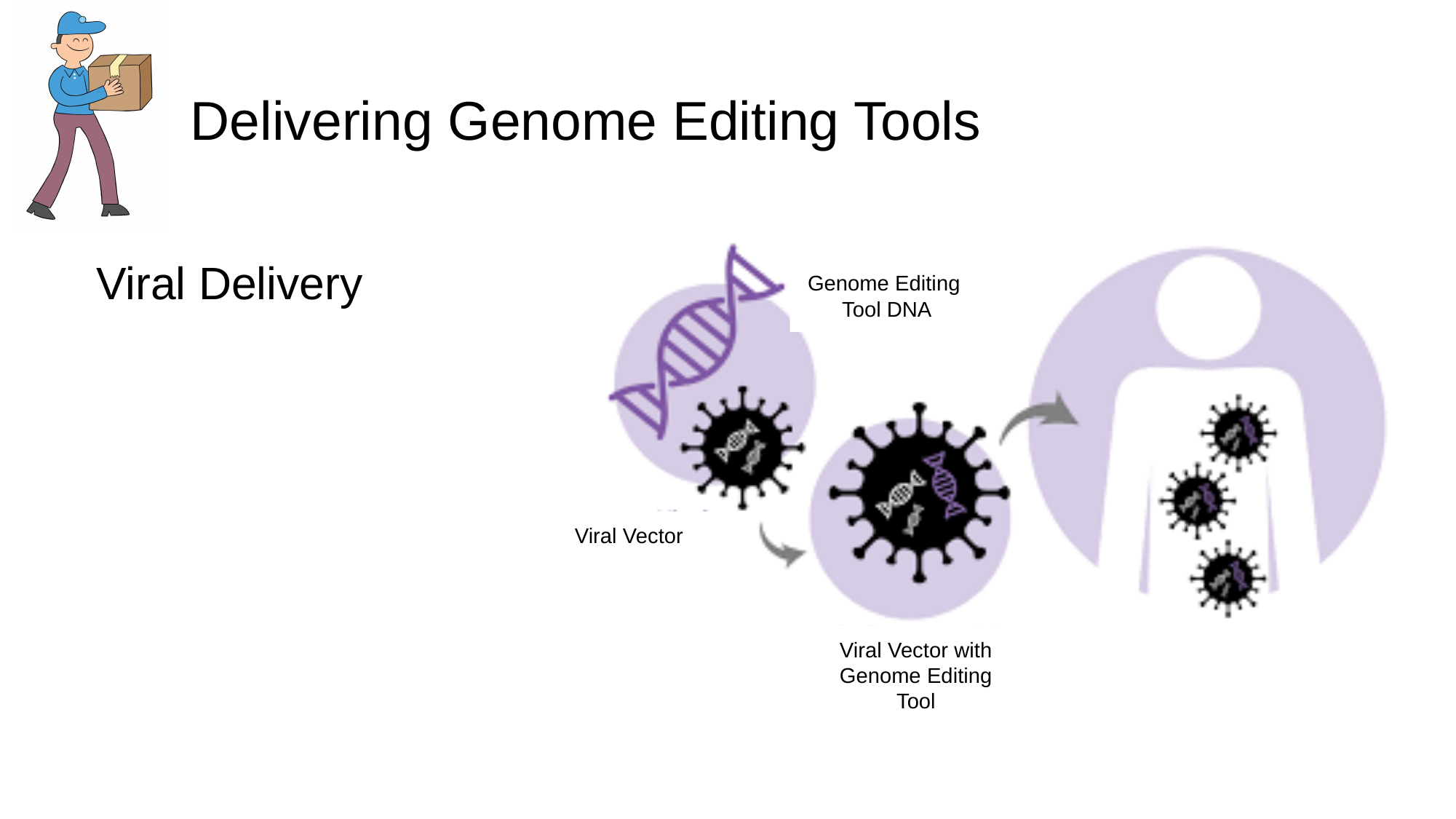

#### Delivering Genome Editing Tools
Viral Delivery
Genome Editing
Tool DNA
Viral Vector
Viral Vector with Genome Editing Tool

### Slide 57
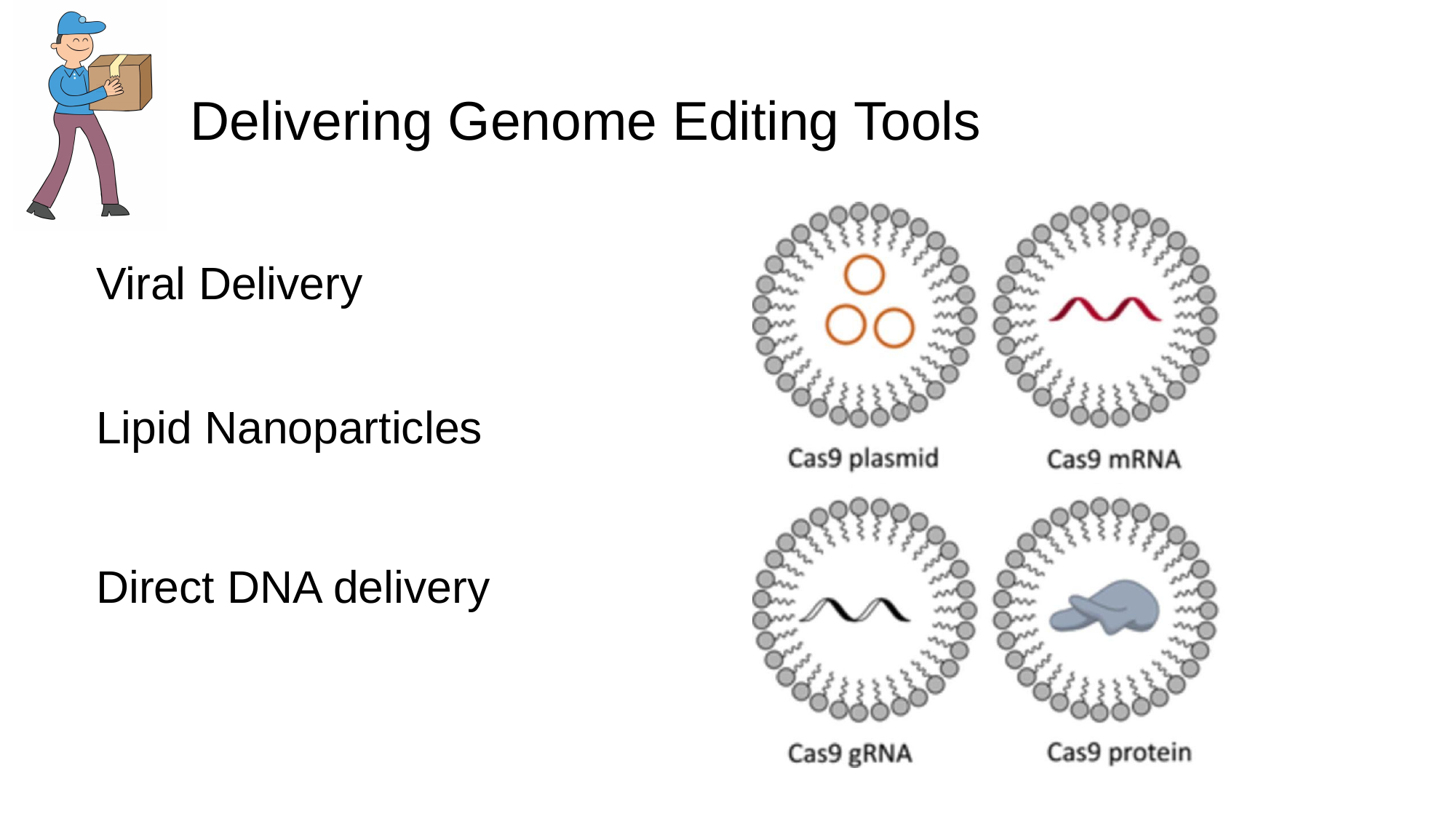

#### Delivering Genome Editing Tools
Viral Delivery
Lipid Nanoparticles
Direct DNA delivery

### Slide 58
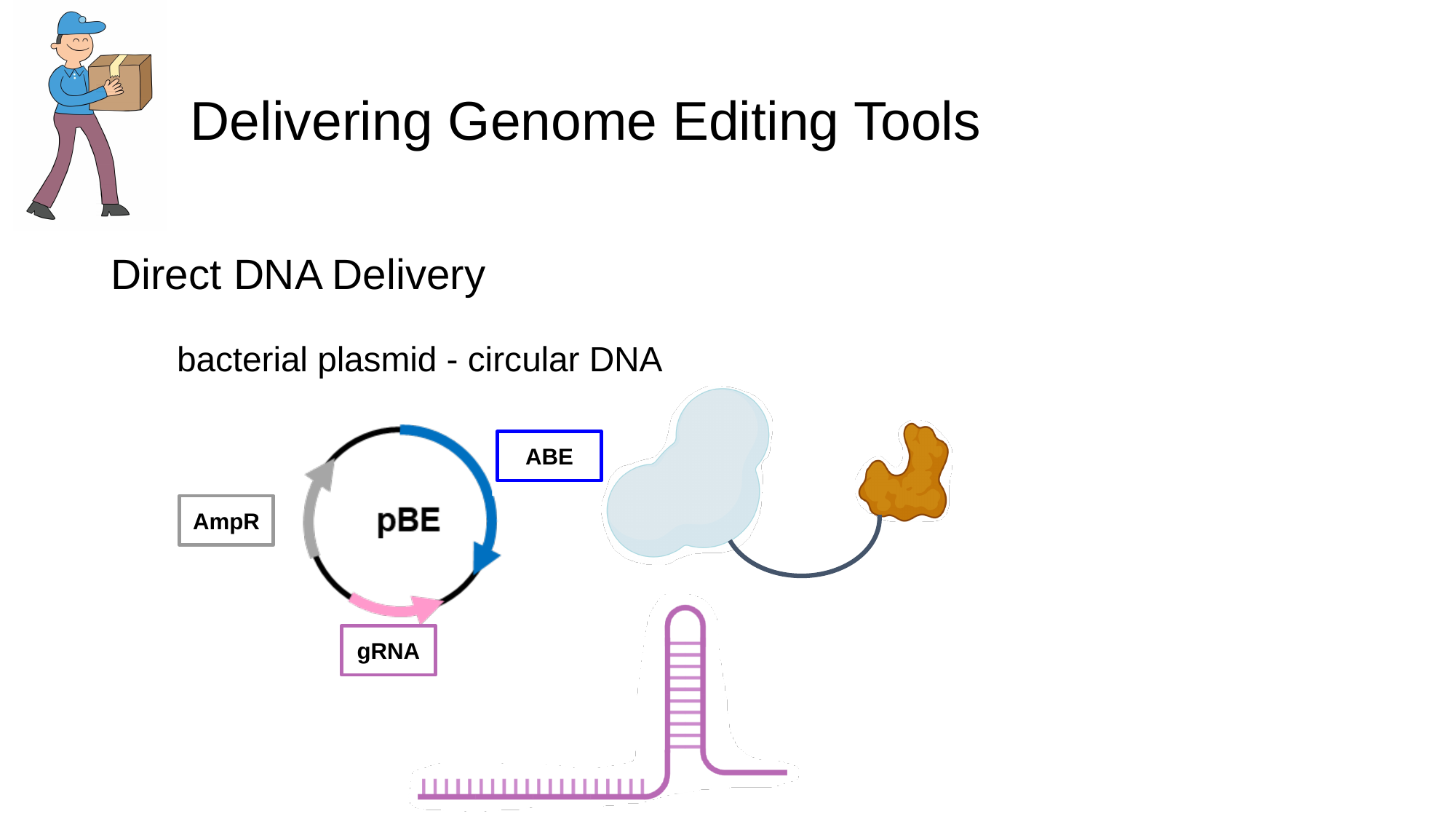

#### Delivering Genome Editing Tools
Direct DNA Delivery
bacterial plasmid - circular DNA
ABE
AmpR
gRNA

### Slide 59
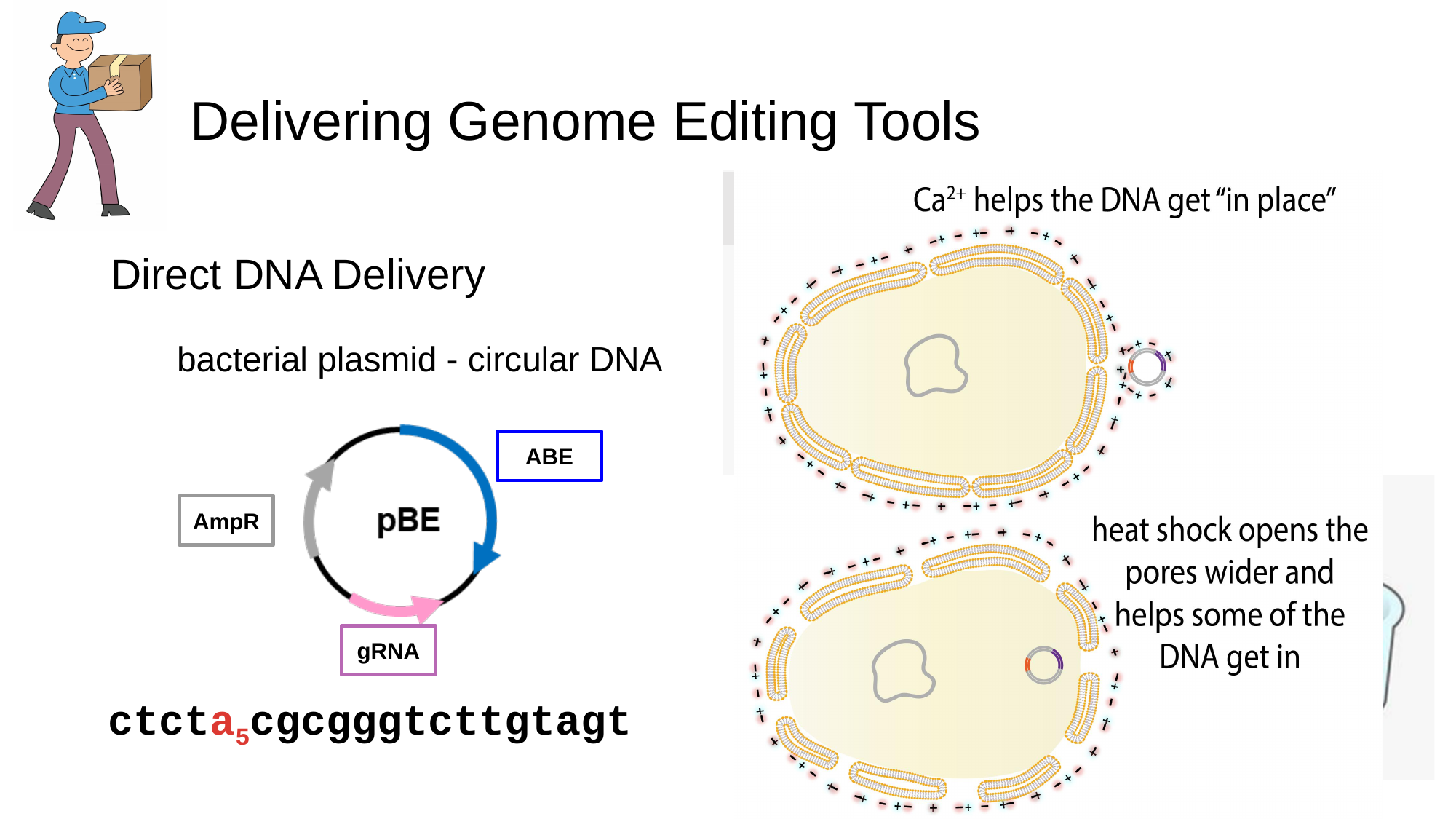

#### Delivering Genome Editing Tools
Competent bacterial cells
Plasmid/salt mixture
Direct DNA Delivery
bacterial plasmid - circular DNA
ABE
ABE
AmpR
AmpR
gRNA
gRNA
ctcta_5_cgcgggtcttgtagt

### Slide 60
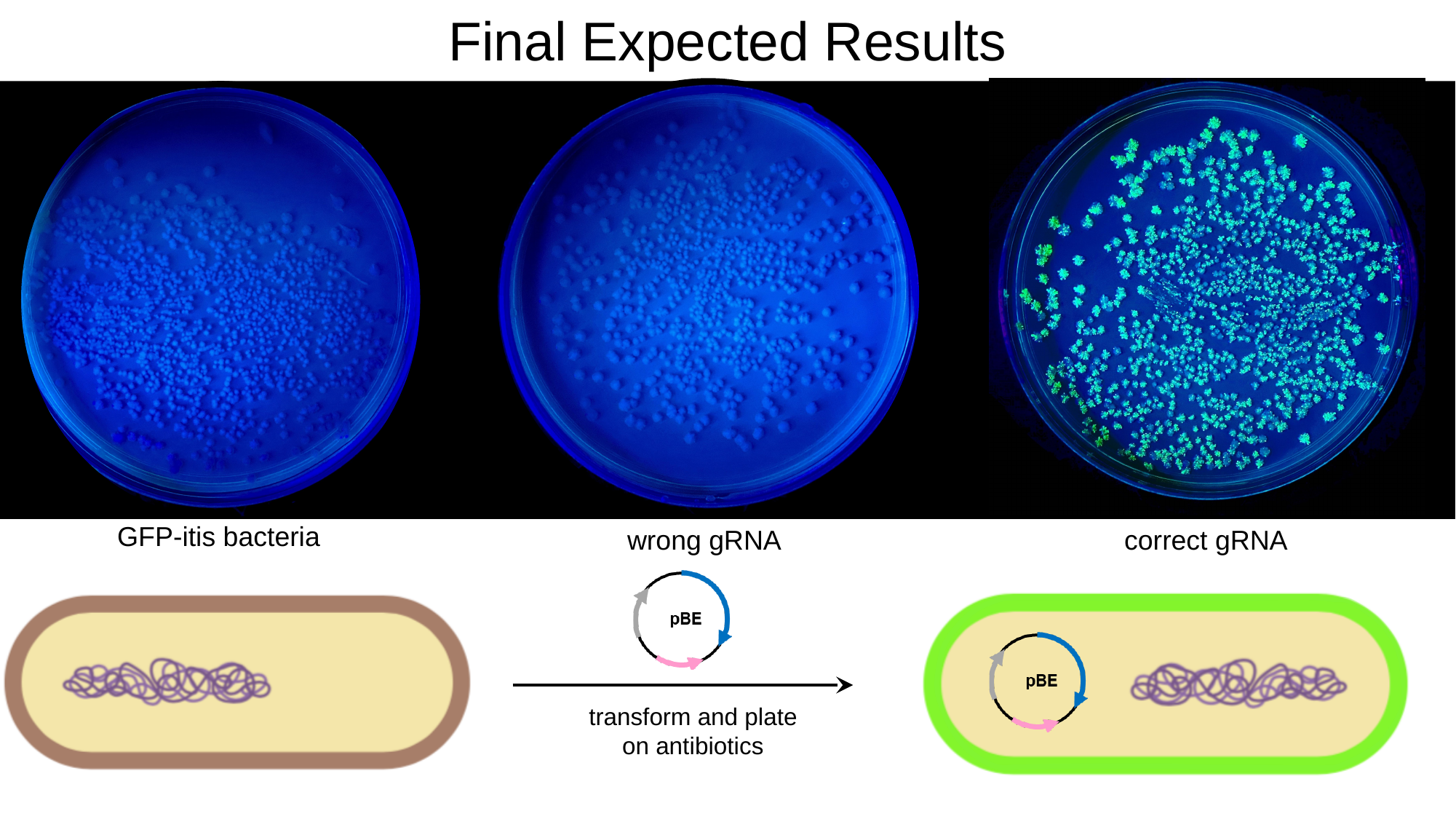

Final Expected Results
wrong gRNA
correct gRNA
GFP-itis bacteria
transform and plate on antibiotics

### Slide 61
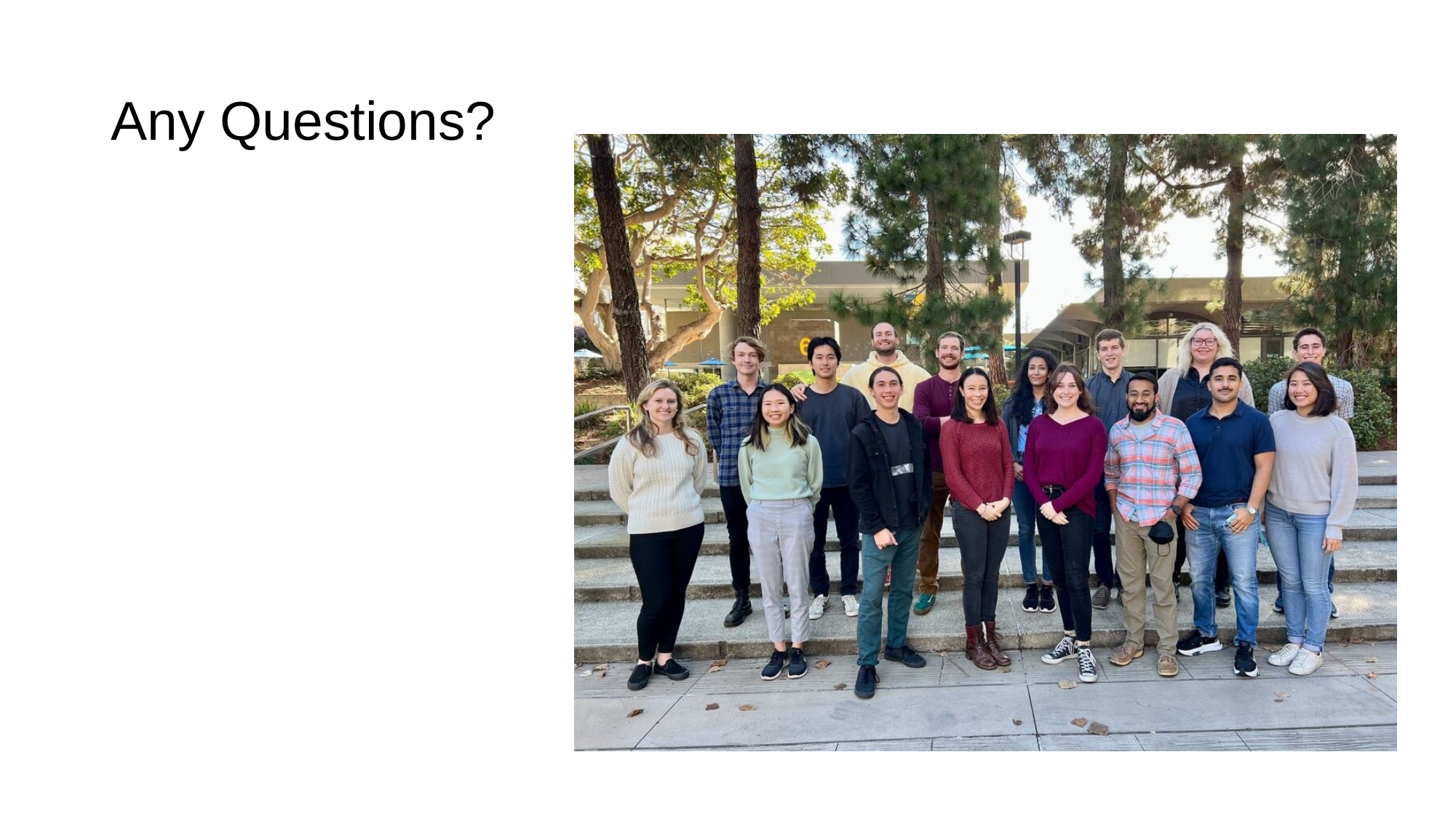

#### Any Questions?

### Slide 62
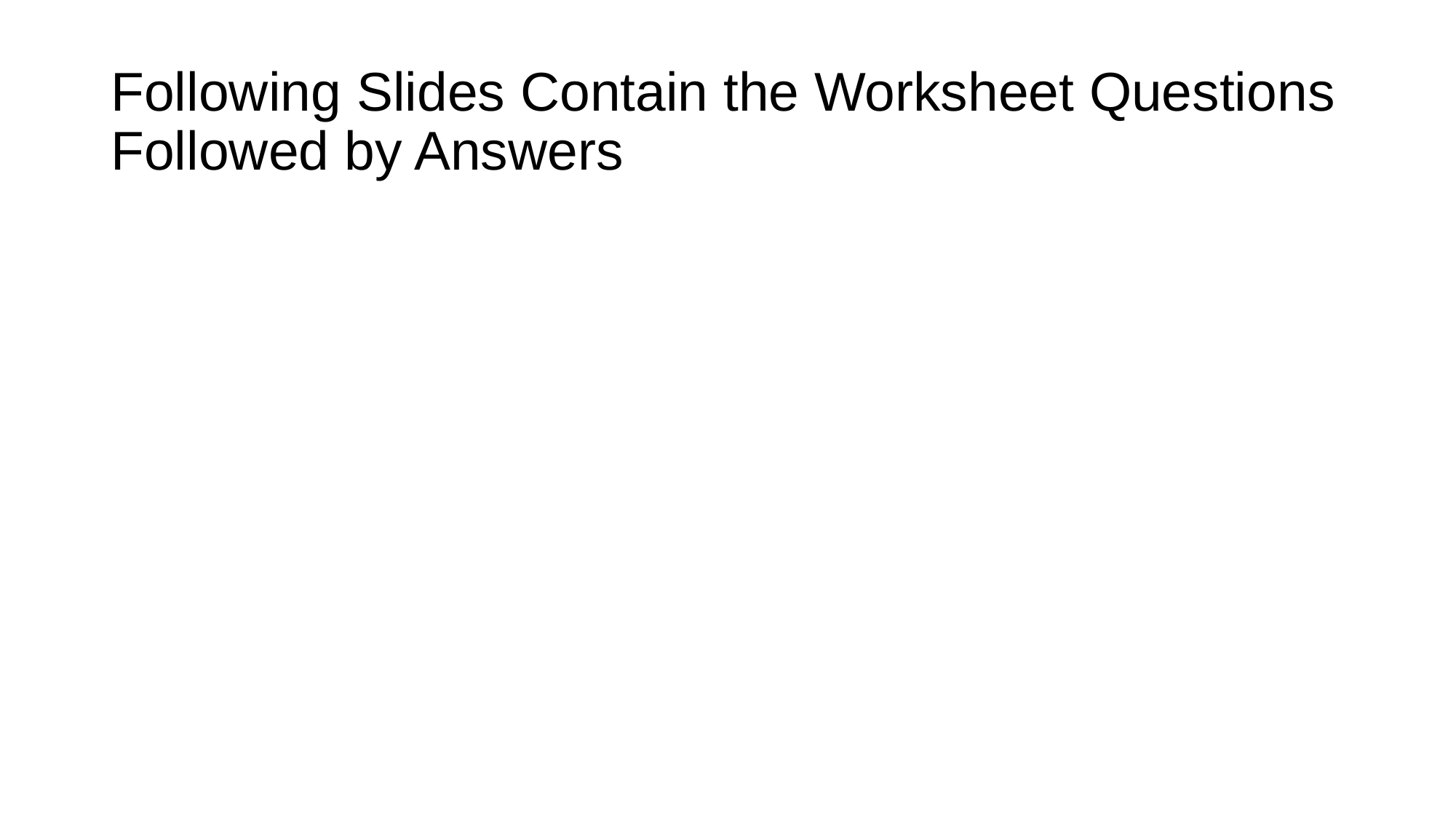

#### Following Slides Contain the Worksheet Questions Followed by Answers

### Slide 63
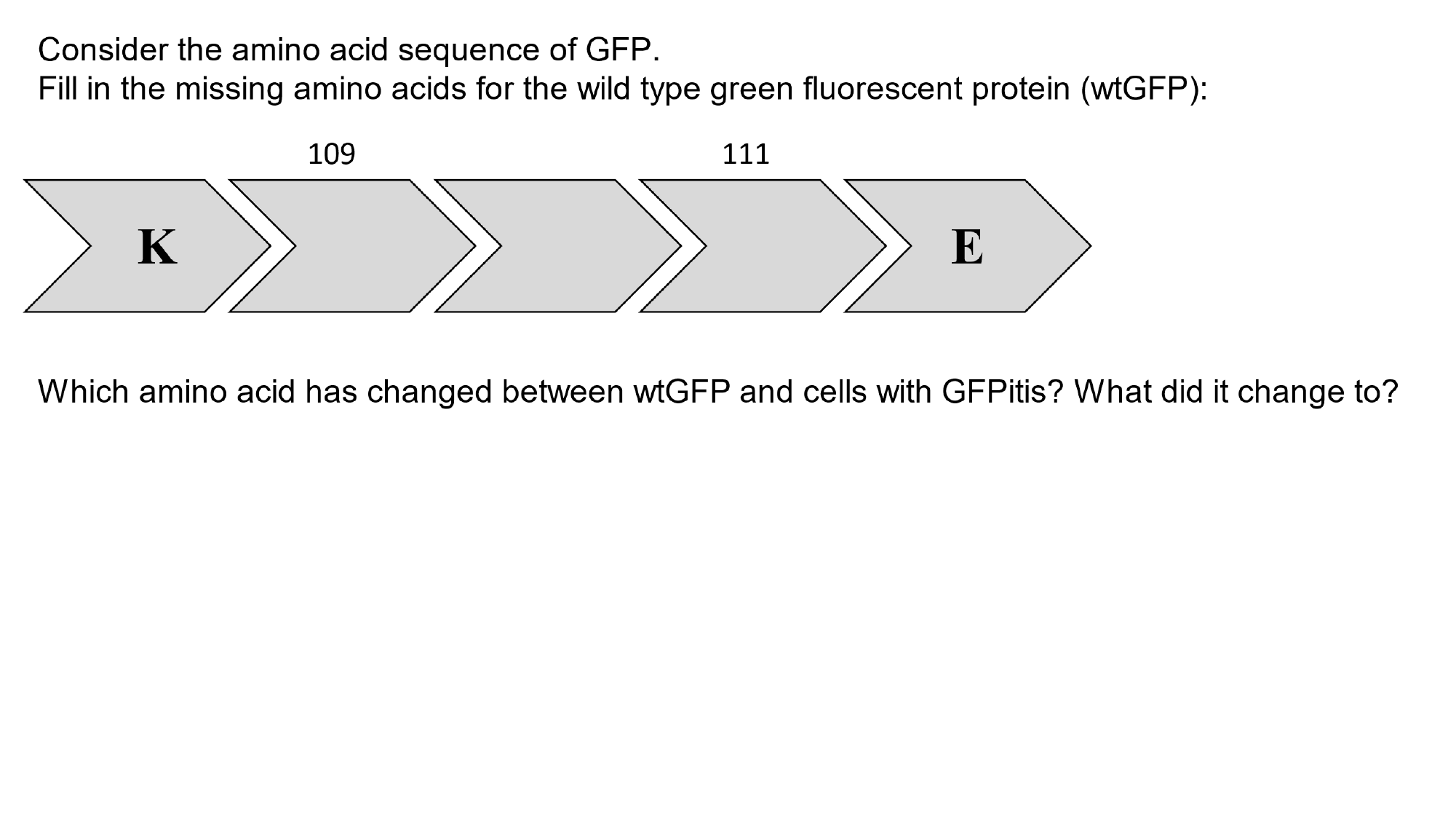

### Slide 64
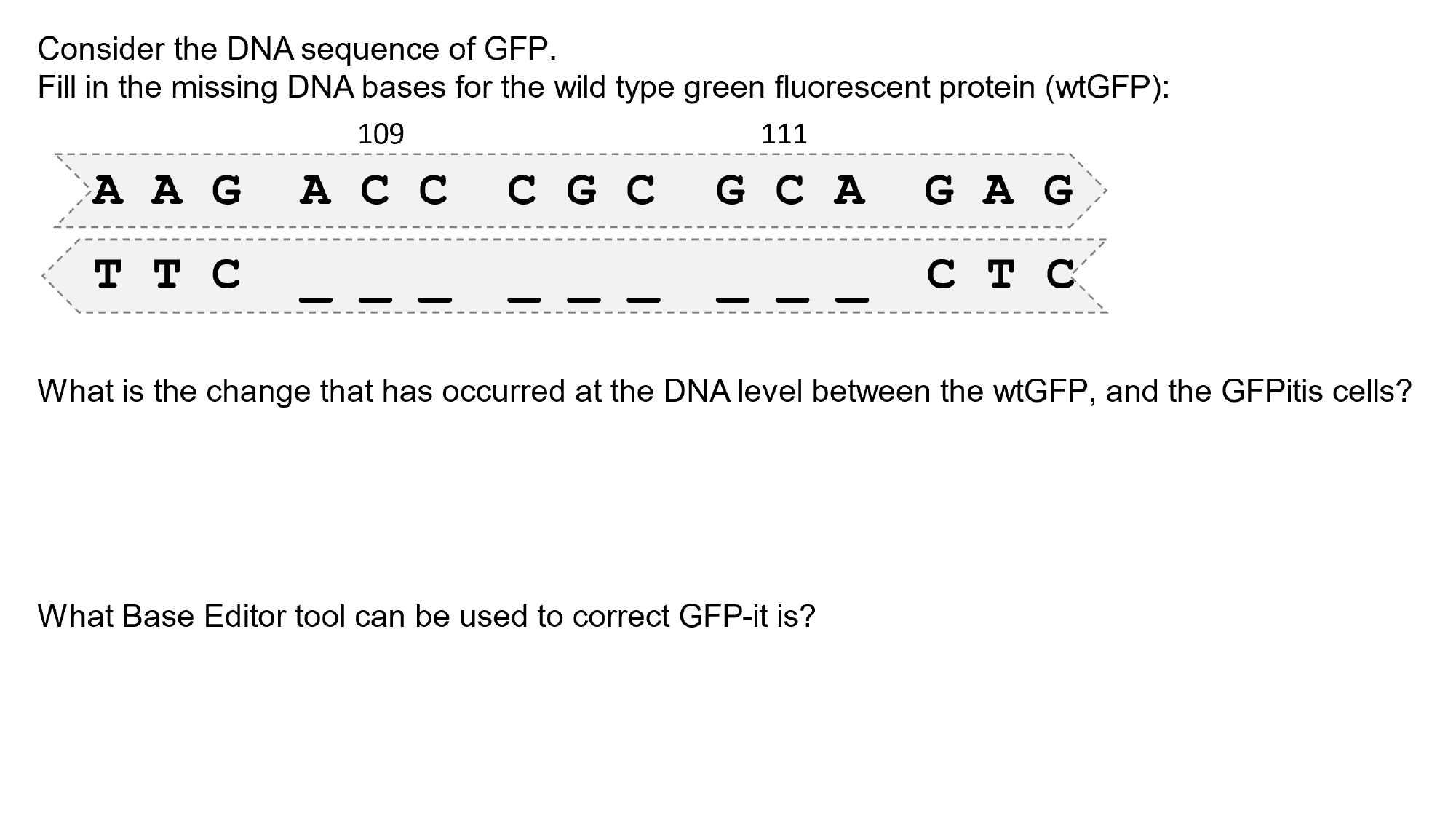

### Slide 65
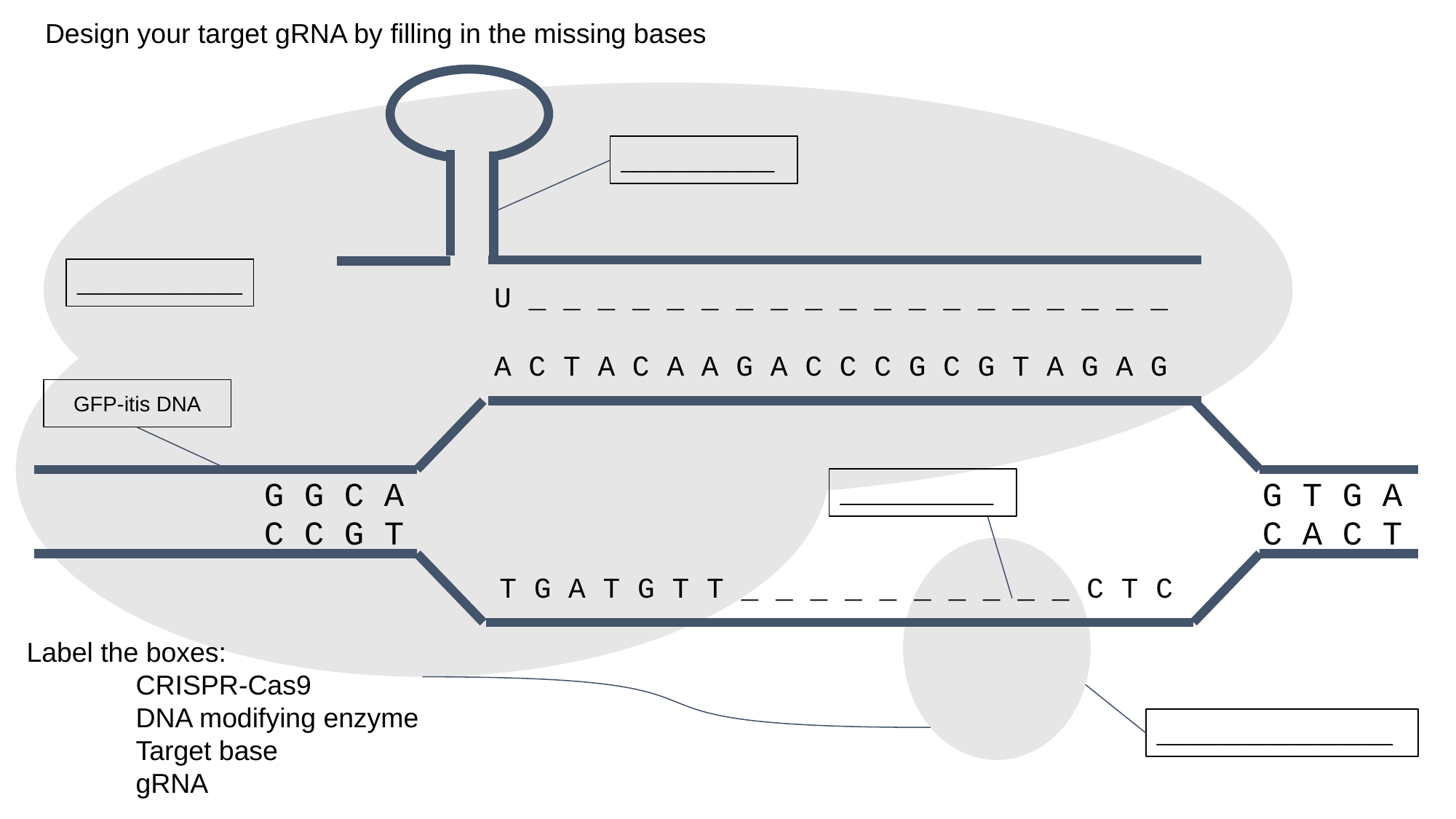

Design your target gRNA by filling in the missing bases
_____________
______________
U _ _ _ _ _ _ _ _ _ _ _ _ _ _ _ _ _ _ _
A C T A C A A G A C C C G C G T A G A G
GFP-itis DNA
 G G C A
G T G A
_____________
 C C G T
C A C T
T G A T G T T _ _ _ _ _ _ _ _ _ _ C T C
Label the boxes:
	CRISPR-Cas9
	DNA modifying enzyme
	Target base
	gRNA
____________________

### Slide 66
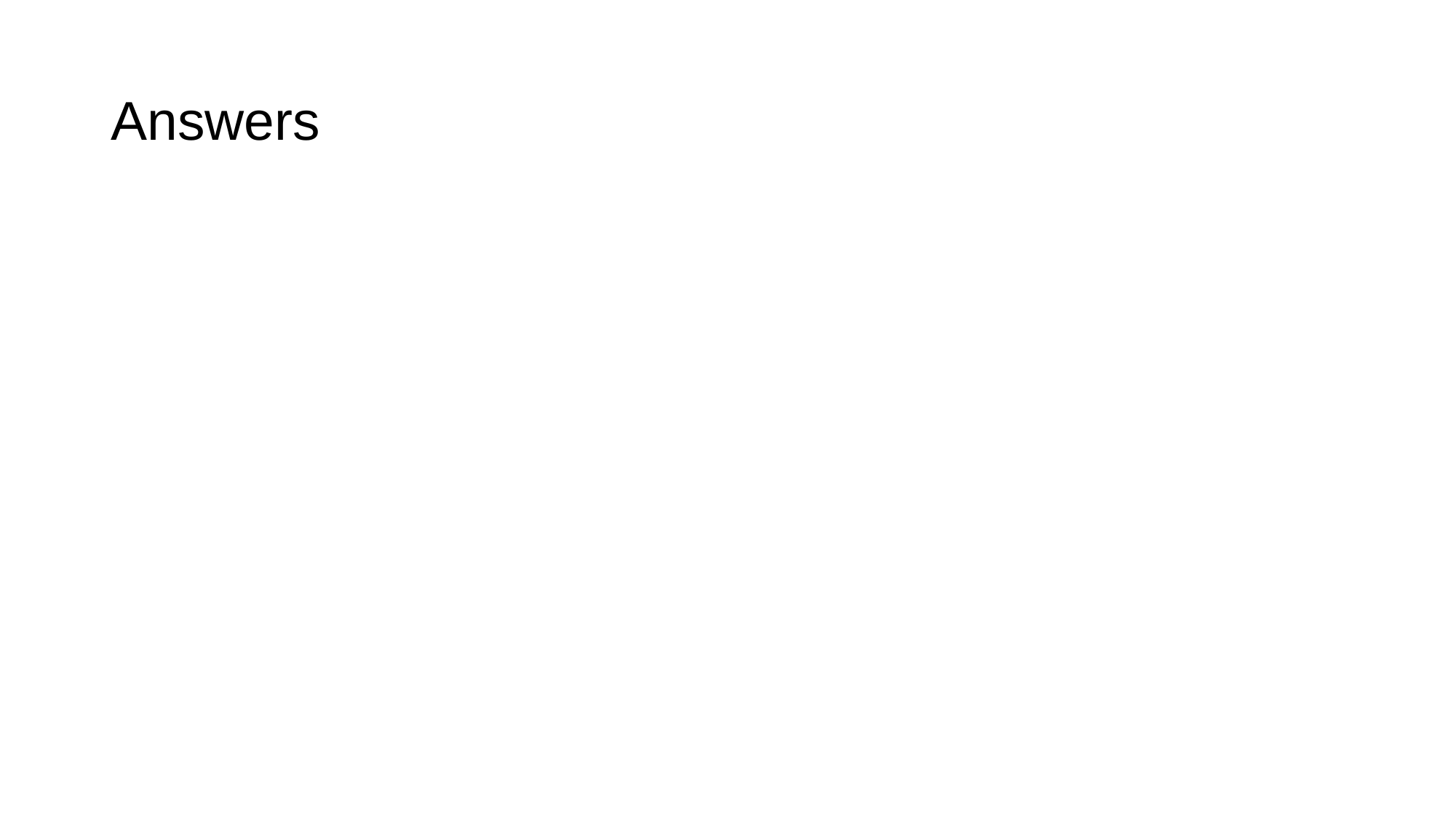

#### Answers

### Slide 67
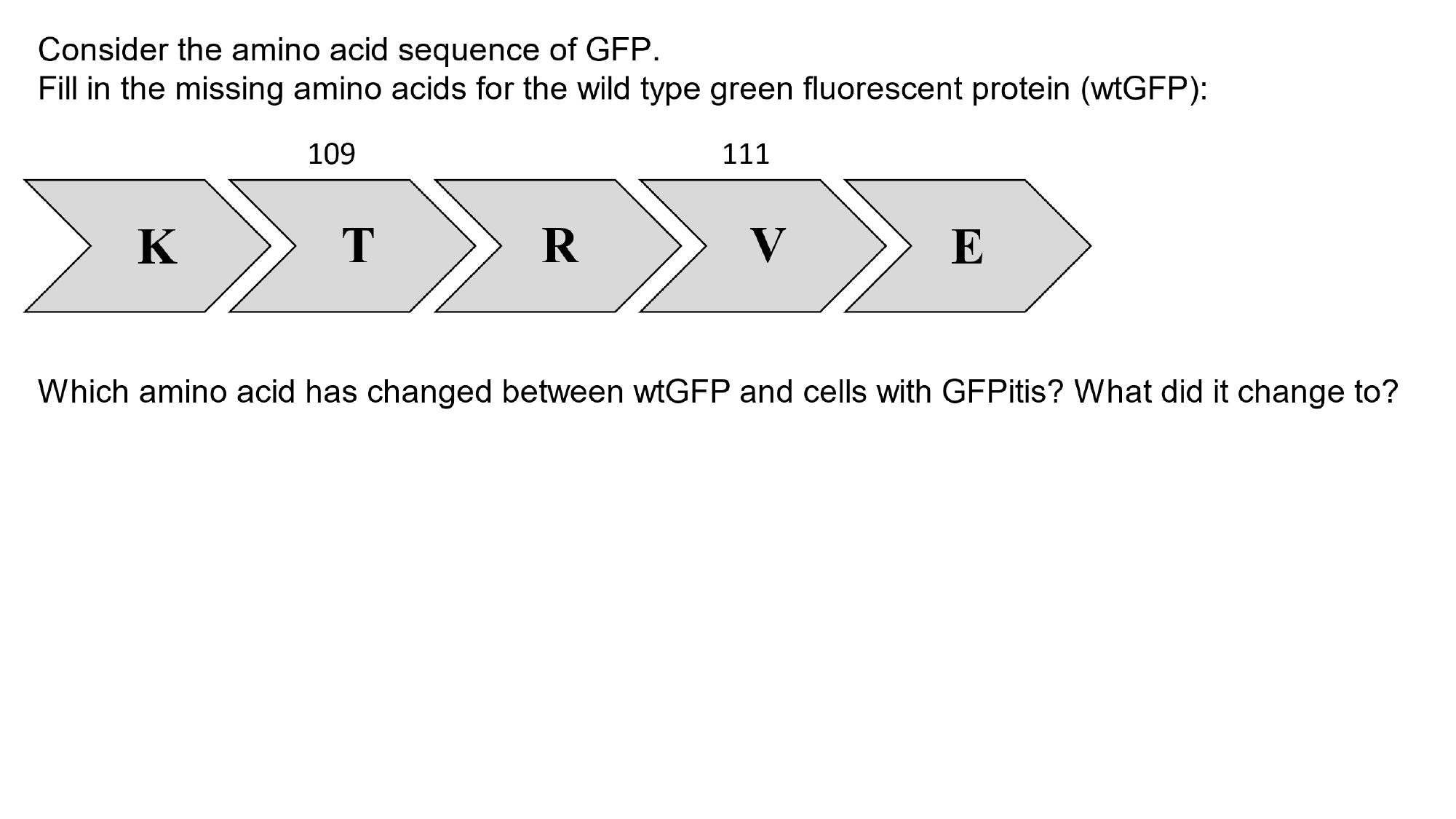

### Slide 68
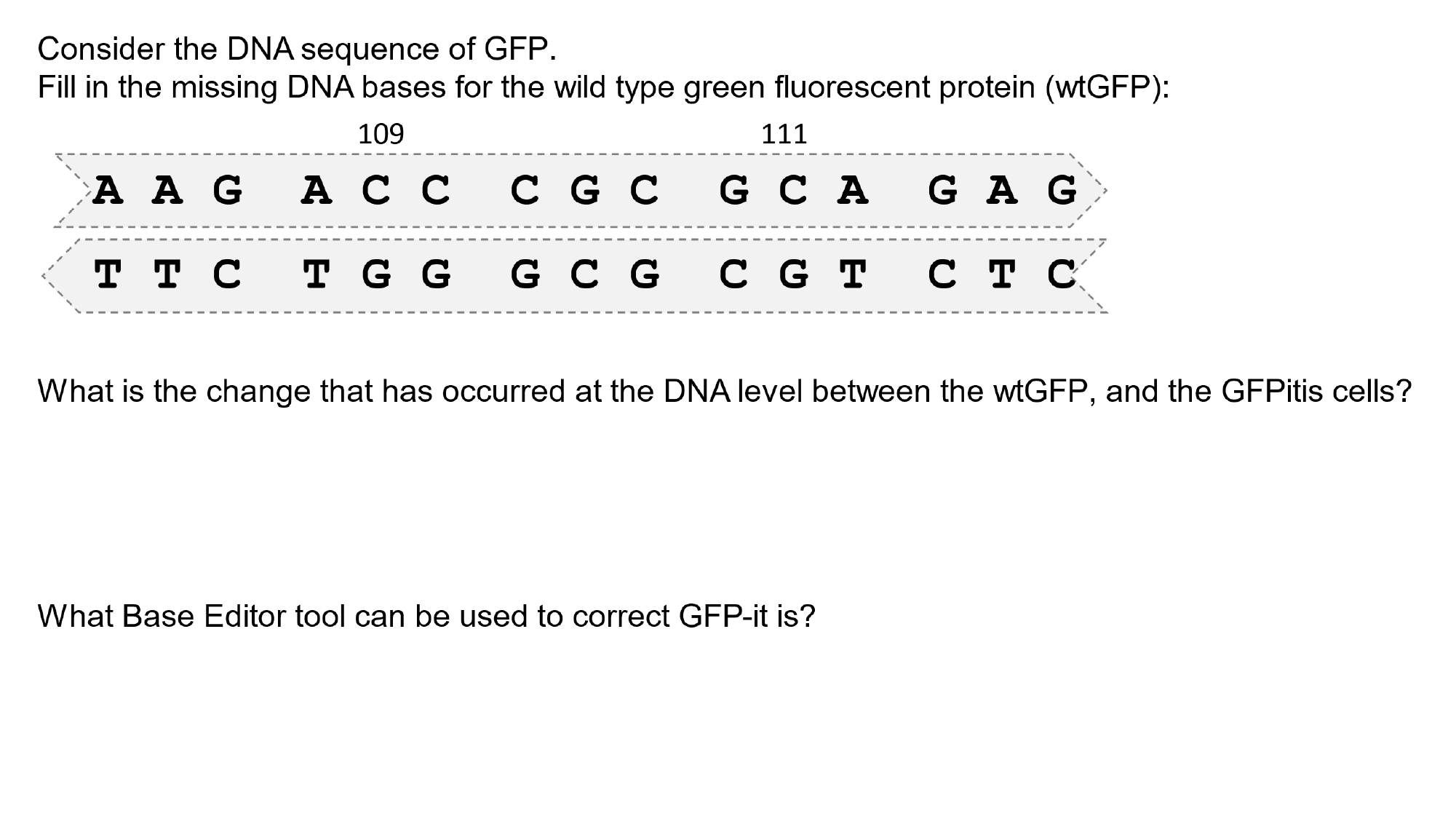

### Slide 69
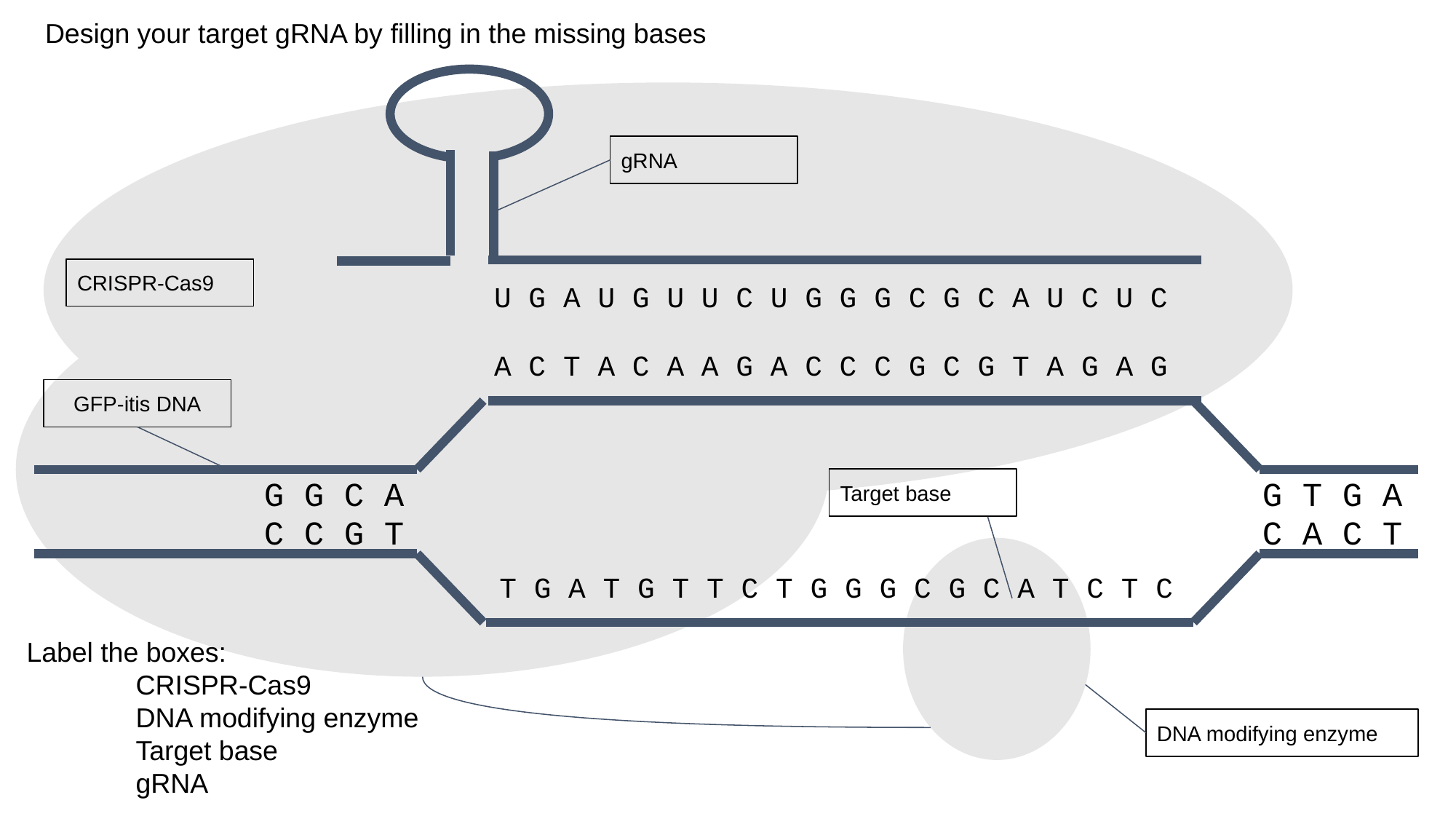

Design your target gRNA by filling in the missing bases
gRNA
CRISPR-Cas9
U G A U G U U C U G G G C G C A U C U C
A C T A C A A G A C C C G C G T A G A G
GFP-itis DNA
 G G C A
G T G A
Target base
 C C G T
C A C T
T G A T G T T C T G G G C G C A T C T C
Label the boxes:
	CRISPR-Cas9
	DNA modifying enzyme
	Target base
	gRNA
DNA modifying enzyme
