## Supplemental Material - Worksheet for "Curing “GFP-itis” in Bacteria with Base Editors: Development of a Genome Editing Science Program Implemented with High School Biology Students"

Name: ___________________________

### BASE EDITING WORKSHOP

**Use this packet to take notes, write questions, and make observations throughout the workshop and activity:**

Consider the amino acid sequence of GFP.
Fill in the missing amino acids for the wild type green fluorescent protein (wtGFP):

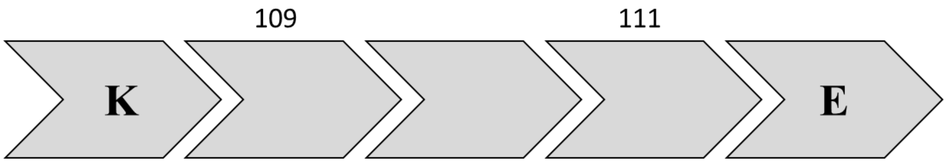


Which amino acid has changed between wtGFP and cells with GFPitis? What did it change to?

Consider the DNA sequence of GFP.
Fill in the missing DNA bases for the wild type green fluorescent protein (wtGFP):

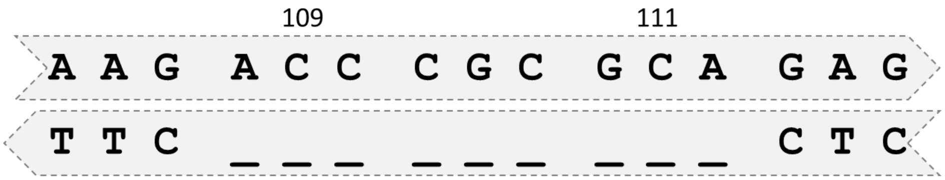


What is the change that has occurred at the DNA level between the wtGFP, and the GFPitis cells?

What Base Editor tool can be used to correct GFP-it is?

Name: ____________________________

Design your gRNA by filling in the missing bases, and labeling the different parts of the Base Editor tool:

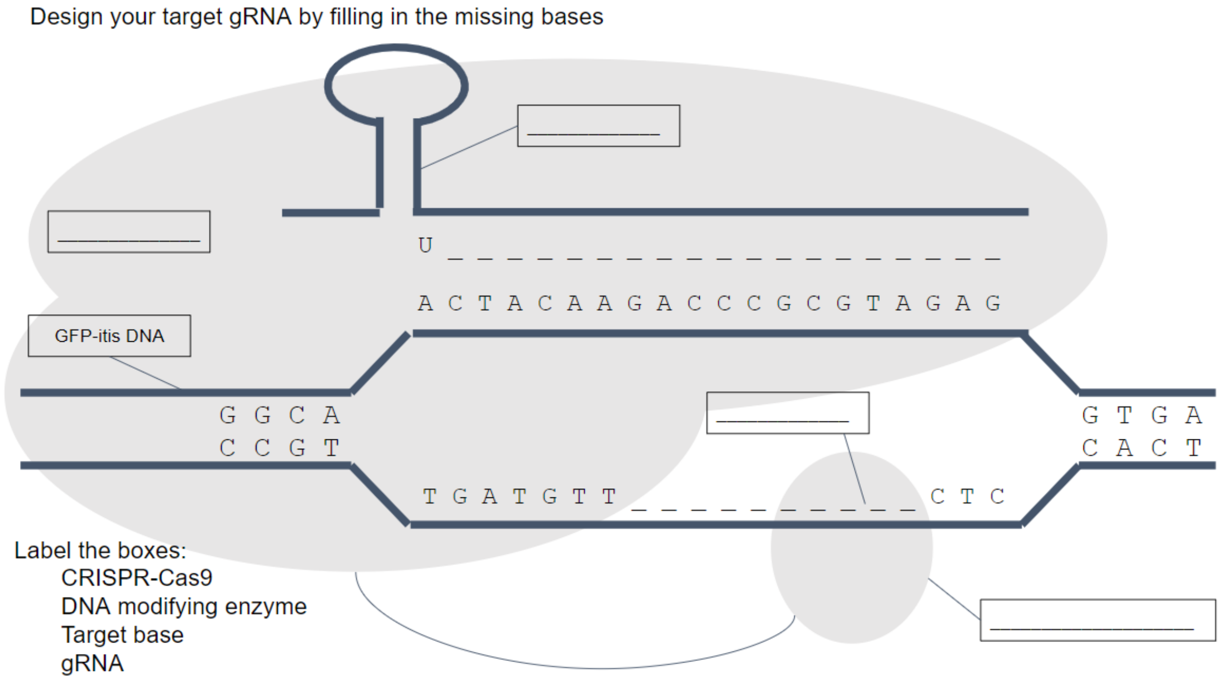


**Day 1 Questions to Consider:**

1. Will the bacteria fluoresce green when using pBE-NT (non-targeting gRNA)? Explain your reasoning.

2. When using the pBE-T (targeting gRNA), what base will the adenosine deaminase base editor change? What will it change to? What effect will this have?

3. There is currently a lot of debate surrounding the ethics of genome editing. Can you think of an ethical question(s) scientists need to ask/answer before using this technology? (Keep them in mind for the Q&A Panel on Day 3).

Name: ___________________________

### BASE EDITING ACTIVITY

**Materials:**

*Biologics and Chemicals*:

- 2 plasmids
  - Targeting gRNA base editor plasmid (pBE-t)
  - Non-targeting gRNA base editor plasmid (pBE-nt)
- 2 aliquots chemically competent GFP-itis bacterial cells
- 5x KCM salt solution
- Sterile DI water
- Sterile bacterial recovery media

Standard lab supplies

- 1.7mL snap top tube
- Lab markers
- Micropipettes and tips
  - 200-1000μL
  - 20-100μL
  - 1-10μL
- Bacterial spreader loops
- Petri dishes with Agar containing maintenance antibiotic (50ng/uL ampicillin)
- Ice
- Water bath (42C)
- Bacterial incubator (37C)
- UV light source (~470nm)
- UV light filtering glasses

**Safety and Waste Disposal Reminders**

While working with bacteria, be sure to dispose of any tips, spreader loops and empty tubes in appropriate Biohazard waste containers. Always wear UV filter glasses when working with blue/UV light.

**Micropipette Reminders (courtesy of Edvotek pipettes)**

1. **SET** the micropipette to the appropriate volume by adjusting the dial.
2. **PLACE** a clean tip on the micropipette.
3. **PRESS** the plunger down to the *first* stop. **HOLD** the plunger down while placing the tip beneath the surface of the liquid.
4. Slowly **RELEASE** the plunger to draw sample up into the pipette tip.
5. **DELIVER** the sample by slowly pressing the plunger to the first stop. Depress the plunger to the second stop to expel any remaining sample. **DO NOT RELEASE** the plunger until the tip is out of the sample container.
6. **DISCARD** the tip by pressing the ejector button. Use a new tip for the next sample.

Name: ___________________________

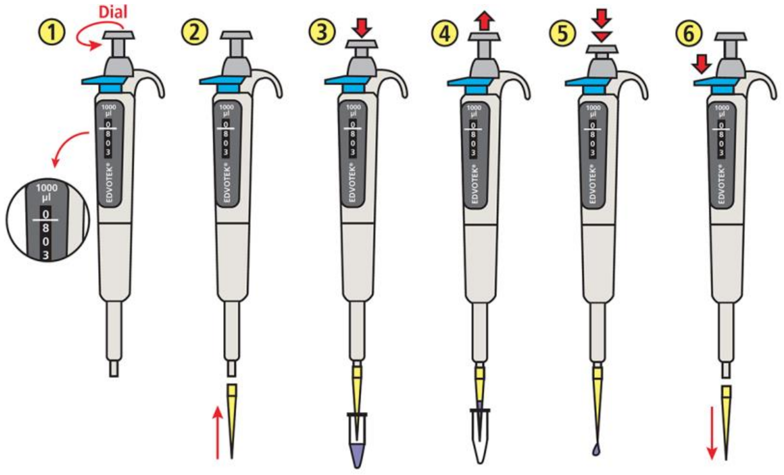


**Protocol Day 1:**

1. Prepare two aliquots (two tubes) of 70µL sterile DI water + 20µL 5x KCM in a 1.7mL snap top tube on ice. Label your tubes “T” for targeting, and “NT” for non-targeting. To “T”, add 1µL pBE-t  and to “NT” add 1µL pBE-nt. Rest your tubes on ice for ~2 minutes.

2. Treat the “GFP-itis" cells by adding a 100µL aliquot of “GFP-itis” bacterial cells to each pBE solution. Incubate on ice for an additional 2 minutes.

3. In a 42C water bath, heat shock your tubes for 75 seconds by letting them sit in the bath with the bottom of the tube where the cells are submerged. Return to ice immediately and rest for 2 minutes

4. Add 800µL of sterile bacterial media to your transformation tubes for recovery. Let the sample “Recover”  for a minimum of 30min, by placing the tubes in a 37C shaker. Label half your agar plate “T” and the other half “NT” and allow them to pre-warm at 37C. You may also wish to add your/your group initials to the plate to help identify it tomorrow.

5. After recovery, add 25µL of the transformed culture onto its appropriate half of the labeled agar plate containing maintenance antibiotics and spread out using a spreader loop. Allow the colonies to grow at 37C for 24hrs.

Name: ___________________________

### Day 2 Questions to Consider:

1. Why do the agar plates contain antibiotic? What might happen if the plate preparers forgot to add the antibiotic?

2. All good scientists include controls in their experiments. What are the controls being used in this exercise? What do they help demonstrate?

3. Tomorrow, you will get a chance to ask some graduate students working in the field of base editing questions about the genome editing field, their personal science projects, their career paths or whatever else you'd like. use this space to jot down a question or two for the panel!

**Protocol Day 2:**

6. Image the plates with the transilluminator and compare GFP output in targeting (pBE-t) and non-targeting samples (pBE-nt).
